# Koopman Analysis Reveals Collective Modes Governing Cellular Dynamics

**DOI:** 10.64898/2026.11.18.767672

**Authors:** Swapnil Dutta, Wenqing Yu, Zhiqian Zheng, Amitava Giri, Jianhua Xing

## Abstract

Living systems are complex nonlinear systems that operate far from thermodynamic equilibrium, and this nonlinearity makes their dynamics difficult to analyze. Several linearization techniques have been developed to address this problem, but most are local. Koopman operator theory is a powerful alternative that linearizes a system globally. Here we show that it not only achieves global linearization but also reveals collective modes that govern the dynamics of complex, out-of-equilibrium non-linear systems. Applying the approach to whole-cell dynamical models, we find that these collective modes have physical and biological significance and relate to distinct processes in the cell cycle and in cell-state transitions. Collective and normal modes are widespread in many-body systems, particularly in condensed matter (e.g., magnons, phonons, hydrodynamic modes) and in macromolecular dynamics (e.g., normal mode analysis). By demonstrating that such modes also exist in cell-state transition dynamics, and in non-equilibrium nonlinear systems more generally, we offer a new theoretical-physics perspective on these systems.

The collective modes separate into slow and fast modes. Long-term dynamics are governed by a few dominant slow modes that form a “slow manifold” of much lower dimension than the original state space. Thus, although the cell is a very high-dimensional system, its cell-cycle dynamics ultimately reduce to a low-dimensional space. Each mode comprises the coordinated behavior of groups of genes and reflects the structure of the underlying gene regulatory network. The modes can also be decomposed into terms that behave like damped harmonic oscillators, drawing an analogy between the modes of a nonlinear system and the normal modes of many-body systems. Finally, our analysis suggests that the dominant modes are closely associated with biological processes in the cell cycle, including mitosis, spindle checkpoint signaling, and DNA replication and repair.

Cells are controlled by thousands of interacting genes, which makes their behavior hard to understand. We show that a mathematical tool from physics, Koopman operator theory, can distill this complexity into a small number of collective patterns of gene activity, analogous to the vibrational modes of molecules and solids. Applied to computational models of the cell cycle, including one built from single-cell data for human lung cancer cells, a few slow modes govern long-term dynamics. Each mode links to distinct processes such as mitosis and DNA replication, and the modes allow genes to be grouped by function. This approach offers a physics-based route from large cellular models to mechanistic insight.

---

Living cells are complex dynamical systems whose behavior emerges from nonlinear interaction among numerous molecular components acting across multiple time and length scales. Cellular functions such as homeostasis, oscillations, growth, and differentiation arise from tightly coupled gene regulatory, metabolic, and signaling networks operating far from thermodynamic equilibrium. While experimental advances, notably single-cell techniques, have enabled increasingly detailed measurements of cellular states, translating this high-dimensional information into mechanistic understanding remains a major challenge.

Mathematical modeling based on dynamical systems theories has been successful in analyzing regulatory networks at small scales (1, 2). Extension of the framework to large-scale networks is an active research area. Specifically, whole-cell models aim to address this problem by integrating diverse molecular processes into unified dynamical descriptions (3–5). However, their high dimensionality and strong nonlinearity complicate analysis and interpretation. As in other many-body systems, the central difficulty is not specifying the microscopic dynamics, but identifying the effective degrees of freedom that govern system-level behavior (6–9).

In condensed matter physics, analogous challenges are addressed through the concept of collective modes, which are emergent dynamical variables that organize the behavior of systems with many interacting constituents. Phonons, spin waves, and hydrodynamic modes provide low-dimensional, predictive descriptions that transcend microscopic details. In the context of macromolecules, normal mode analyses have provided insights into structure-function relations (10–13). This perspective suggests that cellular dynamics, despite their biochemical complexity, may also be governed by a small number of collective modes reflecting coordinated activity across molecular pathways.

Identifying such modes in cellular systems is challenging due to nonlinearity, thermodynamic non-equilibrium driving, and the absence of a clear separation of time scales. Linearization around steady states captures only local behavior, while many data-driven dimensionality reduction techniques lack dynamical interpretability (14). A framework is therefore needed that can extract global, dynamically meaningful structures from complex nonlinear models.

Koopman operator theory and its approximation by dynamic mode decomposition methods have emerged as powerful tools for the data-driven reconstruction of the dynamics of nonlinear dynamical systems by recasting nonlinear dynamics as linear evolution in a space of observables (15, 16). In this work, we instead exploit the spectral decomposition of the Koopman operator to yield eigenfunctions and modes that characterize coherent dynamical patterns, offering a principled definition of collective modes for systems whose governing equations are available but not transparent for mechanistic insight. These modes are globally defined in state space and can be directly connected to underlying biological processes.

Specifically, we apply Koopman analysis to both simple and data-reconstructed cell-cycle models and show that their long-time, high-dimensional dynamics can be approximately described by a small number of dominant collective modes. These modes capture coordinated behavior across a complex gene regulatory network and provide an interpretable, reduced description of cellular dynamics. By explicitly linking large-scale dynamical models to concepts of collective motion from condensed matter physics, our work establishes a new framework for understanding how complex cellular behaviors emerge from underlying biochemical networks.

## Theory

### Koopman operator theory

Many physical and biological systems can be described as dynamical systems whose dynamics are governed by ordinary differential equations (ODEs) of the following generic forms:

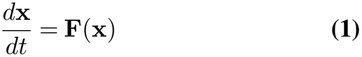

for continuous systems and

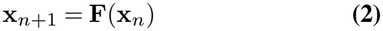

for discrete systems. Here, **x** (**x**_*n*_) ∈ ℳ ⊆ ℝ^*N*^ is generally a multicomponent finite *N* -dimensional vector representing the physical state of the continuous (discrete) system in the state space ℳ, and **F** : ℝ^*N*^ → ℝ^*N*^ ^1^ is a multicomponent vector function acting on the state vectors. For the discrete system, a time interval [0, *T* ] is discretized into *k* time snapshots [*t*_0_ = 0, *t*_1_, *t*_2_, .., *t*_*n*_, .., *t*_*k*_ = *T* ], and **x**_*n*_ is the state vector at the discretized time snapshot *t*_*n*_. For most biological systems, the function *F* is nonlinear in **x** (or **x**_*n*_), which makes the analysis difficult. Note that, experimentally, the observables are often not the physical state **x** (or **x**_*n*_) directly but functions of the physical state of the system. The Koopman operator governs the dynamics of these observables. Here, we present a brief description of Koopman operator theory. For simplicity, we restrict the presentation to continuous systems. A more elaborate, self-contained description is presented in *SI Appendix* (section S1.A), which also includes the case of discrete systems.

For a continuous system, we have a class of Koopman operators *K*^*t*^ governing the dynamics of the observable function *ϕ* through

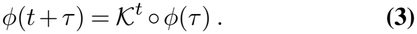

This class of Koopman operators *K*^*t*^ is generated by an infinitesimal generator 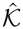 whose action on the observable *ϕ* is given by

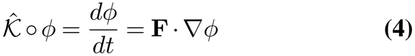

where ∇ is the gradient with respect to the state vector. For more details, see *SI Appendix*, section S1.A.3.

Thus, the Koopman operator lifts the dynamics from the state space ℳ to the observable space ℱ. The Koopman operator is linear, and therefore so are the dynamics in the observable space. However, unlike the space of physical states, which is finite-dimensional, the observable space is generally infinite-dimensional. Thus, as we go from the state space to the observable space, the dynamics go from being generally nonlinear to linear, at the expense of the dimensionality changing from being finite to infinite.

The eigenfunctions *ϕ* of the Koopman operator, also known as Koopman eigenfunctions (KEFs), satisfy

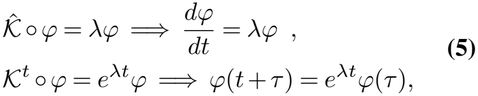

for continuous systems. More details are provided in *SI Appendix*, section S1.A.4.

It is possible to construct a basis set of the observable space using KEFs. In that case, any observable function *ϕ* can be written as a linear combination of KEFs:

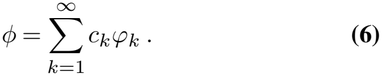

Given that Eq. (5) prescribes how a Koopman eigenfunction evolves with time for a continuous system, the linear combination in Eq. (6) evolves according to

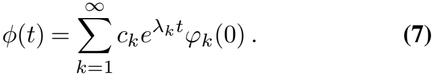

Experimentally, one often measures multiple observables or multicomponent vectors, which can be arranged as a vector function **g** : ℳ →ℂ^*m*^

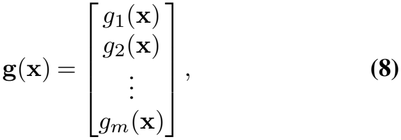

where each *g*_*i*_ : ℳ →ℂ is an observable function and hence, can be expanded as a linear combination of KEFs using Eq. (6) (more details in *SI Appendix*, section S1.A.5) as follows:

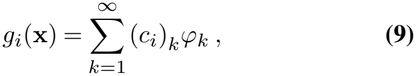

where (*c*_*i*_)_*k*_ can be explicitly determined using dual vectors 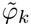 of the KEFs *ϕ*_*k*_ as

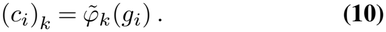

Therefore, we can write

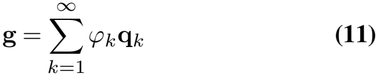

where

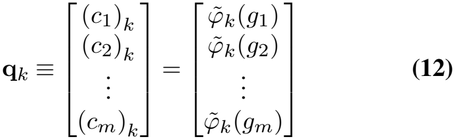

is known as the Koopman mode corresponding to the KEF *ϕ*_*k*_.

In the special case where

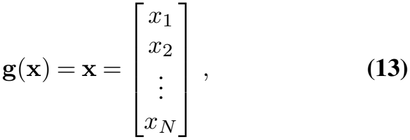

we can describe the evolution of the state vector **x**(*t*) using Eq. (7) and Eq. (12) as

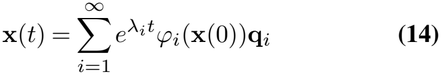

for continuous systems. Thus, the state vector and its evolution can be described as a combination of collective modes. The collection of triplets 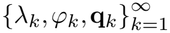 is known as the Koopman mode decomposition of **g**(**x**) (17).

### Contribution from Complex modes

For a pair of complex conjugate modes, their contribution to the Koopman spectral decomposition in Eq. (14) is given by

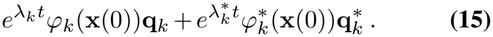

We obtain a central result of this work by rewriting the terms in Eq. (15) as

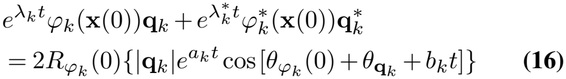

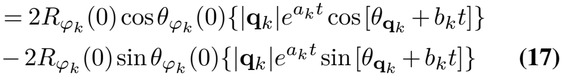

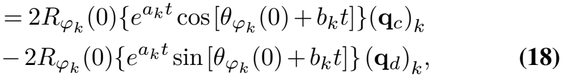

where

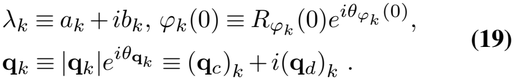

That is, the sum can be split into sine and cosine terms. In Eq. (17), all the initial state information is included in the prefactors 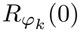 sin 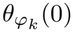 and 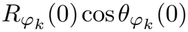, respectively. In Eq. (18), the initial state information is included in both the prefactor 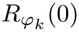 and the time-varying terms in the curly brackets. In each equation, the two terms in the curly brackets differ only by a phase of 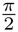, and resemble the dynamics of a damped harmonic oscillator, as one expects from a ‘normal’mode, with the prefactor analogous to the amplitude of the normal mode. However, the crucial difference is that here the underlying dynamics are nonlinear, unlike the harmonic oscillator. That is, Koopman analysis can manifest nonlinear dynamics in analogy to a linear harmonic oscillator. Both |**q**_*k*_| and 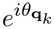 are column vectors whose dimension equals that of a Koopman mode, which in turn equals the dimension of the state space, with

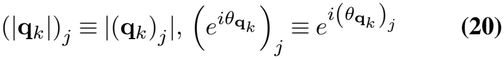

and 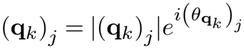.

### Physical interpretation of Koopman spectral decomposition

Eq. (16) can be interpreted physically in two ways, which are discussed below.

#### I. Collective modes

One way is to consider the sine or cosine term in Eq. (17) as a collective time-varying mode, weighted by a prefactor that depends on the initial state. The dynamics of the state-space vector (see Eq. (14)) are then a weighted sum of collective modes. When a Koopman mode **q**_*k*_ is real, the corresponding KEF *ϕ*_*k*_, although generally complex, gives real values when restricted to state-space trajectories^2^, implying that 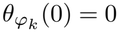. Therefore, the second term (i.e., the sine term) in Eq. (17) vanishes.

Physically and biologically, these collective modes represent the coordinated temporal evolution of different components of the state vectors. For example, in a biochemical network model describing gene transcriptional dynamics, different components of the state vector correspond to the expression of different genes, and hence the overall dynamics of the cell’s transcriptional state can be interpreted as a combination of collective modes of genes, where each collective mode corresponds to coordinated behavior among different genes. These collective modes are the main objects of interest and the crux of this article.

#### II. Coordinate representation

Another interpretation is to use Eq. (18) to regard the Koopman modes **q**_*k*_ as basis-like vectors of the state space and the cosine term 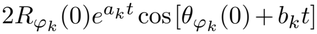 or the sine term 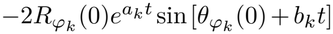 as the coordinates corresponding to those basis-like vectors. When a Koopman mode is real, (**q**_*d*_)_*k*_ = 0, and therefore the second term (i.e., the sine term) vanishes.

These coordinates are time-varying and capture the dynamics of the system. Note that the Koopman modes are not basis vectors in a strict mathematical sense, because there are infinitely many of them in Eq. (14) while the state space is only finite dimensional. Thus this infinite set of Koopman modes is not linearly independent. In fact, each Koopman mode has *N* components, where *N* is the dimension of the state space.

### Extended Dynamic Mode Decomposition

Extended dynamic mode decomposition (EDMD) is an algorithm used to approximate the Koopman operator and Koopman spectrum of a dynamical system (18). We used this algorithm to numerically obtain the Koopman spectrum of a three-variable cell-cycle model and a cell-cycle model reconstructed from scRNA-seq data. A detailed elaboration of this algorithm is presented in *SI Appendix*, section S1.A.6.

### State space classification based on collective modes

Based on the characteristics of the collective modes over long times, the entire state space can be divided into different subspaces:

### Invariant manifold

This subspace is formed from modes with zero or purely imaginary eigenvalues, which correspond to the fixed point(s) and the Fourier modes of the limit cycle(s) of the dynamical system. As *t* → ∞, the evolving state **x**(*t*) eventually falls onto the invariant manifold.

### Damping manifold

This subspace is composed of modes with eigenvalues that have negative real parts and are exponentially damped over time. Starting from an initial state outside the invariant manifold, the system is guided by the dynamics of these modes through the damping manifold and relaxes into the invariant manifold. Therefore, we call these modes *relaxation modes*.

### Unstable manifold

This subspace is composed of modes with eigenvalues that have positive real parts. Therefore, these modes grow exponentially over time. This generally corresponds to regions around unstable fixed points and saddle points, through which the system transits while being repelled from these points.

In the following analyses we focus on individual basins of attraction, each of which can be decomposed into an invariant manifold and a damping manifold. We can write the solution describing the evolution of a dynamical system within such a basin as

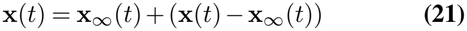

where **x**_∞_ corresponds to the asymptotic part of the solution which lies in the invariant manifold. The remaining part **x**(*t*) − **x**_∞_(*t*) corresponds to the transient part of the solution on the damping manifold. The mathematical formulation of Eq. (21) and the algorithm that we used to separate the relaxation modes from the invariant manifold are provided in detail in *SI Appendix*, section S1.B.

We can rewrite the Koopman spectral decomposition (see Eq. (14)) to explicitly manifest the invariant part **x**_∞_(*t*) and the transient part, in alignment with Eq. (21), as

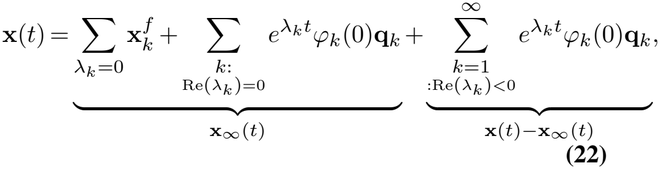

where *λ*_*k*_ is the eigenvalue corresponding to the Koopman eigenfunction *ϕ*_*k*_, and the corresponding Koopman mode is **q**_*k*_. The first two terms on the right-hand side of the equation describe fixed points and limit cycles that belong to the invariant manifold and correspond to **x**_∞_(*t*) in Eq. (21). The remaining term contains the relaxation modes corresponding to **x**(*t*) − **x**_∞_(*t*) in Eq. (21). A summary of the Koopman spectral decomposition, the invariant manifold part **x**_∞_(*t*), and the relaxation part **x**(*t*) − **x**_∞_(*t*) for different cases of asymptotic late-time trajectory behavior is given in Table 1.

**Table 1.**
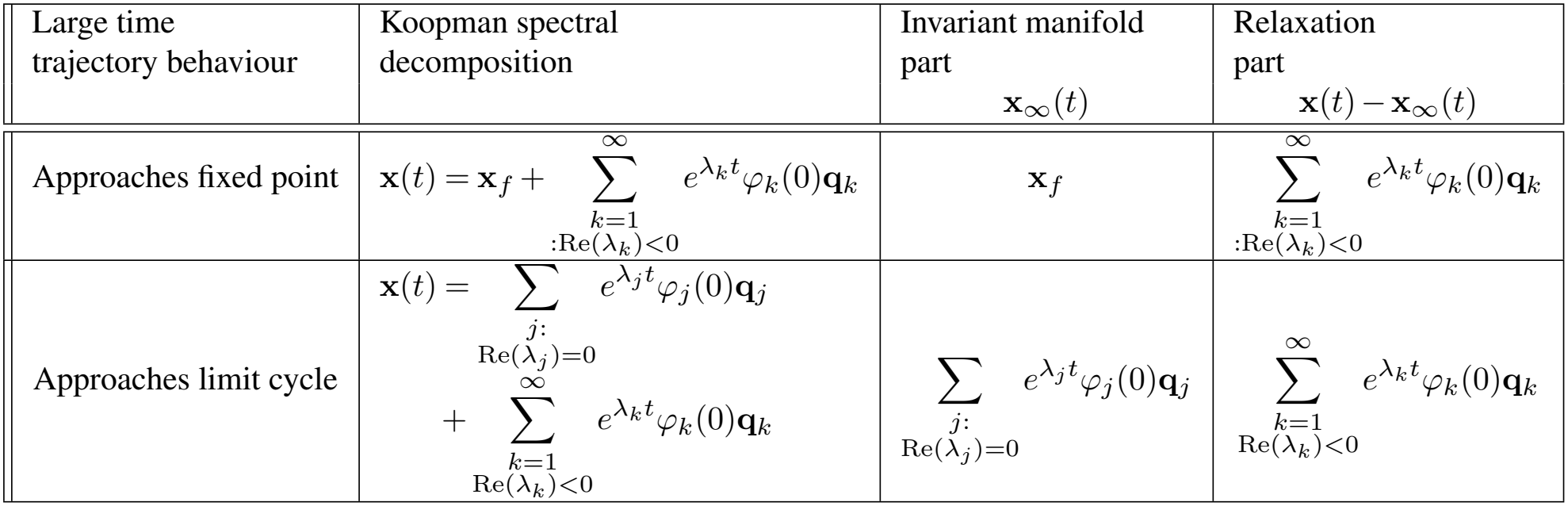
Summary of the invariant manifold (**x**_*∞*_(*t*) from Eq. (21)) and the transient part containing relaxation modes (**x**(*t*) − **x**_*∞*_(*t*) from Eq. (21)), for different cases, assuming continuous systems.

| Large time trajectory behaviour | Koopman spectral decomposition | Invariant manifold part<br>$\mathbf{x}_{\infty}(t)$ | Relaxation part<br>$\mathbf{x}(t)-\mathbf{x}_{\infty}(t)$ |
|---|---|---|---|
| Approaches fixed point | $\mathbf{x}(t)=\mathbf{x}_f+\sum_{\substack{k=1\\:\mathrm{Re}(\lambda_k)<0}}^{\infty} e^{\lambda_k t}\varphi_k(0)\mathbf{q}_k$ | $\mathbf{x}_f$ | $\sum_{\substack{k=1\\:\mathrm{Re}(\lambda_k)<0}}^{\infty} e^{\lambda_k t}\varphi_k(0)\mathbf{q}_k$ |
| Approaches limit cycle | $\mathbf{x}(t)=\sum_{\substack{j:\\\mathrm{Re}(\lambda_j)=0}} e^{\lambda_j t}\varphi_j(0)\mathbf{q}_j$<br>$+\sum_{\substack{k=1\\\mathrm{Re}(\lambda_k)<0}}^{\infty} e^{\lambda_k t}\varphi_k(0)\mathbf{q}_k$ | $\sum_{\substack{j:\\\mathrm{Re}(\lambda_j)=0}} e^{\lambda_j t}\varphi_j(0)\mathbf{q}_j$ | $\sum_{\substack{k=1\\\mathrm{Re}(\lambda_k)<0}}^{\infty} e^{\lambda_k t}\varphi_k(0)\mathbf{q}_k$ |

### Reconstruction of genome-wide dynamical models from scRNA-seq data

With the accumulation of high-throughput single-cell genomics data, efforts have emerged to reconstruct Eq. (1) from the data. For example, assume that a single-cell RNAseq dataset provides the transcriptomic state **x** and its transcription velocity**v** 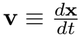 for each of *M* measured cells. Here the vector **x** gives the mRNA concentrations of *N* genes, and **v** is often estimated from snapshot single cell data using biophysical models of mRNA turnover dynamics. Then one learns the function **F**(**x**) by minimizing the overall difference between **F**(**x**) and the measured **v**(**x**) at the sampled cell states. In practice, one can use a function-space expansion with a dictionary {*ψ*}, **F**(**x**) ≈ Σ_*i*_ **c**_*i*_*ψ*_*i*_(**x**, *θ*_*i*_), and the fitting is reduced to determining **c** and *θ* (5). The basis functions can be either pre-specified or parameterized by a neural network. For the models examined in this work, the basis functions {*ψ*} were chosen as sums of one- and two-variable Hill functions.

## Results

### Three-variable Cell-cycle Model

We first applied Koopman analysis to a three-variable model of the cell cycle (19). The three molecular species involved are Cyclin B–associated CDK1, APC*/*C^Cdh1^, and APC*/*C^Cdc20^, with their effective activities denoted by CycB, Cdh1 and Cdc20, respectively. For brevity, we subsequently also refer to the protein species using these notations. Fig. 1A shows the regulatory network diagram. Here, *S* refers to the strength of an external signal, e.g., a growth factor. CycB activates Cdc20 through a switch-like induction, Cdc20 promotes the activation of Cdh1, and active Cdh1 degrades CycB, closing a negative feedback loop that can generate sustained oscillations. In addition, CycB inhibits Cdh1, representing phosphorylation-based inactivation of APC*/*C^Cdh1^. The set of three coupled autonomous ODEs governing the dynamics is given in Fig. 1A.

**Fig. 1.**
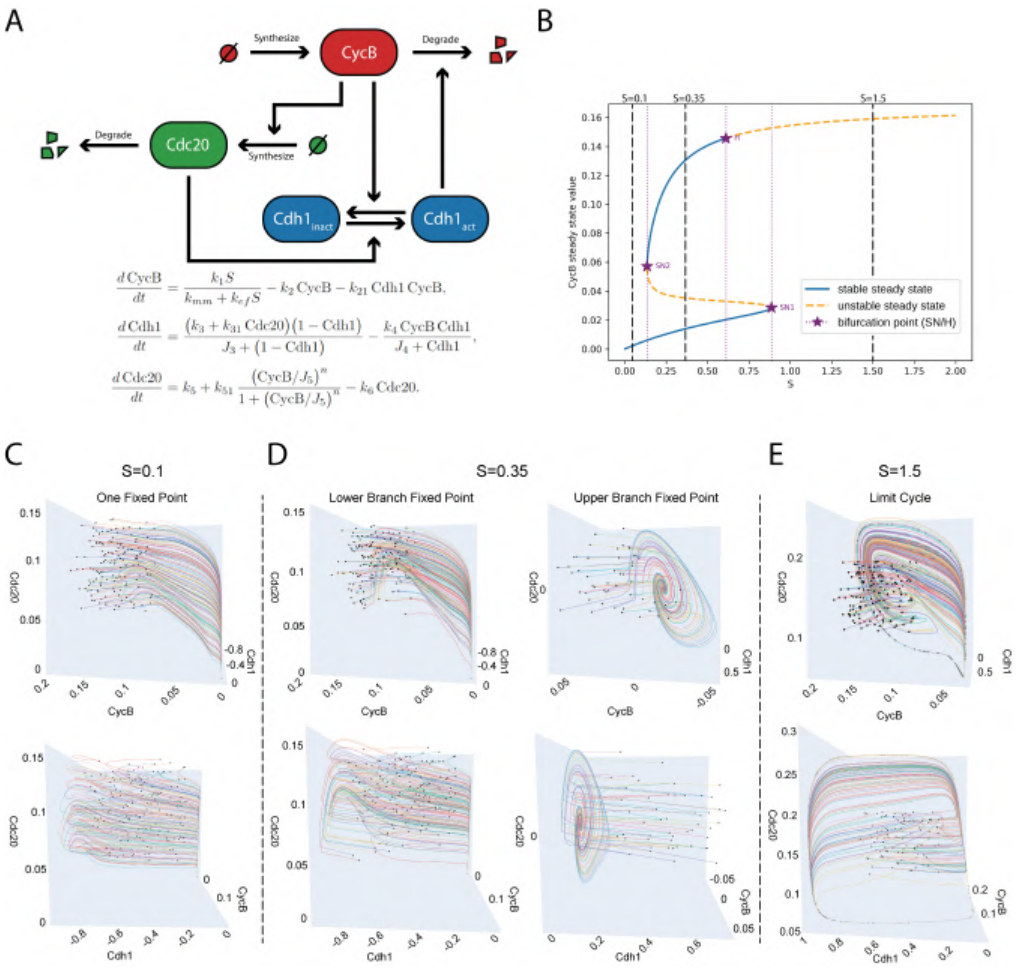
The three-variable cell-cycle model. **A** Regulatory network diagram and the ODEs. **B** Bifurcation diagram with respect to S, showing the steady-state value of `CycB`. **C** Example input trajectories within the basin of attraction around the fixed-point attractor used to train EDMD for *S* = 0.1. Two different views of the same 3D plot are presented here. **D** Same as (C) except *S* = 0.35. Left: the fixed-point attractor (labeled fixed point 0), which lies on the lower branch of the bifurcation diagram, the same branch on which the *S* = 0.1 fixed point lies. Right: the fixed point attractor (labeled fixed point 1) that lies on the upper branch of the bifurcation diagram. **E** Same as (C) except for the basin of attraction around the limit cycle attractor at S=1.5 .

Figure 1B shows a typical bifurcation diagram of the model for a specific set of parameters. Upon varying the signal strength *S*, the model shows three distinct bifurcation regions separated by two saddle-node bifurcation points, SN1 and SN2, and one Hopf bifurcation point H. For *S <* SN1, there is only one stable fixed point. In the range SN1*< S <*SN2, there are two stable fixed points and a saddle fixed point. For *S >*H, the stable fixed point on the top bifurcation branch becomes unstable and a limit cycle emerges around it. Also indicated in Fig. 1B are three representative values of *S* (0.1, 0.35, and 1.5), one from each bifurcation region, for which we performed Koopman analysis. Figures 1C-E show sample trajectories in the state space within the basin of attraction of each stable attractor. These trajectories were used to train EDMD for the Koopman mode decomposition.

We obtained the Koopman eigenvalue spectrum numerically using EDMD at *S* = 0.1, which contains both real and complex values (Fig. 2A). An eigenvalue close to zero corresponds to the fixed point, within numerical error. Both mode selection with a greedy algorithm and direct examination of the mode amplitudes (see *SI Appendix*, section S2.C for more details) reveal that the dominant contributions to the longterm relaxation dynamics of the trajectories come from three complex conjugate pairs of modes whose eigenvalues have the smallest negative real parts (Figs. S1A and B), which are labeled in pairs as modes [1, 2], [3, 4], [5, 6], and shown in red in Fig. 2A. The remaining faster modes, which are shown in blue in Fig. 2A, contribute to the temporal dynamics at early times and then die off rapidly (Fig. S1C). Therefore, we can separate the fast and slow dynamics of the system using the Koopman spectrum.

**Fig. 2.**
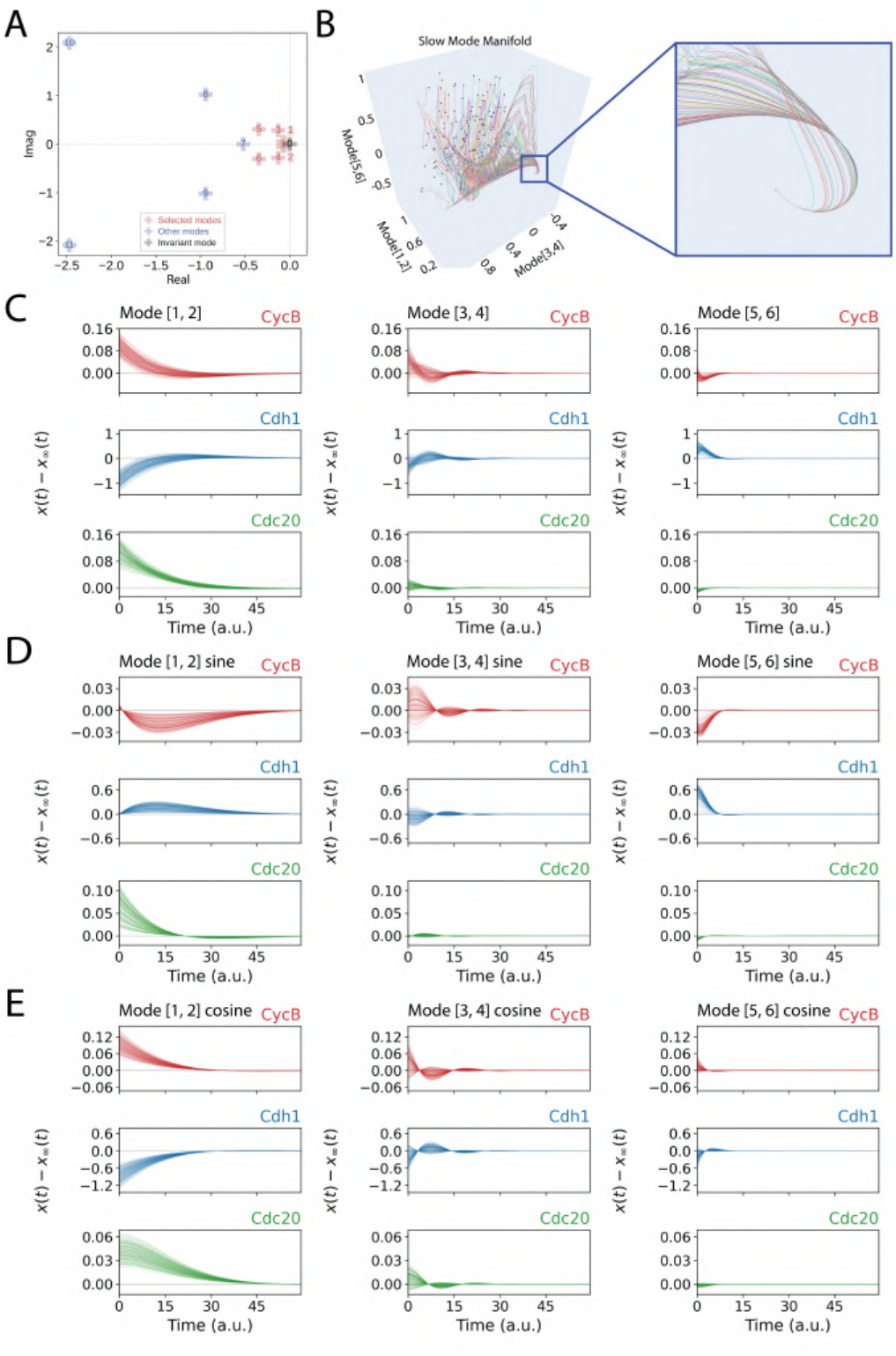
Dynamics and EDMD results showing the three slowest pairs of relaxation modes of the three-variable toy model at *S* = 0.1. **A** Koopman eigenvalue spectrum. The eigenvalues of the three conjugate pairs of the slowest decaying relaxation modes are shown as red crosses. The Koopman mode with zero eigenvalue, corresponding to the fixed point, is shown as a black cross. **B** Trajectories projected onto the space spanned by the cosine parts (see Eq. (18)) of the three pairs of the slowest Koopman relaxation modes. This is the form of coordinate representation in the Koopman mode space. Also shown is a magnified view of trajectories converging to a one-dimensional slow manifold and then to the fixed point. (**C**-**E**) The overall sum (see Eq. (16)) and the sine and cosine terms (see Eq. (17)) of three selected mode pairs [1,2], [3,4], [5,6] of the CycB, Cdh1, and Cdc20 components for selected individual trajectories, after removing the steady-state contribution **x**_*∞*_(*t*), which in this case is the time-invariant fixed point.

### Slow manifold and low dimensional structure

Consider the ensemble of trajectories starting from different initial states. Eq. (S71) suggests that, with the lapse of time, more and more modes die off and the trajectories converge to a manifold formed by the surviving modes whose effective dimensionality in the state space decreases with time. Figure 2B shows such convergence, where the trajectories were plotted in the coordinate representation formed by the cosine parts of the three slowest pairs of Koopman relaxation modes, [1, 2], [3, 4], and [5, 6]. The initial points are scattered throughout the 3D representation. The trajectories first converge to an effective 2D surface as the modes [5, 6] die off, then to a 1D line after modes [3, 4] die off, and finally to the fixed point.

### Biological interpretation of the dominant collective modes

The Koopman modes reflect the coherent temporal evolution of different species in the system. Fig. 2C shows the temporal evolution of the three pairs of dominant relaxation modes (Eq. (16)). For the slowest relaxation mode pair [1, 2], the trajectories of both CycB and Cdc20 start from above-steady-state values and then show damped oscillations. Compared with these two proteins, Cdh1 shows anti-phased temporal dynamics. The temporal evolution of the corresponding sine and cosine terms (see Eq. (17)) of this pair of modes (Figs. 2D and E) behaves like that of damped harmonic oscillators, as expected, showing that these collective dynamical modes can indeed be viewed as analogous to normal modes in many-body physics.

The Koopman mode dynamics reflects how the regulatory network governs the relaxation dynamics. Consider the sine term of mode pair [1, 2]. The Cdc20 level starts with a large deviation above its steady-state value and then decreases because of increased degradation. The above-steady-state Cdc20 level results in an initial decrease of the CycB level because of its inhibition of the latter, and this inhibition decreases with time as the Cdc20 level drops. Meanwhile, the Cdh1 level initially increases from its steady-state value because of Cdc20-to-Cdh1 activation and then relaxes back. That is, this mode involves all three protein species.

The other two pairs show similar but faster damped oscillations (Figs. 2D and E). Mode pair [5, 6] describes mostly fluctuations in the CycB and Cdh1 levels around their steadystate values, while the Cdc20 level remains close to its steady-state value throughout. That is, the mode is localized to the CycB-Cdh1 sub-network. This observation, that high-frequency modes tend to be localized to certain regions of the regulatory network while low-frequency modes are more delocalized, is consistent with what is observed in normal mode analyses of molecular systems (13).

### Evolution of Koopman modes manifests critical slowing– down

At the increased value *S* = 0.35, the relaxation dynamics around each of the fixed points (Figs. S2 and S3) is again dominated by a few Koopman modes but differ notably from that at *S* = 0.1. For fixed point 0, both pairs of the slowest relaxation modes, i.e., the mode pairs [1, 2] and [3, 4], move closer to the origin along the real axis in the eigenvalue plane (Fig. S2A) and consequently show damping dynamics that are more stretched over time (Figs. S2E-G). That is, the trajectories quickly relax into the manifold formed by these two pairs of modes. Such two-phase relaxation is more prominent for fixed point 1 (Fig. S3) and reflects how the underlying vector field evolves when the system approaches the critical points, i.e., SN1 and H (Fig. 1B). For example, upon approaching H along the upper bifurcation branch, the stable fixed point gradually loses its stability, and the magnitude of the vector field **F** within its neighborhood diminishes, until a limit cycle starts to appear after passing H. The spiral-in dynamics within the neighborhood of the fixed point is well described by mode 1, which has a real eigenvalue, and the mode pair [2, 3]. Metaphorically, one can imagine a potential system in which a potential well initially exists but its bottom becomes flatter and flatter as the system moves closer to a critical point; after passing the critical point, the potential takes the shape of a Mexican-hat, with the original stable fixed point at he top of the hat (20).

At *S* = 1.5, the system has passed H and entered a regime of the bifurcation diagram with sustained limit-cycle oscillations (Figs. S4 and S5). The Koopman modes of the limit cycle reduce to its Fourier modes. The remaining Koopman modes describe fast relaxation from an initial state to the limit cycle.

### Koopman analysis of human A549 cell-cycle model reconstructed from scRNA-seq data

Next, we applied Koopman analysis to a cell-cycle model reconstructed from scRNA-seq data of 1,151 human A549 cells. Details of the numerical procedures for the Koopman analysis are given in *SI Appendix* (section S2). The model consists of 399 ODEs describing the mRNA synthesis and degradation dynamics of 399 cell-cycle-dependent genes. The synthesis terms reflect direct and indirect regulation of each gene, involving 11,695 one-to-one regulatory edges and 47 two-to-one regulatory edges modeled by single-variable and two-variable Hill functions, respectively. Given that some genes are regulated only by other genes, we defined a core regulatory network consisting of 65 genes that contribute to non-self-regulation.

We simulated the trajectories of all 399 genes using the 1,151 sampled cell states as the initial states. These trajectories converged to multiple fixed points, which were further grouped into three K-means clusters based on their expression profiles (Fig. 3A and SI Appendix S6). To identify the cell-cycle stages of these fixed-point groups, we examined the expression of genes involved in DNA replication and mitotic regulation. Group 1 shows low to intermediate expression of DNA-replication-related genes such as *CCNE1, CCNE2, CDC6, E2F1, E2F2, PCNA, TYMS, MCM2, MCM6*, and *CCNA2*, together with low expression levels of the mitotic regulators *CCNB1, CCNB2*, and *PLK1*. This expression pattern suggests a state before S-phase entry, potentially a state undergoing G1-phase arrest (21, 22). Group 2 shows the highest expression levels of all ten genes involved in DNA replication and its regulation, while *CCNB1, CCNB2*, and *PLK1* are expressed at lower levels than in group 3. This pattern is consistent with a state that has not passed the S/G2 checkpoint, i.e., a state undergoing S-phase arrest (21, 23). Group 3 shows lower expression levels of DNA-replication-related genes and their regulators than group 2, but the highest expression level of *CCNB1, CCNB2*, and *PLK1*, consistent with an expression profile before passing the G2/M checkpoint, i.e., a state undergoing G2-phase arrest (21)(Fig. S6).

**Fig. 3.**
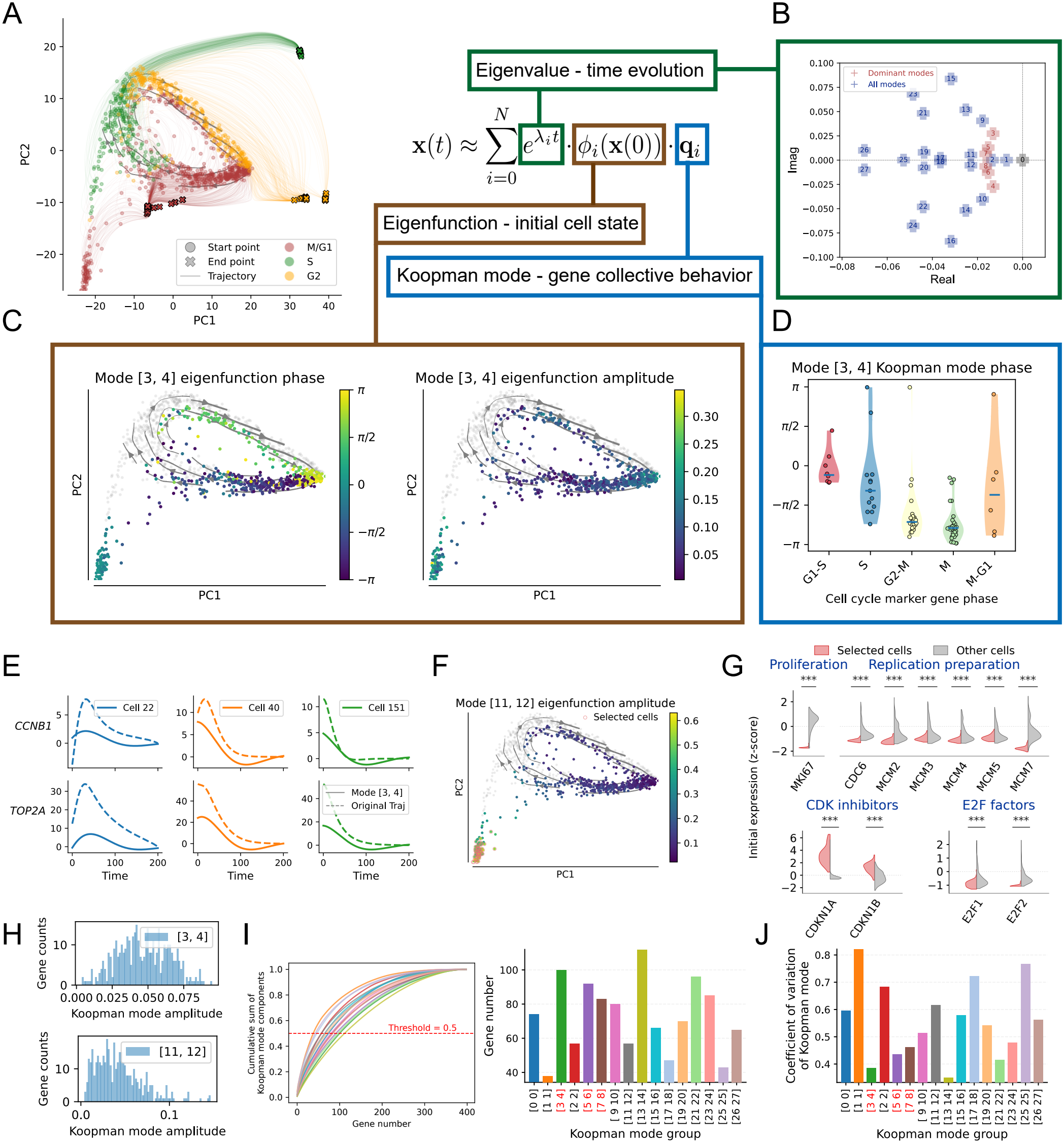
EDMD analysis of the A549 cell-cycle model in the M/G1 basin of attraction. **(A)** Simulated trajectories used to train the EDMD model, projected onto a two-dimensional PCA space. The trajectories converge to three groups of fixed points, indicated by different colors. **(B)** Spectrum of Koopman eigenvalues in the M/G1 basin. The eigenvalue corresponding to the fixed point is indicated by a black dot, the dominant modes by red dots, and the remaining modes by blue dots. **(C)** Eigenfunction phase 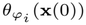 (left) and amplitude 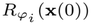 (right) of mode pair [3,4] evaluated at initial cell states in the M/G1 basin and projected onto the two-dimensional PCA space. Cells belonging to other basins are shown in gray. **(D)** Koopman mode phases of cell-cycle marker genes in mode pair [3,4], shown as black dots, with the mean phase for each cell-cycle stage indicated by a blue bar. **(E)** Time evolution of two genes (dashed lines) and the corresponding contributions from mode pair [3,4] (solid lines) in three cells with different initial states. **(F)** Eigenfunction amplitude of mode pair [11,12] projected onto the two-dimensional PCA space. The top 10% of cells with the highest eigenfunction amplitudes are circled in red and referred to as selected cells. **(G)** Expression profiles of selected marker genes in the selected cells identified in (F), compared with those in the remaining cells. The expression patterns suggest that the cells highlighted by mode pair [11,12] are in the G0 state. **(H)** Histograms of normalized Koopman mode amplitudes across genes for mode pairs [3,4] and [11,12]. **(I)** Cumulative fractions of total Koopman mode amplitude with genes ranked in ascending order of their normalized component amplitudes (left), and the minimum number of genes required to account for 50% of the total mode amplitude for each mode (right). **(J)** The coefficients of variation (CV) of the normalized Koopman mode component amplitudes across genes for each mode.

The presence of three clusters of fixed points, rather than a limit cycle, in the transcriptomic ODE model of proliferating A549 cells is consistent with previous mathematical modeling studies (24). The full cell cycle dynamics also include stochastic switching of multiple cell cycle checkpoint programs, largely at the post-translational modification level, which drives transitions between different basins of attraction. Within each of the three major basins, multiple fixed points with only slight expression differences were observed (Fig. 3A). This may arise from the presence of many positive feedback loops in the ODEs, which can generate multiple sub-basins, analogous to the rugged potential surface of a macromolecular system (25).

We applied EDMD separately to the trajectories within each basin of attraction to obtain an approximate Koopman mode decomposition. As in the three-variable model, we first subtracted the corresponding fixed-point state from each trajectory and used the resulting relaxation dynamics to train the EDMD model. Unlike in the three-variable model, the trajectories within the same attractor cluster converge to slightly different fixed points. Consequently, their local relaxation dynamics, and potentially their Koopman decompositions, may differ. However, because these fixed points are closely spaced, we expect the relaxation dynamics around them to share similar dominant features.

The contribution of a collective mode can be decomposed into three parts, corresponding to the Koopman eigenvalue, the Koopman eigenfunction, and the Koopman mode (Fig. 3A). In the following sections, we analyze these components one by one. We mainly focus on the M/G1 basin of attraction in the main text, while results for the other two basins can be found in the SI Appendix.

### Slow manifold and low-dimensional structure of cell dynamics

The Koopman eigenvalues characterize the temporal evolution of each mode. The inverse of the real part of each Koopman eigenvalue defines its characteristic timescale(*τ*_*i*_ = 1*/*|Re(*λ*_*i*_)|). Fig. 3B shows the spectrum of Koopman eigenvalues in the M/G1 basin of attraction. All eigenvalues have negative real parts, except for a zero eigenvalue corresponding to the stationary mode(black point in Fig. 3B), consistent with relaxation toward stable fixed points

Both mode selection with the greedy algorithm (Fig. S7B)and direct examination of the mode amplitudes (Fig. S7C) identified three complex-conjugate mode pairs, modes [3, 4], [5, 6] and [7, 8], that had dominant contributions to the long-term relaxation dynamics. These modes are among the slowest relaxation modes and are highlighted in red in Fig. 3B. Two real modes, [1] and [2], are even slower but contribute less to the long-term relaxation dynamics, possibly involving some slow cellular processes that account for sustained non-genetic cell-cell heterogeneity. The remaining faster modes contribute to the dynamics at early times and then die off relatively rapidly. Consistent with this interpretation, trajectories reconstructed using only the dominant modes deviate from the original trajectories primarily at early times (Figs. S7D and E). Similar results were obtained for the S (Fig. S8) and G2 (Fig. S9) basins, with two and three complex-conjugate mode pairs, respectively, making dominant contributions to the long-term dynamics.

Projection onto a two-dimensional PCA Space (Fig. S10) shows that the dominant modes accurately reconstruct the trajectories near the fixed points, whereas more modes are needed for the early-time dynamics, particularly in the M/G1 basin. When plotted in the 3D representation spanned by the dominant modes of the M/G1 basin, the trajectories are broadly distributed at the beginning and converge slowly to the fixed points (Fig. S11A). In contrast, trajectories in the S and G2 basins quickly fall onto an effective 2D manifold (Figs. S11B and C). These differences are consistent with the greedy selection results (Figs. S7B, S8B, and S9B), which show that the selection frequencies are concentrated in two or three mode pairs for the S and G2 basins, but are spread over more modes in the M/G1 basin. All these results suggest that, compared with the three-variable model, the A549 model also exhibits relaxation toward a low-dimensional slow manifold, but with a broader spectrum of relaxation timescales.

### Biological interpretations of dominant collective modes

A Koopman eigenfunction encodes the dependence of a mode on the initial conditions and is generally complex-valued. A Koopman mode **q**_*i*_ describes the collective behavior of genes and has the same dimension as the gene space. Close examination of the collective modes of five representative cell-cycle marker genes shows dynamics that resemble those of damped harmonic oscillators, again illustrating their analogy to normal modes, as in the three-variable system (Fig. S12).

To illustrate the biological interpretation of these two components, we examined mode pair [3, 4], one of the dominant modes in the M/G1 basin, as an example. We separately plotted the phase 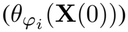 and amplitude 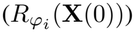 of its Koopman eigenfunction 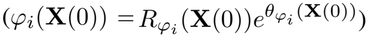 for each initial cell in the 2D PCA space (Fig. 3C). The eigenfunction phase showed a clear correspondence with cell cycle progression (reflected by the gray streamlines in Fig. 3C). Mathematically, such correspondence is expected. Consider a cell state **x**(0) that moves to **x**(*τ* ) as the cell cycle progresses along a streamline, with 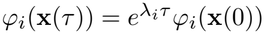. Then for a trajectory starting at **x**^’^ (0) = **x**(*τ* ), we have

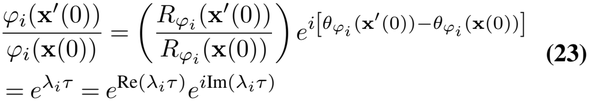

from which we obtain

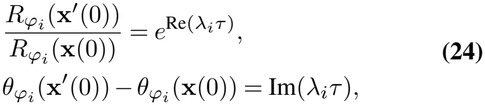

demonstrating that both amplitude scaling and phase shift depend on the time lag *τ* and therefore capture the time-ordering information within the cell cycle.

We next examined the Koopman mode phases of some cell-cycle marker genes (5, 26, 27). For a marker gene indexed by *γ* with its gene expression given by *x*_*γ*_, its Koopman mode phase is given by 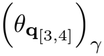 (see the discussion around Eq. (20) for the general definition of 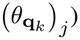. Marker genes associated with the same cell-cycle phase (separately inferred from gene expression profiles by bioinformatics analysis) generally exhibited similar Koopman mode phases, with only a few exceptions in the S and M-G1 cell cycle phases (Fig. 3D). Moreover, the mean Koopman mode phase of the marker genes varies continuously through the cell cycle, decreasing from G1-S to M, and then increasing in M-G1 until it returns close to the initial G1-S value, forming a closed cycle. Thus, the phase organization of mode pair [3,4] is consistent with the temporal ordering of gene expression during cell-cycle progression. Together with the corresponding eigenfunction phase, this finding further supports the close association of mode pair [3,4] with cell-cycle progression. To further support the association between mode pair [3,4] and cell-cycle progression, we examined the trajectories of two genes, *CCNB1* and *TOP2A*, starting from three different initial cells in the M/G1 basin (Fig. 3E). *CCNB1* encodes cyclin B1, which forms a complex with CDK1 to promote mitotic entry and is a key regulator of the G2/M transition (28). *TOP2A* encodes DNA topoisomerase II*α*, whose expression peaks during the G2/M phase (29) and which is essential for maintaining mitotic chromosome structure (30). For both genes, the contributions of mode pair [3,4] closely follow the overall trends of the original trajectories in different cells, demonstrating that mode pair [3,4] captures the time ordering of the initial cell states (Fig. 3E). The pre-ranked gene set enrichment analysis (GSEA), using genes ranked by their normalized amplitudes in mode pair [3,4] (see Methods), also shows enrichment of mitosis-related terms (Fig. S13A).

We next examined mode pair [11, 12], which is slightly faster than the three dominant mode pairs. It has a near-zero imaginary part, indicating negligible oscillatory behavior. Unlike mode pair [3,4], whose KEF amplitude is high across a broad range of cells (Fig. 3C, right panel), mode pair [11,12] exhibits high KEF amplitudes only in a subset of cells outside the cycling manifold (Fig. 3F). We selected the top 10% of cells with the largest eigenfunction amplitudes (circled in red in Fig. 3F) and compared their initial gene expression with that of the remaining cells, which also eventually converge to the G1-arrest fixed points. Compared with the remaining cells, the selected cells had lower expression of the proliferation marker *MKI67* and of several DNA replication genes (Fig. 3G). *MKI67* encodes the proliferation marker Ki67, whose expression is generally low in quiescent cells (31). *CDC6* is required to initiate DNA replication (32). *MCM* genes encode components of the MCM2–7 replication complex, and experiments in A549 cells have shown that reducing their expression can inhibit proliferation (33). Therefore, the lower expression of these genes in the selected cells suggests reduced proliferation. Furthermore, the selected cells had higher levels of *CDKN1A* and *CDKN1B* transcripts. *CDKN1A* and *CDKN1B* encode p21 and p27, respectively. Both proteins can inhibit cyclin–CDK complexes and can thereby inhibit cell cycle progression, so higher expression of their genes indicates growth arrest (34). Additionally, the lower expression of *E2F1* and *E2F2*, which promote the expression of DNA replication genes (35), indicates cell cycle inhibition. Therefore, the expression profiles of these genes suggest that the cells highlighted by mode pair [11, 12] are in the G0 phase.

The corresponding results for the S and G2 basins are presented in *SI Appendix*, sections S3.B.2 and S3.B.3, respectively. Gene Ontology (GO) analysis shows that the two dominant collective modes of the S basin are enriched in mitosis-related terms (Fig. S13B), while, for the G2 basin, two of the three dominant modes are closely associated with DNA-replication-related processes and the other dominant mode is related to mitosis (Fig. S13C).

### Mode localization

We next investigated the degree of localization of Koopman modes in the context of the regulatory network, asking whether a mode is broadly distributed or concentrated in a small subset of genes. We first examined the distributions of the normalized Koopman mode component amplitudes across genes (Figs. 3H and S14A). For some modes, such as mode [3,4], the distributions are relatively broad and approximately symmetric, similar to a normal distribution (Fig. 3H, upper panel), indicating that this mode exhibits a global pattern across genes. In contrast, some other modes, such as mode [11,12], exhibit strongly right-skewed distributions with long tails (Fig. 3H, lower panel), indicating that large mode amplitudes are concentrated in a relatively small subset of genes.

To further quantify this property, we ranked genes in descending order of their normalized Koopman mode component amplitudes and determined the minimum number of genes required to account for half of the total mode amplitude. For the three dominant modes [3,4], [5,6] and [7,8] in the M/G1 basin, this number is 100, 92, and 83 genes, respectively, while mode [11,12] requires only 57 genes (Fig. 3I). We also calculated the coefficients of variation (CV) of the normalized Koopman mode component amplitudes across genes (Fig. 3J). The three dominant modes exhibited some of the lowest CV values, whereas mode [11,12], associated with the dynamics of the specific subset of G0 cells, showed a relatively high CV. Together, these results suggest that the dominant modes in the M/G1 basin tend to be both slow and global. However, this relationship does not always hold in the other two basins (Fig. S14), suggesting that measures other than the number of genes, e.g., intra- and inter-community crosstalk (8), should be examined to investigate the network localization of Koopman modes.

### Collective behavior of genes and network coarse-graining

Even when restricted to only the 65 core genes, the complex network remains complex (Fig. 4A) and is not transparent enough for dissecting mechanistic insights into cell-cycle regulation. Therefore, we set out to coarse-grain the network by grouping genes into clusters. Gene clustering is commonly based on the expression matrix, which describes the expression levels of genes across cells. However, similarity in gene expression across different cells does not necessarily imply related functions. We hypothesized that dynamic information, which emerges directly from interactions within the large and complex regulatory network underlying gene expression, may provide a more informative basis for gene clustering.

**Fig. 4.**
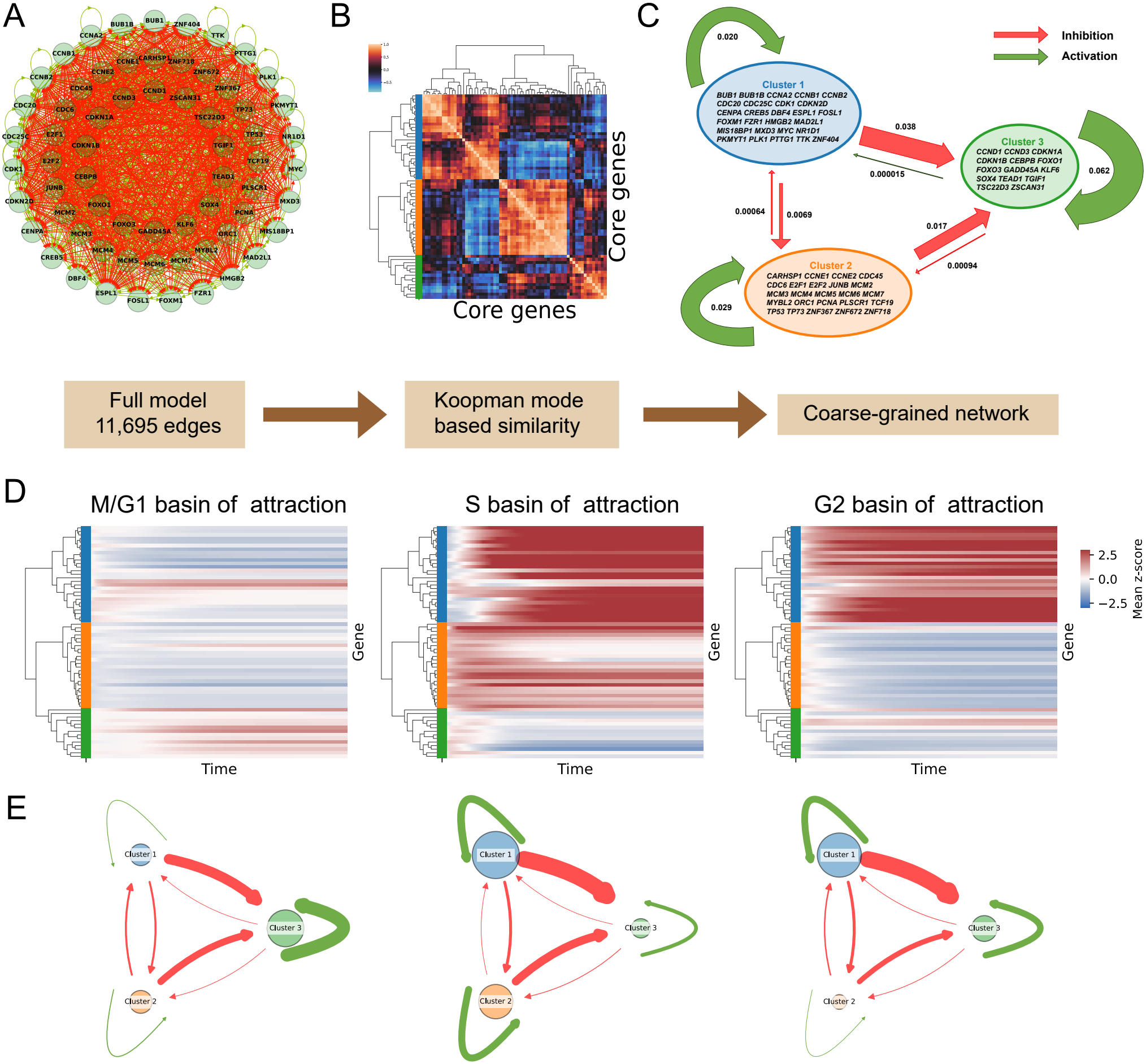
Network coarse-graining based on Koopman modes. **(A)** Gene regulatory network of the 65 core genes. **(B)** Koopman mode-based similarity matrix (see Methods) and hierarchical clustering of the core genes, revealing a three-cluster structure. **(C)** Coarse-grained network showing the three gene clusters and the regulatory interactions between them. **(D)** Heatmap showing the mean expression profiles of the 65 core genes across trajectories converging to each fixed point group. **(E)** Coarse-grained gene regulatory network. Node sizes represent the mean expression levels of genes within each cluster at the fixed-point, while edge widths represent the average interaction strengths between clusters (see Methods).

Noting the low-dimensional structure of cell dynamics, we first performed clustering using only the dominant modes that constitute the slow manifold. For each gene *i*, we constructed a vector **V**_*i*_ containing its components across the dominant Koopman mode pairs from all three basins of attraction:

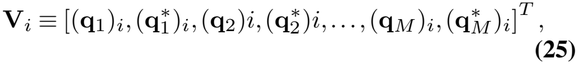

where **q**_*k*_ denotes the *k*-th dominant Koopman mode and *M* is the total number of dominant complex-conjugate mode pairs across the three basins. Both modes in the conjugate pair were retained, yielding a 2*M* × *N* matrix after stacking all the *N* column vectors **V**_*i*_ next to each other, where *N* is the number of genes. We then calculated the pairwise normalized Hermitian inner products between the constructed Koopman mode vectors of the 65 core genes (see Methods in *SI Appendix*, section S2.F). Because both modes in a conjugate pair were retained, the Hermitian inner products are real-valued. Furthermore, normalization makes the similarity measure insensitive to differences in gene expression levels and amplitudes of the Koopman modes. Next, through hierarchical clustering of this similarity matrix, we identified three gene clusters (Figs. 4B and S15A).

We also performed clustering using all Koopman modes. The resulting clusters differed from those based on the dominant modes by only two genes (Fig. S15B), further indicating that the few dominant modes capture most of the relevant dynamical information of the cell system. We further examined two other clustering methods based on the expression matrix and flattened trajectories (see Methods in *SI Appendix*, section S2.F). The trajectory-based clustering (Fig. S15C) agrees well with the Koopman mode-based clustering, reflecting that both methods incorporate system dynamics. In contrast, the expression matrix-based clustering shows a larger difference (Fig. S15D) and does not reflect well the functional and dynamical correlations between genes. For example, the two members of the D-type cyclin family, *CCND3* and *CCND1*, are in two different matrix-based clusters, but they are in the same koopman-mode-based cluster.

To quantify the similarity between clustering results, we calculated pairwise Spearman correlations among the four clustering methods (Fig. S15E). The resulting correlation matrix agrees with our observation that the three methods utilizing dynamical information (dominant Koopman mode-based, all Koopman mode-based, and trajectory-based clustering) show greater similarity to one another than to the expression-based clustering.

To further characterize the biological functions of the three cluster given by the dominant mode-based clustering, we performed GO analysis (Fig. S16A) and examined their gene content. The results showed that the three clusters primarily comprised genes involved in DNA replication, mitosis, and G1 checkpoint signaling, respectively. We then calculated the mean regulatory weight between each pair of clusters to construct a coarse-grained regulatory network of the three clusters (see Methods). The resulting network showed strong self-activation within each cluster and mutual inhibition between different clusters (Fig. 4C). The network structure resembles an expanded toggle-switch motif and gives rise to the three groups of fixed points (Figs. 4D, E and S16B). In the M/G1 basin, self-activation of gene cluster 3 promotes its transcriptional activity, while the other two clusters remain at low activity levels. In the S basin, the activity of cluster 3 is suppressed by inhibition from cluster 1 and 2, while the relatively weak mutual inhibition between clusters 1 and 2,together with their own self-activation, allows them to be activated simultaneously. In contrast, in the G2 basin, only cluster 1 exhibits high activity. Together, these results suggest that Koopman mode-based coarse-graining can reduce a complex gene regulatory network to a simple regulatory motif, in which three self-activation loops and their mutual inhibition provide a mechanistic explanation for the existence of distinct basins of attraction.

## Discussion

Within a cell, a tremendous number of chemical species interact with each other, and numerous biochemical reactions take place at any given time. One research strategy that has long been adopted in cell biology is to investigate individual species and individual interactions. Even with advances in various single-cell techniques, such a reductionist approach has inherent limitations. A different strategy widely used in statistical physics and condensed matter physics focuses on the emergent, coordinated dynamics of different components in a dynamical system. After some transient time, one can often describe the dynamics of such a system with a small number of collective modes.

Koopman analysis and the related EDMD analysis provide a theoretical and computational framework for identifying collective modes from detailed dynamical equations. A Koopman mode manifests the coherent dynamics of multiple genes in that mode, reflecting the underlying gene regulatory relations. The relative phase difference between different genes in a given collective mode remains constant throughout the cell cycle dynamics.

The Koopman modes also provide a procedure for coarse-graining the underlying dynamical model. Genes can be grouped into clusters by analyzing their relative phase differences over the dominant collective modes. As demonstrated with the A549 cell cycle model, genes grouped into the same cluster have relatively small phase differences between them and hence have similar dynamics and behave similarly in the cell cycle. Thus, in the context of cell dynamics, different genes, which together form a high dimensional gene space, can be grouped into a few clusters, with genes in each cluster changing their expression collectively. This simplifies the cell state dynamics within the cell cycle, an therefore helps in understanding and analyzing the dynamics.

Different collective modes can be associated with different cellular processes. For the human A549 cell-cycle model, the dominant collective modes are associated with biological processes such as mitotic cell cycle progression, spindle checkpoint signaling, and DNA replication and repairing. Therefore, the cell cycle can be viewed as a combination of different dynamic modes that have different time scales and involve coherent motion of different groups of genes, and are closely related to different cell cycle-dependent processes.

Different modes may involve different “effective” numbers of gene clusters, or communities, in a regulatory network. We observe examples implying that fast modes tend to be more localized in the gene regulation network, involving smaller numbers of genes, while slow modes tend to be relatively delocalized across the gene regulation network, involving larger numbers of genes. This observation of localized high-frequency modes and delocalized low-frequency modes is consistent with what is observed in normal mode analyses of molecular systems. However, more systematic analyses are needed to evaluate whether this observation holds generally.

The Koopman decomposition of cell-state dynamics also provides a coordinate representation. This representation is helpful in visualizing how the cell state dynamics, starting from a high dimensional space, relaxes into the slow manifold, which has a much lower dimension, finally falling onto the attractor, which is in the invariant manifold.

Thus, the two different interpretations of the Koopman decomposition provide different, in some sense complementary physical and biological insights to the cellular dynamics. The collective mode representation manifests the cellular dynamics as a linear combination of different collective modes consisting of coherent dynamics of multiple genes-with different modes connected to different types of biological processes occurring within the cell, while the coordinate representation shows how the cell state dynamics eventually boils down to a low dimensional space.

In this work we restricted our analyses to two cell-cycle models, and some questions remain for further study. For example, it is mechanistically interesting to examine how well the collective cell-cycle modes are conserved among different cell types or even through evolution. The Koopman spectrum of the three-variable model varies with a control parameter and shows critical slowing-down. Such analyses can provide insights into how cells respond to perturbations and could thereby guide cell differentiation or reprogramming (7, 8). Given that cells are far from thermodynamic equilibrium and that cellular dynamics are non-Hermitian, one expects rich physics along the transition, e.g. high sensitivity to specific perturbations, leading to a diabolic point where two eigenvalues become zero, or an exceptional point where two eigen-vectors also become linearly dependent (36). Applications of the Koopman analysis to high-dimensional cellular systems also encounter some technical challenges, and it is desirable to test advanced Koopman methods (37–40) in this context. In summary, Koopman analysis can be a powerful tool for studying complex nonlinear nonequilibrium systems, and cellular dynamics in particular. It complements efforts to model cellular dynamics in granular detail by providing mechanistic insight into the coarse-grained, essential ingredients.

## Supporting Information Appendix

### S1: Elaboration on Theory

#### S1.A. Koopman Operator Theory: Theoretical and Computational Framework

There are comprehensive reviews on the Koopman operatory theory, its derivative methods such as dynamic mode decompositions, and applications in the context of data-driven dynamics reconstruction (15, 16, 18). Here we provide a pedagogical self-contained introduction of the relevant theoretical and computational framework.

##### S1.A.1. The system and its dynamics

Many systems in nature dynamically evolve with time, which can be modeled using equations of the form

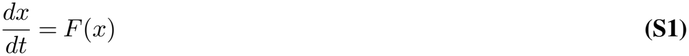

if they evolve continuously with time and

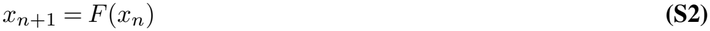

if the time is discretized.

Consider a system whose state space is defined by *N* dimensional real-valued coordinates (*x*_1_, *x*_2_, …, *x*_*N*_ ). In compact notation, the system state is denoted by an *N* -dimensional vector

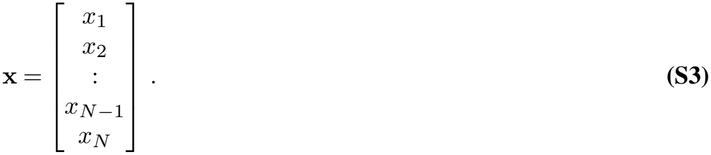

Clearly, **x** ∈ ℝ^*N*^ . Let the statespace (domain), which is a subset of ℝ^*N*^ be denoted by ℳ.

Consider a continuous system. Suppose that we are interested in the dynamics of the system in the time interval 0 ≤ *t* ≤ *T* . Let *I* = [0, *T* ] be the time interval. The evolution of the system in this time interval is given by

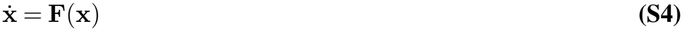

where

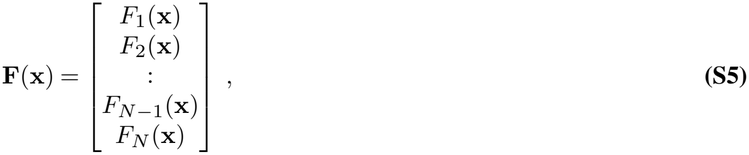

implying **F** : ℝ^*N*^ → ℝ^*N*^ .

As for a discrete system, we are concerned with the time evolution of the system measured at discrete time snaps. Let the time at those discrete time snaps be designated as {*t*_0_, *t*_1_, *t*_2_, .., *t*_*n*−1_, *t*_*n*_} with the corresponding state vectors at those time snapshots being {**x**_0_, **x**_1_, .., **x**_*n*−1_, **x**_*n*_} with **x**(*t*_*i*_) = **x**_*i*_ where *i* ∈ {1, 2, .., *n* − 1, *n*}. In this case, the dynamic is given by

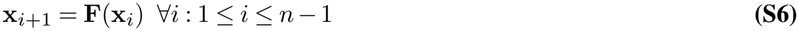

with **F** : ℝ^ℕ^ → ℝ^ℕ^ having *N* components, having the form given in Eq. (S5). Also, each **x**_*i*_ ∈ ℝ^*N*^ as the statespace is *N* dimensional and therefore,

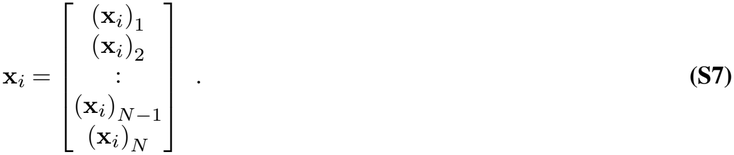

##### S1.A.2. Space of observables

Let us consider the space of all observable functions (also referred to as measurement functions) *ϕ* : ℳ → ℂ that act on the state space ℳ. This space of all observable functions ℱ forms an infinite dimensional linear vector space. In order to define an inner product in this vector space, we assume ℱ = *L*^2^(ℳ, *ρ*) in accordance with (18), where *ρ* is a positive, single valued analytical function with ||*ρ*||_ℳ_ = ∫_ℳ_ *ρ*(**x**)*d***x** = 1. In other words, ℱ is the Lebesgue space of all square integrable functions. This implies that F is also automatically a Hilbert space with the inner product between any two observable functions *ϕ*_1_ and *ϕ*_2_ defined as

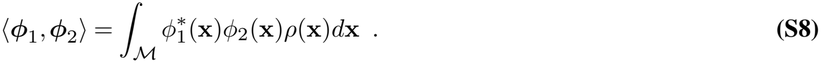

In Eq. (S8) above, we used bold symbol ***ϕ*** to represent *ϕ* as a vector in the observable space ℱ while we use the normal symbol *ϕ* to represent *ϕ* = *ϕ*(**x**) as an observable function *ϕ* : ℳ → ℂ acting on the state space ℳ. From now on, we will keep the following convention. Whenever we talk of inner product (or outer product) in observable space ℱ (and its dual space ℱ ^*^ which will be introduced later), we will use bold font on the observable functions to emphasize them as vectors. Otherwise, we will use normal non-bold font to denote observable functions.

##### S1.A.3. Koopman operator

###### Discrete systems

The Koopman operator *K* : ℱ → ℱ acts on the space of observable functions ℱ. For a discrete system, its action on an observable function *ϕ* ∈ ℱ is given by

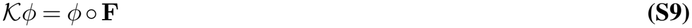

where ◦ means composition of *ϕ* with *F* . So the Koopman operator governs the evolution of the observable *ϕ* in discrete time. Therefore, the function **F** governs the dynamics in state space ℳ while the Koopman operator *K* governs the dynamics in the observable space ℱ. The dynamics defined by **F** and the dynamics defined by *K* can be regarded as two different parameterizations of the same behavior. They are related by the “full state observable” **g**(**x**) = **x** and the Koopman eigenvalues *µ*_*k*_, eigenfunctions *ϕ*_*k*_, and Koopman modes **v**_*k*_ that are required to construct the full state. We will see what they mean in the next few sections. Note that *g*(**x**) = **x** = {*x*_1_, …, *x*_*N*_ }. So, *g* can be split into *N* functions *g*_1_, *g*_2_, .., *g*_*N*_ : *g*_*i*_(**x**) = *x*_*i*_∀*i* ∈ {1, 2, .., *N* }. So, the functions *g*_*i*_(**x**) can be thought of as observable functions in ℱ.

###### Continuous systems

For continuous systems, the dynamics, as mentioned earlier, is governed by Eq. (S4). Let’s define a flow map operator, or time-*t* map F^*t*^ : ℳ → ℳ defined by (17)

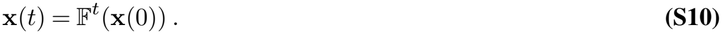

The family of Koopman operators *K* : ℱ → ℱ, parameterized by time *t*, are defined, in terms of their actions on an observable function *ϕ* : ℳ → ℂas (17)

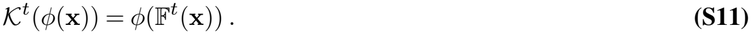

That is, the Koopman operator *K*^*t*^ shifts the observable function *ϕ* forward by time *t*. Therefore, the action of koopman operator *K*^*t*^ on *ϕ* can also be written as

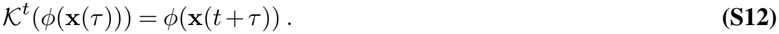

Unlike discrete systems, where we have one Koopman operator with its action defined by Eq. (S9), for continuous systems, we have a family of Koopman operators *K*^*t*^ with their actions defined by Eq. (S11) or equivalently Eq. (S12).These family of Koopman operators are generated by the infinitesemal generator 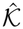 defined as(17)

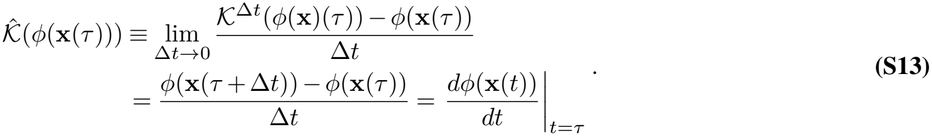

By using Eq. (S4) and the chain rule, we have

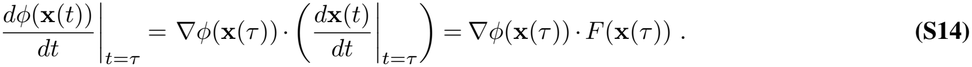

Using Eq. (S13) and Eq. (S14), the action of infinitesimal generator 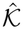 on the observable function *ϕ* can be written as

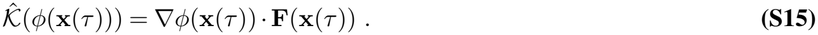

The infinitesimal generator 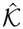 can be thought of as being analogous to the generator of a Lie algebra while the set of Koopman operators *K*^*t*^ can be thought of as being analogous to the elements of the corresponding Lie group generated by the infinitesimal generator. It should be noted that this is just an analogy as in strict mathematical sense, the set of Koopman operators *K*^*t*^ does not always form a group but is always a semigroup^3^.

##### S1.A.4. Koopman eigenvalues and eigenfunctions

###### Discrete systems

For a discrete system with dynamics governed by Eq. (S6) and the Koopman operator defined by Eq. (S9), an observable function *ϕ* ∈ ℱ is a Koopman eigenfunction with a corresponding eigenvalue *µ* if it satisfies the equation

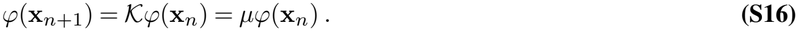

Therefore, the time evolution of a Koopman eigenfunction for a discrete system is

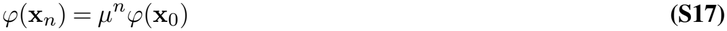

or more generally

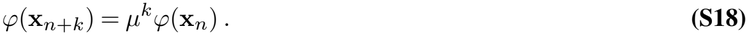

###### Continuous systems

For continuous systems with dynamics governed by Eq. (S4) and the group of Koopman operators defined by Eq. (S12), which are generated by the infinitesimal operator 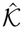 defined by Eq. (S13)(or equivalently Eq. (S15)), an observable function *ϕ*(**x**) is a Koopman eigenfunction of 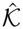 with corresponding eigenvalue *λ* if it satisfies

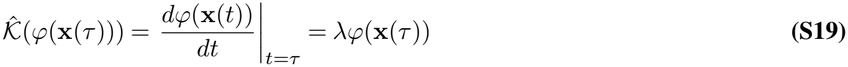

or equivalently

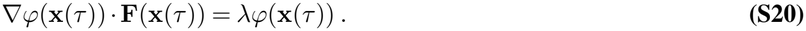

The time evolution of a Koopman eigenfunction of 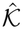 can be obtained by integrating Eq. (S19) from which we obtain

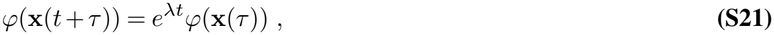

which can also be written as

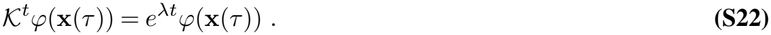

Thus a Koopman eigenfunction of the infinitesimal Koopman generator operator 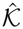 with Koopman eigenvalue *λ* is also an eigenvalue of the family of Koopman operators *K*^*t*^ with a Koopman eigenvalue *e*^*λt*^.

###### Connection between discrete and dynamical systems

A Koopman eigenfunction of a continuous system is also the Koopman eigenfunction of a discrete system and vice versa. The corresponding Koopman eigenvalues are different but related. Let *ϕ*(**x**) be the eigenfuntion of the infinitesimal Koopman generator 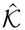 with eigenvalue *λ*. Then, *ϕ*(**x**) satisifes Eq. (S19) and Eq. (S21). Now, let’s consider a discretized version of the same physical system. Let is discretize a time interval [0, *T* ] into discrete time snaps {*t*_0_, *t*_1_, .., *t*_*n*_} with time interval Δ*t* between two successive time snaps. With **x**(*t* = 0) = **x**_0_ and **x**(*t* = *t*_1_) = **x**_1_, we obtain, from Eq. (S21)

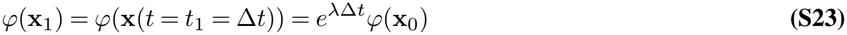

by taking *τ* = 0 and *t* = Δ*t*. Comparing with Eq. (S16), we see that *ϕ* is also a Koopman eigenfunction of the discretized version with eigenvalue *µ* = *e*^*λ*Δ*t*^ implying

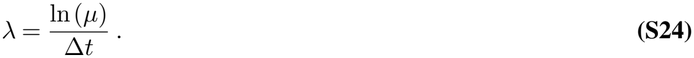

So, Eq. (S24) relates the Koopman eigenvalue of a continuous system and its discretized version. This equation is very important for the Koopman amalysis of our cell-cycle models as we discretize the system while doing numerical analysis and then report the results back the in continuous version .

Also, the complete set of Koopman eigenfunctions can form a complete basis of ℱ. So, if B = {*ϕ*_1_, *ϕ*_2_, ..} is a complete basis of ℱ, any observable function *ϕ* ∈ ℱ can be written as

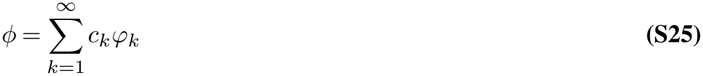

which is true for both discrete and continuous systems. Please note that since ℱ is infinite dimensional, ℬ has infinite elements.

Also, if we know the representation of an observable function *ϕ* at any point of time in terms of a Koopman eigenbasis, we would know how *ϕ* evolves as a function of time. For a continuous system,

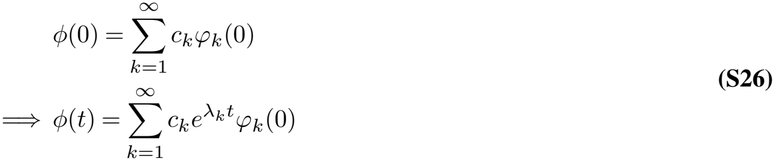

where *ψ*(0) ≡ *ψ*(*t* = 0) for any artitrary *ψ* ∈ ℱ. Also, to go from first line in Eq. (S26) to second line, we have used Eq. (S22). For a discrete system,

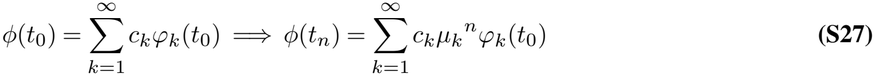

where *t*_*n*_ = *n*Δ*t* is the *n*^th^ discrete time snap.

##### S1.A.5. Koopman modes

In experimental observations, one often measures a set of observable functions instead of single observable function. Let

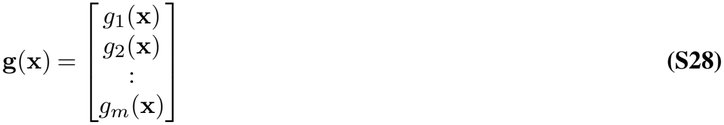

be a set of *m* such observable functions. Then, each *g*_*i*_ can be expanded in a Koopman eigenbasis in accordance with Eq. (S25). The coefficients corresponding to the eigenfunctions in expansion of *g*_*i*_ can be expressed in a generalized way using the concept of *dual vectors* which we shall elaborate shortly.

It is to be noted that the Koopman operator is generally not guaranteed to be normal or self-adjoint, and therefore, it is not guaranteed to produce an orthonormal Koopman eigenbasis. Therefore, in general the completeness relation Σ_*n*_ ***ϕ***_*n*_ ⊗ ***ϕ***_*n*_ = I which holds for an orthonormal basis *does not* generally apply.

Instead, for a complete Koopman eigenbasis ℬ = {*ϕ*_1_, *ϕ*_2_, ..} of ℱ, we have

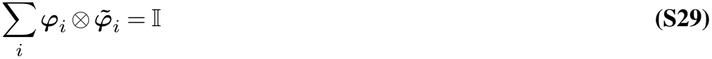

where 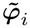 is the dual vector of the Koopman Eigenfunction ***ϕ***_*i*_ and belongs to the dual space ℱ^*^ of the observable space ℱ. The dual space ℱ^*^ consists of linear functions 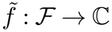, which act on the observables^4^. Also, 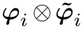 denotes the outer product between 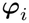 and 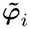. We will briefly elaborate the notion of outer product with a simple example shortly below. Please note that in similar spirit of what is mentioned in the discussion below Eq. (S8), we use bold symbol ***ϕ***_***i***_ to represent the Koopman eigenfunction *ϕ*_*i*_ as a vector in the observable space ℱ while we use the normal symbol *ϕ*_*i*_ to represent *ϕ*_*i*_ = *ϕ*_*i*_(**x**) as an observable function *ϕ* : ℳ → ℂ acting on the state space ℳ.

The inner product between a vector 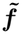 belonging to the dual space ℱ^*^ and a vector ***ϕ*** belonging to the observable space ℱ is defined as

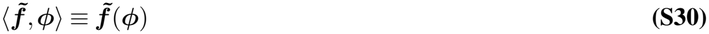

where 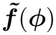 denotes the linear functional 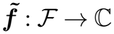 operating on the vector ***ϕ*** ∈ ℱ and the notation 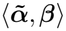 denotes the inner product between the vector 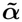 belonging to the dual space and ***β*** of the original vector space.

Also, it should be noted that for the Koopman eigenfunction ***ϕ*** with eigenvalue *λ*, the dual vector 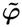 is an eigenvector of the adjoint of the Koopman operator (which is related to the Perron-Frobenious operator) with an eigenvalue *λ*^*^, which is the complex conjugate of *λ*. Mathematically,

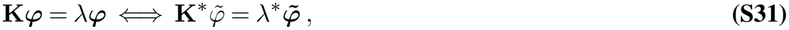

where **K** is used as a general notation of Koopman operator for both discrete and continuous systems and **K**^*^ is its adjoint operator. An alternative interpretation of the dual vector 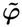 is that it can be thought of as the *left eigenvector* of the Koopman operator satisfying

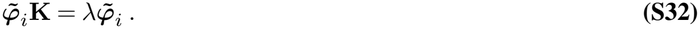

Also, for the complete Koopman eigenbasis ℬ = {*ϕ*_1_, *ϕ*_2_, ..} of ℱ, the set of their corresponding dual vectors 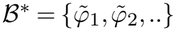 forms a complete eigenbasis of the dual space ℱ ^*^ and these two sets of bases B and B^*^ form a biorthogonal system satisfying

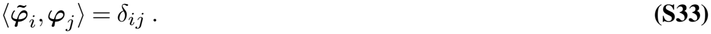

One physically intuitive way of understanding the notion of a vector space and it’s dual space is through the following simple example.

Let us consider the vector space V ≡ ℂ^3^ of 3 × 1 column vectors over the field of complex numbers. Then, it’s dual space V^*^ will be the vector space of all 1 × 3 row vectors over the field of complex numbers. One possible choice of basis for V is

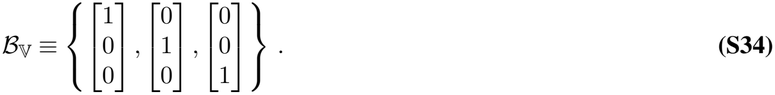

The corresponding dual basis will be

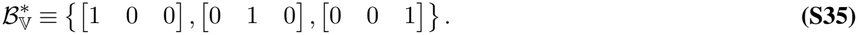

In the general for a vector **w** ∈ V, it’s dual vector 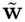 will be it’s complex conjugate. That is,

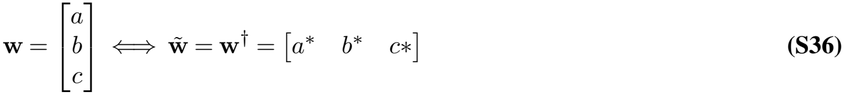

where **w**^†^ denotes the complex conjugate (also known as Hermitian conjugate) of the vector **w**. Also, in general, for an arbitrary vector **w**_1_ ∈ V and dual vector 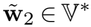, the outer product 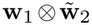 and inner product 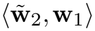 between them are given simply by the matrix multiplication. That is to say,

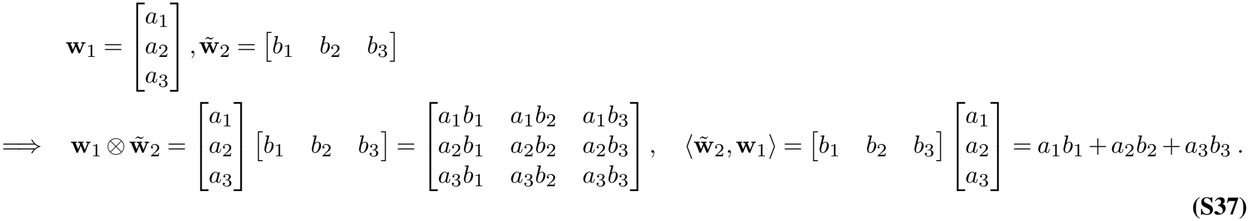

Then, for the basis ℬ _V_ defined in Eq. (S34) and its dual basis B^*^ defined in Eq. (S35), one can easily verify the completeness relation in Eq. (S29) and biorthogonality relation in Eq. (S33).

Now, we get back to expressing *g*_*i*_ in the Koopman eigenbasis by relating the coefficients of expansion with the concept of ‘dual vectors’. Using the general completeness relation Eq. (S29), we have

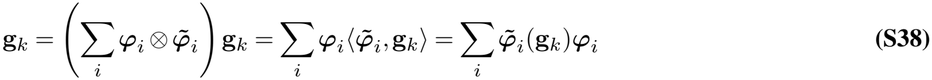

where in the last step, we have used Eq. (S30) and for 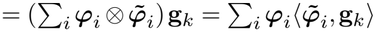, we have used the identity 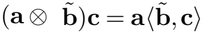. One easy way to verify this identity is through the above example of the V ≡ ℂ^3^ and its dual vector space V^*^. Then, using Eq. (S38), *g*(**x**) in Eq. (S28) can be expanded as

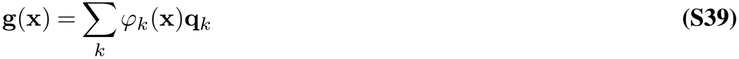

where

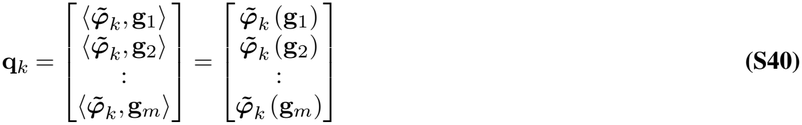

are known as the *Koopman modes* corresponding to the Koopman eigenfunction *ϕ*_*k*_(**x**).

Since we know how the Koopman eigenfunctions evolve for both discrete and continuous cases (see Eq. (S18) and Eq. (S21) respectively), we can use that with Eq. (S39) to determine the time evolution of **g**(**x**). For discrete case, we have

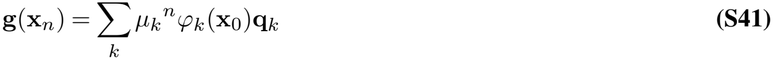

and for continuous case, we have

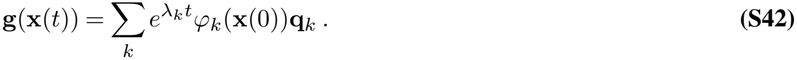

In the special case where **g**(**x**) is the fullstate observable **g**(**x**) = **x**,using Eq. (S41) we have

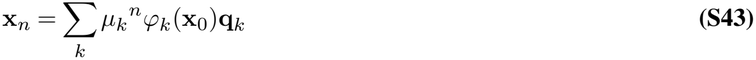

for discrete case, and using Eq. (S42),

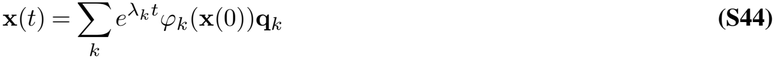

for continuous systems. Here, the Koopman mode **q**_*k*_ is given by

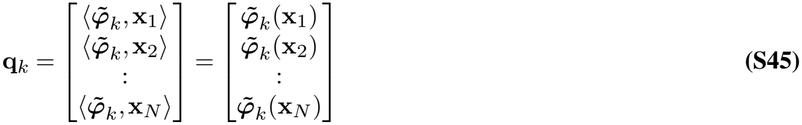

where **x**_*i*_ is the *i*^th^ component of **x**, as shown through Eq. (S3). However, here in Eq. (S45), we are denoting **x**_*i*_ in bold font, unlike in Eq. (S3) to emphasize that **x**_*i*_ : ℳ → ℝ ⊂ ℂ is an observable function and in the above Eq. (S45), is interpreted as a vector in observable space ℱ. As mentioned earlier, we see that the fullstate observable *g*(**x**) = **x** relates the dynamics of the observable space ℱ (which is governed by the Koopman operator and can be expressed in terms of Koopman eigenfunctions) with the dynamics in state space ℳ.

The sequence of triplets 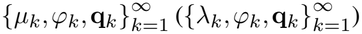 is known as the Koopman mode decomposition of *g*(**x**) for discrete (continuous case) (17).

##### S1.A.6. Extended Dynamic Mode Decomposition (EDMD)

Given a dynamical system defined by Eq. (S1) (for continuous time systems) or Eq. (S2) (for discrete time systems), the Extended Dynamic Mode Decomposition(EDMD) is an algorithm to approximate the Koopman Operator, and hence, subsequently, the Koopman spectrum, which constitutes the Koopman Eigen-values, Koopman Eigenfunctions and Koopman Modes. Since in practice, a dynamical system is often numerically studied generally by discretizing the system and expressing its dynamics as a discrete time series data, below we confine our discussion of EDMD to discrete systems only.

###### Pre-requisites for the EDMD algorithm

###### Discrete time series data set

We train an EDMD algorithm using a time series data set capturing the dynamics a system in the form of *P* snapshot pairs of the system 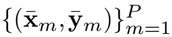 taken at some interval, i.e., 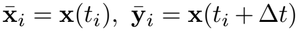 with 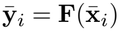, where **F** governs the dynamics according to Eq. (S2) and *t*_*i*_ ∈ {*t*_1_, *t*_2_, .., *t*_*P*_ }. Here, the time interval of interest is discretized into *P* time snapshots *t*_1_, *t*_2_, .., *t*_*P*_ . Note that the time interval between these consecutive time snaps need not necessarily be uniform. This time series data set is a prerequisite for the EDMD.

###### Dictionary of generic basis functions

Another prerequisite is the need for a dictionary *D* of generic basis functions _1_, *ψ*_2_, .., *ψ*_*K*_. That is,

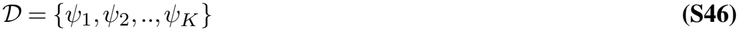

is a set of *K* generic basis functions. In the context of EDMD, these basis functions are observable functions and hence, belong to the infinite-dimensional observable space ℱ. These basis functions need to be linearly independent. Therefore, the linear span ℱ _*D*_ of the basis functions in dictionary *D* forms a *K* dimensional subspace of ℱ.

###### Approximating the Koopman operator

Since ℱ_*D*_ is a finite-dimensional subspace of ℱ, ℱ_*D*_ is generally not invariant under the Koopman operator. In other words, the action of Koopman operator *K* on an observative function *ϕ* ∈ ℱ _*D*_ may map the function out of ℱ _*D*_. The action of the Koopman operator *K* on an observable *ϕ* ∈ ℱ _*D*_ can be mathematically expressed as

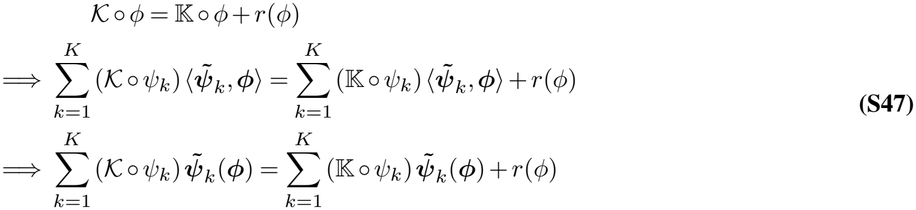

where the second line of Eq. (S47) follows from the generalized completeness relation Eq. (S29) for any arbitrary basis, and the third line follows from Eq. (S30), with 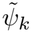 being the dual vector of *ψ*_*k*_. Also, K is a *K* × *K* matrix which is an approximation of the Koopman operator *K* confined to operating on ℱ_*D*_^5^. Here, *r*(*ϕ*) is the residual part that lies outside of ℱ_*D*_. To be more precise, *r*(*ϕ*) lies in ℱ − ℱ _*D*_. In order to find the best approximation of K, we try to minimize the residual part *r*(*ϕ*) for any general *ϕ* ∈ ℱ _*D*_. The residual part *r*(*ϕ*) in general is a function that acts on the statespace ℳ and, therefore, can be trained on the time series data set mentioned earlier to minimize itself. We define a quantity

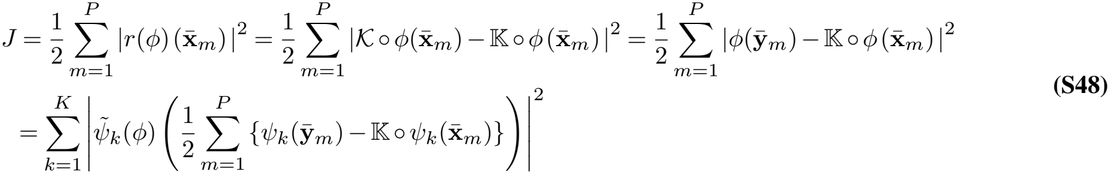

which incorporates the residue on the training dataset. We seek to minimize it. Please note that *ϕ* ∈ ℱ_*D*_ is arbitrary and therefore, in Eq. (S48), *J* has to be minimized for any arbitrary set of *K* 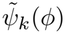s in general. This looks like a least-square problem and we have to find K that minimizes it. Being a least square problem, there cannot be discrete isolated local minima, but a unique global minimum or a continuous family of minima (18). In the later case of continuous family of minima, one requires some kind of regularization, which in this case is truncated singular value decomposition, to ensure that the minimizing solution of K is unique. The K that minimizes *J* in Eq. (S48) is (18)

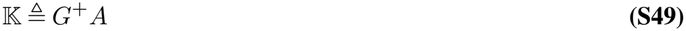

where

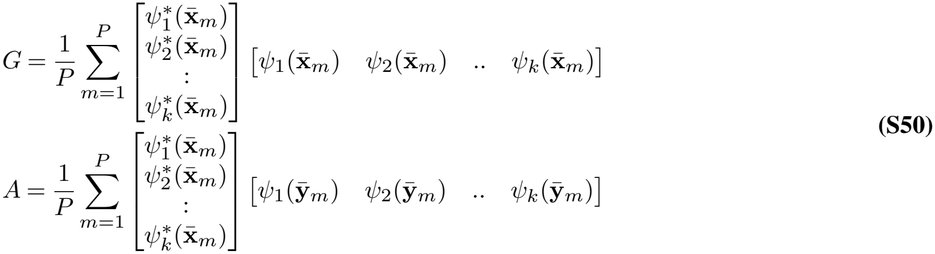

and *G*^+^ is the pseudoinverse (also known as Moore-Penrose inverse) of *G*. Please note that both *G* and *A* are *K* × *K* matrices. So, K : ℱ _*D*_ → ℱ_*D*_ is a finite dimensional approximation of *K* restricted to ℱ_*D*_ that maps any arbitrary *ϕ* ∈ ℱ_D_ to another 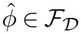 by minimizing the residue at data points.

###### The approximate Koopman spectrum

Once we have found K from Eq. (S49), which is a finite dimensional approximation of *K*, using the EDMD algorithm, we can find the eigenspectrum of K to obtain the finite dimensional approximation of the Koopman spectrum.

###### Approximate Koopman eigenvalues and eigenfunctions

Please note that since we are working in the dictionary basis *D* = {*ψ*_1_, *ψ*_2_, .., *ψ*_*K*_}, the operator K is expressed in that basis. Therefore, once we diagonalize the matrix K and find its eigenvalues by finding the roots of the characteristic polynomial of K and subsequently find the eigenvectors of K, these eigenvectors are expressed in the dictionary basis. So, if one of the eigenvectors of K, corresponding to the eigenvalue *µ*_*j*_, is expressed as the column vector 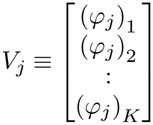, what these elements (*ϕ*_*j*_)_1_, (*ϕ*_*j*_)_2_, .., (*ϕ*_*j*_) represent are actually the coefficient of the eigenvector *ϕ*_*j*_ in the dictionary basis. That is to say,

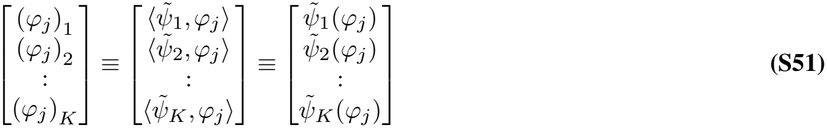

and therefore,

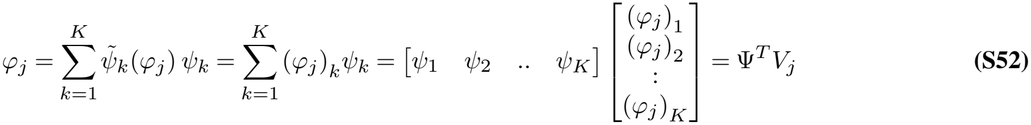

where

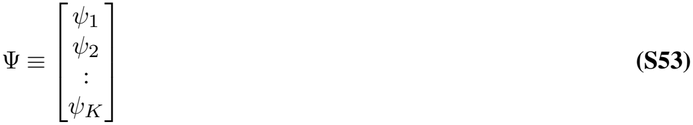

is the generic basis functions of the dictionary *D* expressed in an array as a column vector.

These eigenvalues and eigenvectors are finite dimensional approximations of the Koopman eigenvalues and Koopman eigen-functions respectively.

###### Approximation of Koopman modes

Now, let’s find an approximation to the Koopman modes. Let’s consider the full-state observable

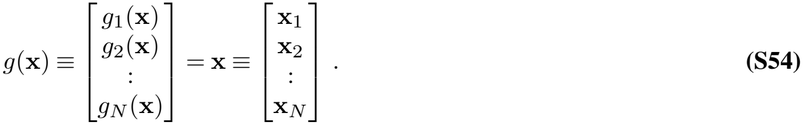

Then, when restricted to the span of the dictionary basis ℱ_*D*_, the observable function *g*_*j*_(**x**) can be expressed as

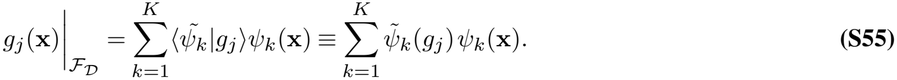

which follows from the completeness relation Eq. (S29) when restricted to ℱ_*D*_ in the dictionary basis. Also, we use the version of Eq. (S33) in ℱ_*D*_ to relate the inner product 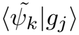to 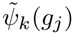. Here, *g*_*j*_(**x**)|_ℱ*D*_ denotes the observable *g*_*j*_(**x**) restricted to ℱ_*D*_. From now on, in this subsection, it will be implicitly understood that *g*_*j*_(**x**) is restricted to ℱ_*D*_, and we will drop the | _ℱ*D*_ to denote *g*_*j*_(**x**)|_ℱ*D*_ simply as *g*_*j*_(**x**). Eq. (S55) can be written in compactified form as

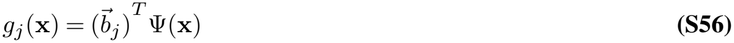

where

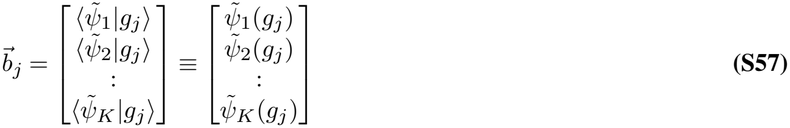

and Ψ(**x**) follows from Eq. (S53).

Having expressed *g*_*j*_(**x**) in the dictionary basis, we can now express the fullstate observable *g*(**x**) as

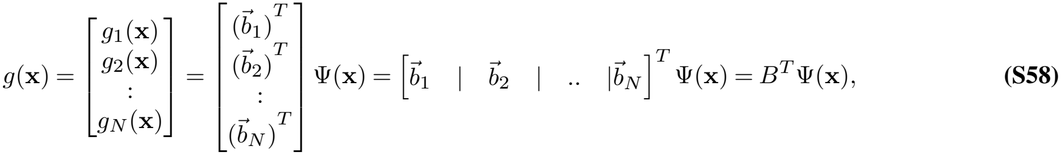

where

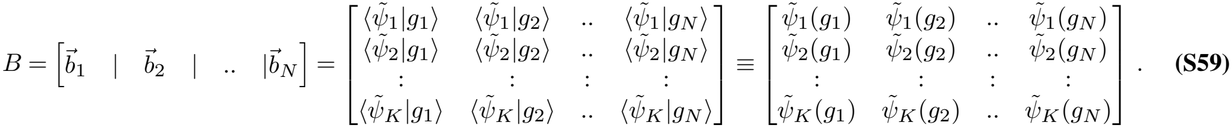

Also, the fullstate observable can be written in terms of “approximate” Koopman modes as

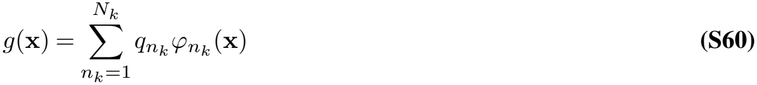

where 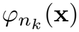 are Koopman eigenfunctions of K acting as approximation to actual Koopman eigenfunctions when restricted to ℱ_*D*_, while *q*_*k*_ are the corresponding “approximate” Koopman modes. Let

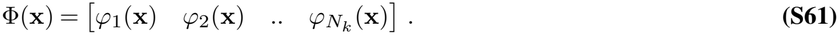

Then Φ(**x**) can be expressed in dictionary basis ℬ as

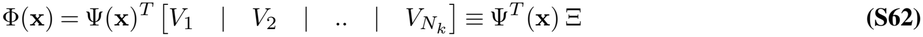

where Ψ(**x**) follows from Eq. (S53) and

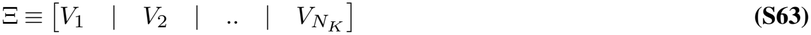

where *V*_*j*_ is an eigenvector of K in the dictionary basis. Therefore, Ξ contains all the eigenvectors of K in dictionary basis stacked next to each other. Then, Eq. (S60) can be written as

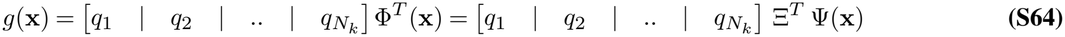

where 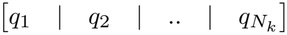 is a *N* × *N*_*K*_ matrix formed by stacking the *N*_*K*_ Koopman modes 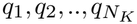, each of which is a *N* × 1 column vector, next to each other. Also, by taking transpose of Eq. (S62), we have Φ^*T*^ (**x**) =Ξ ^*T*^ Ψ(**x**).

Comparing and equating the two different ways in which the full state observable is expressed in Eq. (S58) and Eq. (S64), we have

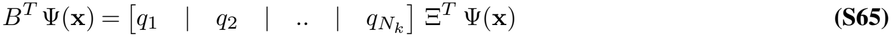

from which we get

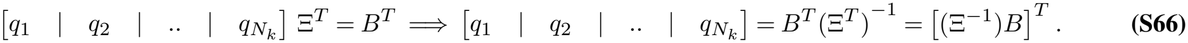

Therefore, if we find the inverse of the matrix Ξ, we can determine the matrix 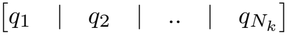 and therefore, the approximate Koopman modes *q*_1_, *q*_2_, .., *q*_*N*_ . In order to find the inverse of Ξ, we note the following. Let 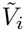 be the left eigenvector of the matrix K with eigenvalue *µ*_*i*_ in the dictionary basis. Please note that in this case, 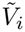 is a 1 × *K* row vector. Then, we have

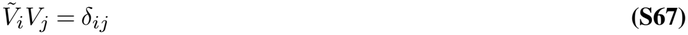

where *V*_*i*_ is the right eigenvector of K with eigenvalue *µ*_*i*_. It is helpful to keep in mind that just like *V*_*j*_ represents the Koopman eigenfunction *ϕ*_*j*_ to be approximate Koopman operator K in the dictionary basis *D*, 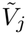 represents the dual vector 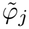 of the Koopman eigenfunction *ϕ*_*j*_ in the dual basis 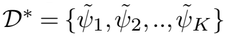 of the dictionary basis *D*. Then from Eq. (S67), we can see that

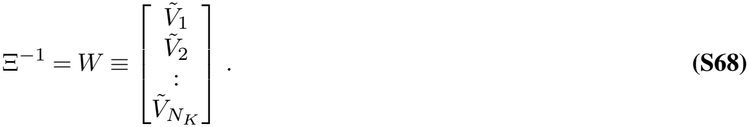

Once we have determined Ξ^−1^, we can determine the Koopman modes from Eq. (S66) as

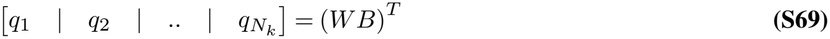

with the Koopman mode *q*_*k*_ given by

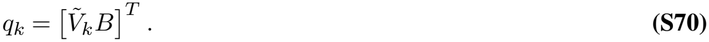

This mode *q*_*k*_ is the finite dimensional approximation of the original Koopman mode just like the corresponding eigenvalue *µ*_*k*_ and eigenvector *ϕ*_*k*_ are the finite dimensional approximations to the Koopman eigenvalue and Koopman eigenfunctions respectively. We summarize the algorithm below.

###### Algorithm summary

Using EDMD, we have determined the eigenvalues *µ*_*k*_ and the corresponding eigenvectors *ϕ*_*k*_ and modes *q*_*k*_ of the matrix K which serves as an approximation of the Koopman operator *K* restricted to the span ℱ_*D*_ of the dictionary basis, which is a finite dimensional subspace of the observable space. Consequently, the eigenvalues *µ*_*k*_ and the corresponding eigenvectors *ϕ*_*k*_ and modes *q*_*k*_ serve as an approximation to the Koopman eigenspectrum. We summarize the algorithm briefly below for simplification of th readers.

The following are the pre-requisites for EDMD:

1. A time series dataset of *M* snapshot pairs 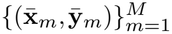 which are *M* pairs of snapshots of the system taken at some interval, i.e., 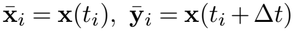. This contain information about the actual dynamics of the system on which we shall train the EDMD to capture the dynamics.
2. A dictionary basis *D* of generic basis functions *ψ*_1_, *ψ*_2_, .., *ψ*_*K*_. We will use this basis to approximate the Koopman eigenfunctions.
3. The matrix *B* defined in Eq. (S59) which contains the total information (weights) of the state vector **x** in the dictionary basis.

Once we have the prerequisites mentioned above, we are ready to apply the following steps to compute the approximate Koopman eigenspectrum.

1. Loop through the training data set 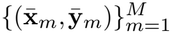 to compute the matrices *G* and *A* from Eq. (S50).
2. Once the matrices *G* and *A* are computed, compute the matrix K 6 *G*^+^*A*. This matrix K is the approximation of the Koopman operator. Find its corresponding eigenvalues *µ*_*j*_ and right eigenvectors *V*_*j*_ which are the approximation of the Koopman eigenvalues and Koopman eigenvectors respectively.
3. Compute the corresponding left eigenvectors 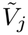 of K. From it, compute the modes 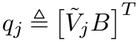 which are the approximations of the corresponding Koopman modes.

#### S1.B. Relaxation modes and invariant manifolds

##### Introduction

We mentioned in the main script that the trajectory **x**(*t*) of a dynamical system (in presence of global attractors) can be separated into two parts: one part **x**_∞_(*t*) that belongs to the invariant manifold and the remaining transient part **x**(*t*) − **x**_∞_(*t*) which consists of the relaxation modes that decay with time and eventually vanish at infinite time. Here, we present a mathematical formulation of the idea and an algorithm of how to separate the relaxation modes from the collective modes that belong to the invariant manifold. We separately consider two cases: when **x**_∞_(*t*) is a fixed point, and when **x**_∞_(*t*) is a limit cycle. We assume the system to be continuous in time, but the algorithms can also extended to discrete time case.

##### S1.B.1. Fixed point case

###### Mathematical Formulation

Let us consider a trajectory **x**(*t*) with initial condition **x**(0) = **x**_0_ and at very large time, the trajectory asymptotically approaches the fixed point **x**_*f*_ .i.e lim_*t*→∞_ **x**(*t*) = **x**_*f*_ . In that case, using Koopman spectral decomposition, the trajectory can be expressed as

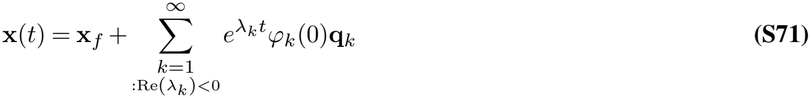

where *λ*_*k*_ is the eigenvalue corresponding to the Koopman eigenfunction *ϕ*_*k*_ and the corresponding Koopman mode is **q**_*k*_. Since Re(*λ*_*k*_) *<* 0, all the Koopman modes **q**_*k*_ decay with time and are therefore, the relaxation modes. The fixed point **x**_*f*_ corresponds to the (linear combination of) Koopman mode(s) with Koopman eigenvalue 0. To be more precise,

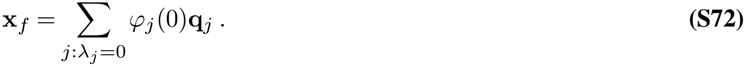

Here, the Koopman eigenfunctions *ϕ*_*j*_ : = *λ*_*j*_ = 0 in eq.Eq. (S72) belong to the same eigenbasis that we used for Koopman spectal decomposition in eq.Eq. (S71).

So, we see from eq.Eq. (S71) that if we do the Koopman spectral decomposition of a variable **y**(*t*) ≡ **x**(*t*) − **x**_*f*_ which is equivalent to a shift of the trajectory in the statespace (so that now it asymptotically approaches to 0), all the corresponding Koopman modes are indeed the relaxation modes. So, we will do Koopman mode analysis of the trajectory **y**(*t*) to get the relaxation modes.

###### Algorithm for relaxation modes and Implementation using EDMD

In order to find the relaxation modes corresponding to a fixed point, we train EDMD using sampling trajectories confined to the basin of attraction of that fixed point. We then implement the following steps:

1. We find the fixed point(s) of the sampling trajectories.
2. For each sampling trajectory **x**(*t*), we find its corresponding shifted trajectory **y**(*t*) ≡ **x**(*t*) − **x**_*f*_, where **x**_*f*_ is the fixed point of that trajectory. Theoretically, all trajectories should have same fixed point, but in numerical analysis, different trajectories may converge to different fixed points having slightly different numerical values within numerical accuracy limit.
3. We run EDMD algorithm on these shifted trajectories to numerically obtain the relaxation modes.

It is to be noted that a more direct approach is to run EDMD directly on the original trajectories without subtracting the fixed point. However, in that case, numerical precision might be lower because in that case, the fixed point is obtained from EDMD, which itself has some numerical error. But, in principle, the fixed point can be found with more numerical precision by simply running the trajectories to see where they converge, or by solving for the zeroes of the vector field.

##### S1.B.2. Limit cycle case

###### Mathematical Formulation

Let’s consider a trajectory **x**(*t*) with initial condition **x**(0) = **x**_0_ such that at infinite time, it approaches a limit cycle. Then, using Koopman spectral decomposition, we can express the trajectory **x**(*t*) as

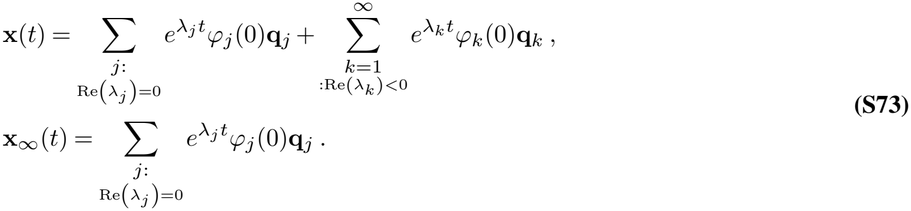

In the above equation, **x**_∞_(*t*) consists of purely oscillatory modes **q**_*j*_, with real part of the Koopman eigenvalues being zero. This part corresponds to the limit cycle and belongs to the invariant manifold. It is to be noted that different components of the Koopman modes **q**_*j*_ actually reduce to Fourier modes of the corresponding component of limit cycle **x**_∞_(*t*). The remaining part **x**(*t*) − **x**_∞_(*t*) consists of modes **q**_*k*_ which decay over time because of real part of their corresponding eigenvalues being negative,and therefore correspond to the relaxation modes. If we somehow manage to extract the part **x**_∞_(*t*) corresponding to the limit cycle, then doing Koopman mode analysis of the remaining part **x**(*t*) − **x**_∞_(*t*) give us the relaxation modes. The challenge is to separate the limit cycle part from the trajectory. We will use Fourier transform to do that.

###### Algorithm for Relaxation Modes and implementation using EDMD

The limit cycle case is mathematically more challenging as it consists of a number of points and not just a single point (unlike fixed point case). So, **x**_∞_(*t*) is not constant (unlike **x**_∞_(*t*) = **x**_*f*_ for the fixed point case) and so we need to find a way to calculate **x**_∞_(*t*) at each time *t*. We implement the following steps:

1. We choose a sufficiently late time *τ* when all the relaxation modes die off and effectively only the limit cycle survives. Of course, theoretically, the limit cycle is reached at *t* → ∞ and therefore, during numerical analysis, at some late time *τ*, the limit cycle is reached only within a certain numerical precision.
2. Let the time period of the limit cycle be *T* . Then, we perform Fourier transformation on each component of **x**(*t*) in the time interval [*τ, τ* + *T* ] to get the Fourier modes, which technically are the corresponding components of the Koopman modes.
3. Once we get the Koopman spectrum of the limit cycle from Fourier analysis, we sum over them to get the limit cycle. It is to be noted that this limit cycle is shifted forward by time *τ* . So, what we get is **x**_∞_(*t* + *τ* ). We then shift it back by time *τ* by multiplying 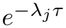 to each Fourier/Koopman mode. This numerically gives us the limit cycle **x**_∞_(*t*).
4. We subtract the numerically obtained limit cycle from original trajectories to get the shifted trajectories **x**(*t*) − **x**_∞_(*t*), which consist only the relaxation modes. We run EDMD on the these shifted trajectories to get the relaxation modes.

It is to be noted that a simpler way to compute the relaxation modes is to run EDMD on the entire trajectories. Following Eq. (S73), the Koopman eigenvalues corresponding to the modes constituting the limit cycle will be purely imaginary while relaxation modes will have negative real part. So, we can set a cut off and if the magnitude of the negative real part of the eigenvalues obtained from EDMD is smaller than the cut off, we can classify the corresponding modeas contributing to the limit cycle and part of the invariant manifold. Else, we classify it as a relaxation mode. However, since EDMD algorithm itself has some numerical error, the limit cycle obtained from EDMD likely has less numerical precision than when numerically computed using Fourier transform.

### S2: Methods

#### S2.A. Interpretation of Koopman spectrum obtained from EDMD

For the A549 cell-cycle model, there are multiple fixed points which can be broadly grouped into three clusters. Note that within each cluster the fixed points are very close to each other in the state space. So we train the EDMD models with the approximation that they share the same relaxation modes,

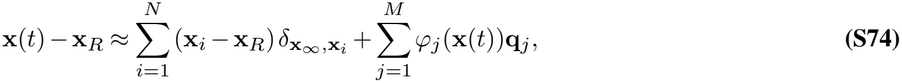

where the index *i* runs over all the fixed points {**x**_*i*_} in the cluster, {*ϕ*_*j*_, **q**_*j*_} are a finite set of *M* Koopman eigenfunctions, and Koopman modes numerically obtained from EDMD and **x**_*R*_ is an arbitrary reference point in the statespace, which can be chosen as the centroid of the cluster of fixed points. Also,

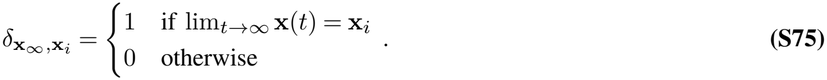

#### S2.B. Hyperparameters optimization of EDMD model

We employed Radial Basis Functions (RBFs) with Gaussian kernels as the EDMD dictionary, selecting center points via K-means clustering (41, 42). This approach introduces two hyperparameters, the dictionary size and the width of Gaussian kernels. A larger dictionary provides a larger function space ℱ_*D*_ ∈ ℱ and therefore, can better approximate the Koopman eigenfunctions but also risking overfitting. Wide Gaussian kernels capture global dynamics, whereas narrow kernels focus on local dynamics. Therefore, these two hyperparameters must be optimized to best fit a specific dynamical system. We initially tested several values for each parameter across a broad range and calculated the Mean Squared Error (MSE) between reconstructed trajectories and the real trajectories for both the training set and the test set. After that we performed a grid search over a narrower range of combinations of these two parameters.

One thing needed to mention here is that the final selection of hyperparameters is not solely based on MSE. We also considered the interpretability of the EDMD results. For example, while an EDMD model with dictionary size of 30 might yield the lowest MSE, the biological meaning of each mode in a size 20 model may be clearer. This stems from the fact that a dynamical system possesses a minimal set of Koopman eigenfunctions. Since any product of two eigenfunctions is also a Koopman eigenfunction, a smaller dictionary often yields more biologically meaningful insights.

#### S2.C. Mode selection and importance evaluation

Theoretically, the Koopman spectral decomposition can contain infinite modes. In numerical analysis, we obtain a finite but large number of modes from EDMD. Not all modes have equal contribution to the trajectory. A smaller subset of modes can be selected to reconstruct the trajectory within desired precision. To obtain a compact spectral description of the dynamics, mode selection was performed at the level of mode groups rather than individual Koopman modes. This procedure combined mode grouping, error-based selection, and amplitude-based filtering.

##### S2.C.1. Mode grouping

Complex eigenvalues appear in conjugate pairs 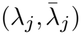 with corresponding conjugate Koopman modes 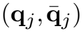. We group these conjugate pairs into mode groups; real eigenvalues form singleton groups. Each conjugate pair contributes a single real-valued oscillatory signal to the reconstruction.

##### S2.C.2. Greedy algorithm

Starting from these mode groups, we identified a reduced subset that preserved reconstruction accuracy. For each training trajectory, reconstruction was first computed using all candidate mode groups. At each iteration, one group was removed and the resulting increase in Root Mean Squared Error (RMSE) was evaluated as,

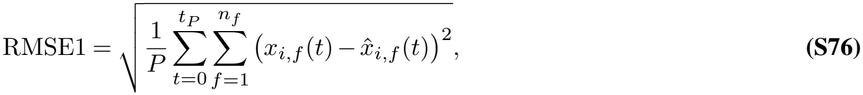

where the time interval is discretized in *P* time snapshots {*t*_1_, *t*_2_, …, *t*_*P*_ }, *n*_*f*_ is the number of state variables and 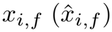 denote the *f* ^th^ component of the training (reconstructed) trajectory 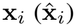. The group whose removal produced the smallest increase in RMSE was discarded. This procedure was repeated until any further removal caused an error increase larger than a preset threshold.

Let *S* denote the current set of mode groups, and let RMSE(*S*) denote the reconstruction error obtained from that set. RMSE(S) is basically RMSE1 (see Eq. (S76)) evaluated using reconstructed trajectory 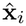 obtained from the mode groups in *S*. At each step, the removed group is

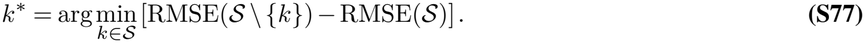

Elimination stops when the smallest possible increase in error exceeds a preset threshold *δ*:

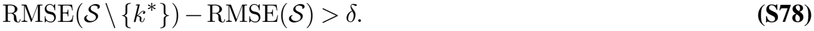

In three variables cell-cycle model, we use *δ* = 0.001 for S = 0.1, S = 0.35 lower branch fixed point, S = 0.35 upper branch fixed point, and S = 1.5. In A549 model, the *δ* is scaled with the root mean square (RMS) of each trajectory, defined as 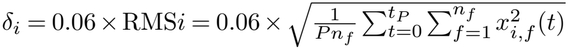. Here, RMS*i* denotes the RMSE without the reconstructed trajectory, that is, with 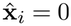. Because the contribution of a mode group may vary across initial conditions, this elimination procedure was carried out separately for all training trajectories. A mode group was retained in the final set only if it remained selected in a sufficient fraction of trajectories. In this way, the final subset reflects mode groups that are repeatedly required across the data set, rather than groups that are important only in a small number of cases.

If *S*_*i*_ denotes the retained set for trajectory *i*, the retention fraction of mode group *k* is

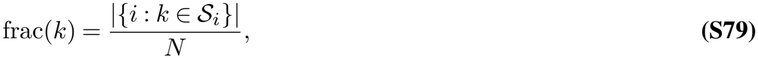

where *N* is the number of training trajectories and {*i* : *k* ∈ S_*i*_} is the set of training trajectories where mode group *k* was retained. As we will see shortly, we use retention fraction to set thresholds for mode selection. A given mode group was *k* was selected if frac(*k*) ≥ *γ*. The mode groups that were finally selected are designated as the *dominant* modes which are needed for trajectory reconstruction. In the three variables cell-cycle model, we reconstructed the trajectories using the retained modes obtained from greedy selection and compared them with ground truth trajectories to decide a reasonable threshold. More specifically, we calculated the RMSE at different thresholds and chose values that reduced the number of mode groups while keeping the reconstruction error low. The thresholds were *γ* =0.6, 0.6, 0.15, and 0.4 for *S* = 0.1, *S* = 0.35 lower branch fixed point, *S* = 0.35 upper branch fixed point, and *S* = 1.5 respectively. For A549 cell cycle model, we used retention fraction threshold of *γ* = 0.6. More on this is elaborated at the end of section S2.C.3.

##### S2.C.3. Amplitude analysis

An error-based selection method like the greedy algorithm identifies mode groups that are needed for accurate reconstruction, but it does not directly measure the size of their dynamical contribution. We therefore quantified the time dependent amplitude of each selected mode group from its reconstructed signal.

For trajectory *i*, the contribution of mode group *k* to feature *f* at time *t* is^6^

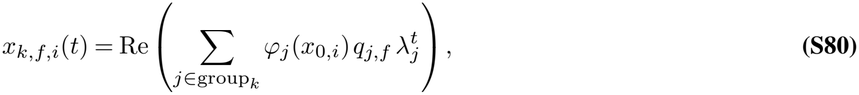

where *ϕ*_*j*_(*x*_0,*i*_) is the Koopman eigenfunction evaluated at the initial state, *q*_*j,f*_ is the *f* th entry of the *j*th Koopman mode **q**_*j*_, and *λ*_*j*_ is the corresponding eigenvalue. Please note that EDMD is a numerical algorithm and operates on discrete time. So, all expressions involving Koopman decomposition are expressed here in discrete time format.

The amplitude of mode group *k* at time *t* was defined as

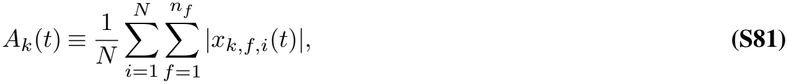

where *n*_*f*_ is the number of state variables, and *N* as mentioned earlier, is the number of training trajectories. To compare mode groups on the same scale, we further defined the amplitude fraction

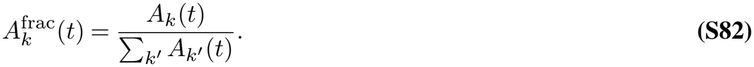

Mode groups with negligible time-averaged amplitude fraction were excluded from the final set. This filtering step removed groups whose contribution remained weak even if they survived the error-based selection step.

Ideally, while selecting dominant mode groups, both retention fraction and amplitude fraction should be taken into consideration simultaneously. As we will see in section S3, for A549 cell cycle model, the retention fraction threshold of *γ* = 0.6 is consistent with the selected modes having high amplitude fraction. For the three-variable cell-cycle model, we actually do a two-step selection process. First we use the greedy algorithm and select modes *above* a certain threshold, as mentioned in section S2.C.2, and then we look at the mean amplitude fraction of those modes. For each mode group, we sum the amplitude fraction across all time points for each trajectory and then calculated the average among trajectories. Then finally select mode groups *above* a certain amplitude fraction threshold. In toy model, we use amplitude threshold 0.01, 0.01, 0.01, and 0.004 for S = 0.1, S = 0.35 lower branch fixed point, S = 0.35 upper branch fixed point, and S = 1.5 respectively.

#### S2.D. Trajectory reconstruction

After mode selection, trajectories were reconstructed by summing the contributions of the retained mode groups. For an initial state **x**_0_, the reconstructed trajectory is

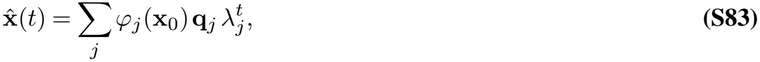

where the sum runs either over all mode groups or over the selected subset, depending on the comparison.

Reconstruction was used in two ways. First, we compared the trajectory reconstructed from all mode groups with that reconstructed from the selected subset (consisting of dominant modes) to determine whether a reduced spectral representation was sufficient to preserve the dominant transient structure. Second, we monitored reconstruction error as mode groups were incorporated progressively, providing a compact summary of their relative contributions to the total fit.

Reconstruction accuracy was quantified by the root mean squared error,

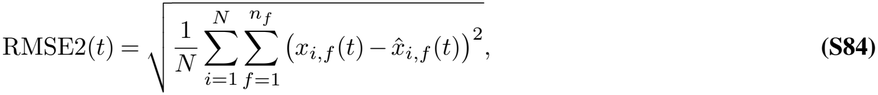

where *N* is the number of trajectories, *n*_*f*_ is the number of state variables and 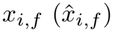 denote the *f* ^th^ component of the training (reconstructed) trajectory 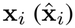. Please note that the reconstruction accuracy RMSE2(*t*) used here is different from RMSE1(*t*) which was used during greedy selection.

In the fixed-point regimes, the retained mode groups primarily describe transient relaxation toward equilibrium. In the limit-cycle regime, separating the persistent oscillatory group from the transient groups allowed reconstruction to distinguish the long-term periodic motion from the decaying approach toward that motion.

#### S2.E. Normalization of Koopman modes

We applied a three-step normalization to place different modes on comparable scales and identify genes that contribute disproportionately strongly to individual modes. Let *q*_*ij*_ denote the complex Koopman coefficient associated with gene *i* and mode **q**_*j*_.

First, each mode was normalized to unit *L*_2_ norm across genes to remove differences in overall amplitude across modes:

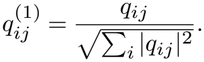

Next, each gene was normalized by its mean absolute coefficient across all modes:

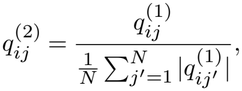

where *N* is the number of modes included in the normalization. This step reduces the contribution of genes that have consistently large coefficients across many modes and emphasizes genes whose contributions are disproportionately large in particular modes. Without this step, the GO terms enriched in different modes were highly similar.

Finally, each mode was normalized again to unit *L*_2_ norm:

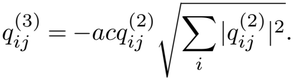

Because we used pre-ranked GSEA, this final normalization does not change the gene ranking within each mode and therefore does not affect the GSEA results. It was included to place the final normalized modes on a common scale.

#### S2.F. Coarse graining and gene clustering

An important application of EDMD in this study is gene clustering. We used Koopman modes to cluster genes and compared the results with those obtained using conventional clustering methods. Here, we focused on the 65 genes that constitute the core gene regulatory network.

##### S2.F.1. Koopman mode based gene clustering

For each gene *i*, we constructed a vector with its components from selected dominant complex conjugate Koopman mode pairs from all three basins of attraction:

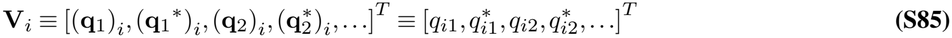

where *q*_*ij*_ ≡ (**q**_*j*_)_*i*_ is the complex Koopman mode component for gene *i* in mode *j*. The modes included in the vector can be selected depending on the analysis. The gene clustering results showed in main text included the dominant modes identified from all three basins. As a robustness check, we repeated the clustering using all Koopman modes and obtained highly similar clustering results (Fig. S2.FB).

We then calculated the pairwise distance between gene vectors using normalized Hermitian inner product 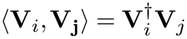 so that we have

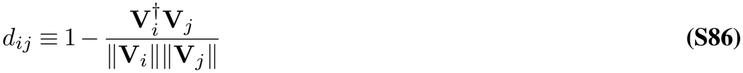

where 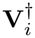 is the Hermitian conjugate of **V**_*i*_ and the norm ∥ **V**_*i*_ ∥ is defined using the Hermitian inner product so that

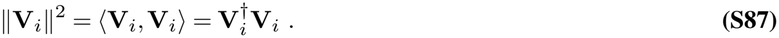

This formula removes differences in the overall magnitude of gene vectors, preventing genes with uniformly large Koopman mode component from receiving disproportionately large weights in the clustering. At the same time, it preserves the relative amplitudes of a gene across different modes as well as the phase information within each complex mode. Therefore, genes were clustered according to their patterns of contribution across dynamical modes rather than their overall coefficient magnitudes. Then, hierarchical agglomerative clustering was performed using the resulting pairwise cosine distances with average linkage. At each step, the distance between two clusters was defined as the average pairwise distance between all genes belonging to the two clusters. The resulting hierarchical tree was cut at a distance threshold of 0.75 to define the final gene clusters. Clustering was implemented using the ‘pdist’, ‘linkage’, and ‘fcluster’ functions in SciPy.

##### S2.F.2. Expression based gene clustering

As a comparison with Koopman mode-based clustering, we also clustered genes directly based on their expression profiles in the original processed scRNA data (which is also the first time point in trajectory data). For each core gene *i*, we constructed a vector containing its expression levels across all cells or trajectories at the first time point,

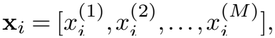

where *M* is the number of cells or trajectories included in the analysis.

Each gene expression vector was independently standardized using a z-score,

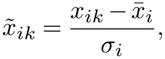

where 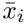 and *σ*_*i*_ are the mean and standard deviation of gene *i* across cells or trajectories, respectively. This removes differences in the overall expression level and scale of variation among genes and allows the comparison to focus on their relative expression patterns across the cell population.

Pairwise distances between genes were then calculated using correlation distance,

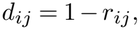

where *r*_*ij*_ is the Pearson correlation coefficient between the expression profiles of genes *i* and *j*. The same hierarchical agglomerative clustering was subsequently performed using average linkage.

##### S2.F.3. Trajectory based gene clustering

As a dynamical comparison with Koopman-mode-based clustering, we also clustered genes directly based on their reconstructed expression trajectories. For each core gene *i*, its expression values across all trajectories and all time points were flattened into a single vector,

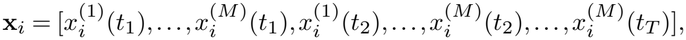

where *M* is the number of trajectories and *T* is the number of time points.

Each gene vector was standardized using the same z-score normalization as in the expression-based gene clustering analysis. Pairwise correlation distances between genes were then calculated using the same procedure described above, followed by hierarchical agglomerative clustering with average linkage.

##### S2.F.4. Comparison of gene clustering methods

To compare the gene relationships captured by different clustering approaches, we compared the pairwise gene-distance matrices obtained from different representations described above. For each representation, the upper-triangular elements of the pairwise distance matrix were represented as a one-dimensional distance vector,

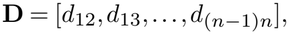

where *d*_*ij*_ is the distance between genes *i* and *j*, and *n* is the number of genes.

The similarity between two representations was quantified by the Spearman rank correlation coefficient between their corresponding pairwise distance vectors. Spearman correlation measures whether the relative ordering of gene–gene distances is preserved between two representations without requiring a linear relationship or identical distance scales. A high positive correlation therefore indicates that two representations capture similar relative relationships among genes.

##### S2.F.5. Calculating edge-strength of coarse-grained network

After obtaining the gene clusters, we sought to simplify the gene regulatory network through coarse-graining. We constructed two types of coarse-grained networks using different definitions and methods: an overall network and a network representing a single time point.

To define an ‘effective’ interaction edge between two clusters in the overall coarse-grained network shown in Fig. 4C, we summed the weights of all one-to-one regulatory edges connecting each pair of genes between those two clusters, assigning positive values to activation and negative values to inhibition. A positive sum was interpreted as a net activating interaction between the two clusters, whereas a negative sum was interpreted as a net inhibitory interaction.

To construct the time-point-specific coarse-grained network in Fig. 4E, we incorporated the expression levels of regulator genes to estimate the activity of individual regulatory interactions. For each one-to-one regulatory edge of activation and inhibition, the edge activities were defined as

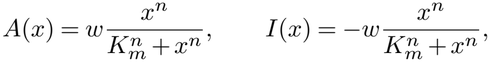

respectively, where *x* is the expression level of the regulator gene at the selected time point, and *w, n*, and *K*_*m*_ are the edge weight, Hill coefficient, and half-maximal concentration, respectively, as determined by Dynamo. The resulting edge activities were aggregated according to the clusters of their regulator and target genes. Specifically, for each pair of clusters, we summed the signed activities of all regulatory edges connecting genes in the regulator cluster to genes in the target cluster. Positive values indicate activating interactions, whereas negative values indicate inhibitory interactions between gene clusters.

#### S2.G. Gene Ontology enrichment analysis

To characterize the biological functions associated with Koopman modes and gene clusters, we performed functional enrichment analysis using GSEApy with Gene Ontology (GO) Biological Process terms from the `GO_Biological_Process_2021` gene set library.

For Koopman modes, we applied the three-step normalization described above and ranked genes according to the absolute value of their normalized Koopman coefficients, 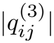. Preranked gene set enrichment analysis (GSEA) was then performed with 1,000 permutations and gene-set sizes restricted to 5–1,000 genes. Enrichment was quantified using the normalized enrichment score (NES), and the top 10 enriched Biological Process terms were visualized for each mode pair.

For gene clusters, we performed over-representation analysis using the `enrichr` implementation in GSEApy. The genes assigned to each cluster were used as the input gene set, and enrichment was assessed based on their overlap with genes annotated to each GO term. Enrichment significance was evaluated using *P* -values adjusted for multiple testing using the Benjamini–Hochberg procedure. The top 10 enriched Biological Process terms ranked by adjusted *P* -value were visualized for each gene cluster.

### S3: Results

#### S3.A. Three-variable cell cycle model

Here, we discuss the results of the three-variable cell cycle model. Since in the main text, we showed results of Koopman analysis for the monostable regime (*S* = 0.1), here we show the results for the remaining two cases: Bistable regime (*S* = 0.35) and limit cycle (*S* = 1.5).

##### Bistable regime

In the bistable regime, we show the results of Koopman mode analysis performed using EDMD algorithm for *S* = 0.35. As mentioned briefly in the main text, there are two fixed point attractors in this region for each value of *S*, on the bottom most and top most branch of the bifurcation diagram, which we call Fixed point 0 and fixed point 1 respectively. We present the sample input trajectories of fixed points 0 and 1 in the left and right column of Figure Panels Fig-1D.

At *S* = 0.35, the two fixed points show different relaxation dynamics and correspond to two different stable arrest regimes. For fixed point 0, modes [1, 2], [3, 4], and [5, 6] have the highest retention fractions and large amplitude fractions among the relaxation modes (Fig. S2B,C). The selected modes reproduce the main trajectory structure, while reconstruction using all modes more closely follows the original trajectories (Fig. S2D). Mode [1, 2] decays more slowly and falls to the fixed point in a long time. Mode [3, 4] shows a damped oscillatory pattern and decays faster than mode [1, 2]. Mode [5, 6] is mainly present at early times and decays rapidly (Fig. S2E–G).

The three modes also show different distributions across the three variables. Mode [1, 2] has contributions in all three variables and is the most global of the displayed relaxation modes. Mode [5, 6] has a more restricted contribution, with most of its change occurring during the early transient. Mode [3, 4] shows an intermediate pattern. After the faster modes decay, the late time relaxation toward fixed point 0 is mainly represented by mode [1, 2], with a smaller contribution from mode [3, 4].

For fixed point 1, the mode structure is different. Modes [1], [2, 3], and [9] show the main contributions among the relaxation modes (Fig. S3B,C). It is to be noted that although mode group [11,12] has a slightly higher retention fraction than mode [1] (see Fig. S3B), its contribution to relative amplitude fraction is negligible (see Fig. S3C). This is because mode [11,12] is a very fast decaying mode with important contribution at very early transient behavior, but relatively insignificant contribution to long term dynamics. As for fixed point 1, similar to fixed point 0, the selected modes reproduce the main trajectory structure, while the reconstruction using all modes more closely follows the original trajectories (Fig. S3D). Mode [2, 3] gives a clear damped oscillatory contribution in CycB, Cdh1, and Cdc20. Mode [1] gives a slower non-oscillatory contribution. Mode [9] is mainly present at the beginning of the trajectory and decays rapidly (Fig. S3E–G).

Among these modes, mode [2, 3] has the broadest contribution across the three variables. Mode [9] is more restricted to the early transient. At later times, the contributions from modes [1] and [2, 3] remain after mode [9] has mostly decayed. These modes describe the slow relaxation near fixed point 1.

Biologically, fixed point 0 represents a more direct arrest state, where the regulatory network quickly relaxes into a steady state. Fixed point 1 corresponds to an arrest state in which the system shows transient oscillatory regulatory dynamics before reaching to the steady state. This suggests that the upper-branch state still retains stronger cyclic regulatory structure, such as delayed feedback among CycB, Cdh1, and Cdc20, before reaching the fix point. Therefore, the two fixed points are not just different steady-state expression levels, they also represent different dynamical mechanisms of arrest.

##### Oscillatory regime

At *S* = 1.5, the modes associated with the sustained limit cycle are separated from the relaxation mode analysis. The remaining modes have negative real parts and describe the transient approach to the limit cycle. Among these modes, [13, 14], [15], [18], [21], and [24, 25] were kept in the greedy selection and amplitude analyses (Fig. S4B,C). Combining the selected relaxation modes describes the transient approach to the limit cycle, while reconstruction using all Koopman modes also reproduces the sustained oscillation (Fig. S4D).

Mode [13, 14] shows the slowest decay among the displayed relaxation modes. It gives a damped oscillatory contribution and remains visible after the other relaxation modes become small. Modes [15], [18], and [21] mainly contribute during the early transient and decay more rapidly. Mode [24, 25] also shows an oscillatory contribution, but its amplitude decreases faster than mode [13, 14] (Fig. S4E–G).

The late time dynamics contain two parts. The sustained limit cycle is represented by the modes with almost zero (Because of numerical error, it is not exactly 0) real part of the Koopman eigenvalue, while mode [13, 14] gives the slowest remaining relaxation toward the limit cycle. The other displayed relaxation modes mainly contribute at earlier times. As mentioned earlier, combining the selected relaxation modes describes the transient approach to the limit cycle.

#### S3.B. A549 cell cycle model

##### S3.B.1. Three attractors

As briefly mentioned in the main text, three attractors were identified in the A549 cell-cycle model. We grouped the simulated trajectories according to the attractors that they approached. To characterize the corresponding basins of attraction, we examined the initial cell-cycle phases and the expression profiles of the final states from trajectories.

The initial cell-cycle phases were inferred using Dynamo (43). In basin 1, 61.3% of cells were initially labelled M or M/G1. In basin 2, 78.9% of cells were initially labelled G1/S. In basin 3, 74.7% of cells were initially labelled S or G2/M.

We next examined the expression profiles of the final states. For each basin, we calculated the median expression of each gene across the final states of trajectories. We compared genes involved in DNA replication and its regulation (44–48) and genes involved in mitotic regulation (49, 50) (Fig. S6). Basin 2 had the highest median expression of all ten genes in the replicationrelated panel, consistent with an expression profile associated with S phase. Basin 1 showed lower expression of these genes than basin 2 and the lowest expression of the three mitotic regulators, suggesting a G1-like state. Basin 3 had the highest expression of the three mitotic regulators, which is consistent with a state associated with G2.

The trajectories in basins 1, 2, and 3 converged to fixed points associated with G1, S, and G2, respectively. This suggests that cells finally stuck in these phases and did not progress to the next phase during the cell cycle.

In the main text, we use the M/G1 basin as an example to present the EDMD analysis in detail, including the interpretation of collective modes, identification of the slow manifold, and GO analysis. Here, we apply the same analysis to the S and G2 basins. To avoid repeating the same procedures, we mainly focus on the results, including the slow manifolds, GO analysis and biological interpretation of the dominant modes.

##### S3.B.2. S basin

We performed similar greedy selection, and amplitude analysis for the S basin (Fig. S8). The results showed that there are only two dominant modes, and these two modes can satisfactorily reconstruct the dynamics. Both modes have small real parts and are among the slowest modes. However, both modes are relatively local in gene space, meaning that they are dominated by only a small number of genes (Fig. S14F-J). Interestingly, the projections of these two modes largely overlap in PCA space (Fig. S10B). Consistent with this similarity, the GO terms of the two modes are also highly similar and are mainly enriched in mitosis-related processes (Fig. S13B).

##### S3.B.3. G2 basin

The greedy selection and amplitude analysis of the G2 basin identified three dominant modes (Fig. S9). These modes belong to the slowest group, although they are not always broadly distributed across cell or gene space (Fig. S14K-O). Two of the three dominant modes largely overlap in PCA space (S15C), suggesting that they represent similar dynamical directions. GO analysis showed that these two highly correlated modes are mainly enriched in DNA replication-related processes, whereas the third mode is mainly enriched in mitosis-related processes (Fig. S13C).

---

**Fig. S1.**
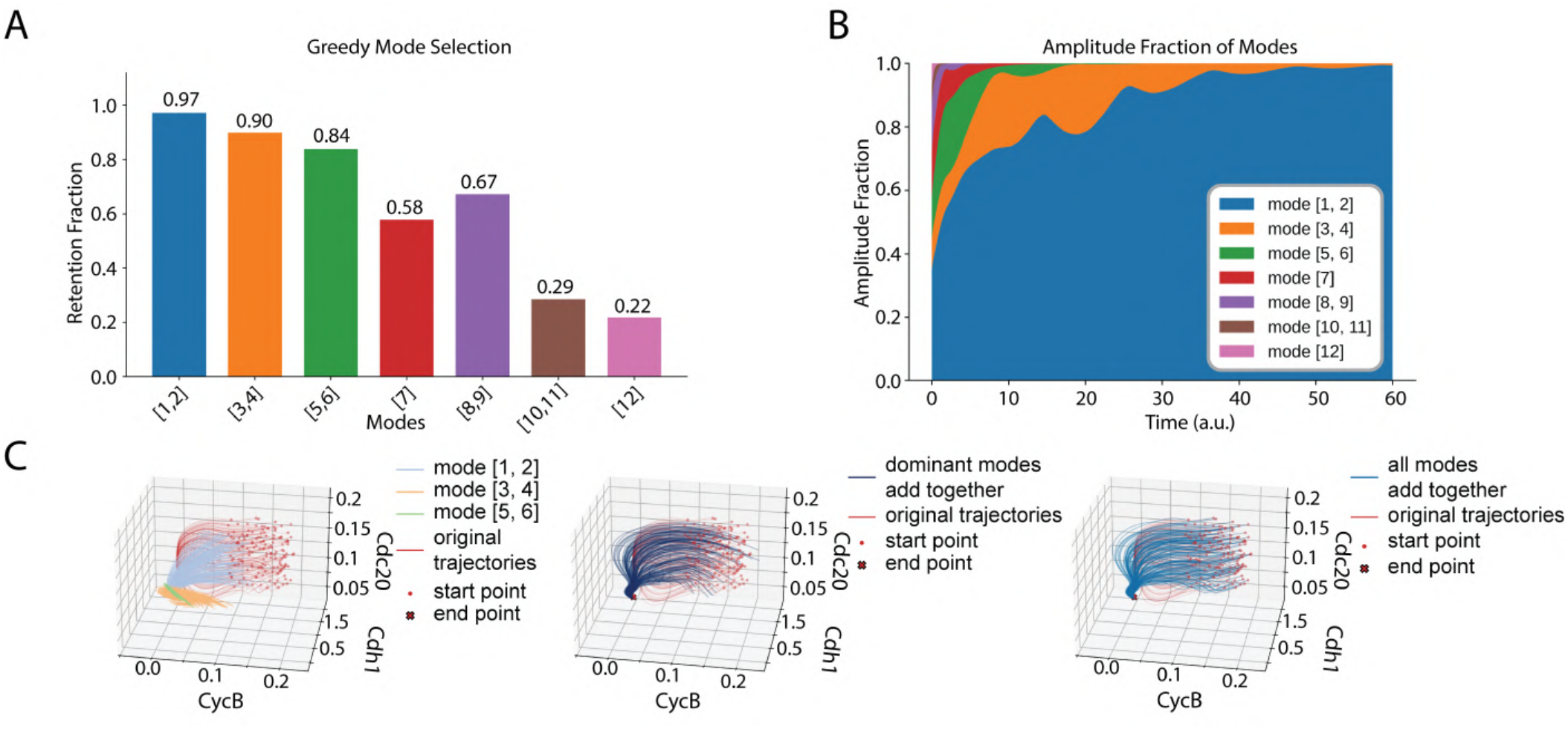
Additional results of Koopman mode decomposition and reconstruction of trajectories at *S* = 0.1. (A) Greedy mode selection results (see Methods). The bars show the retention fraction of modes. Modes [1, 2], [3, 4], and [5, 6] were selected with the highest probability. (B) Relative amplitude fractions of various mode groups over time. See methods for amplitude analysis. (C) Trajectory reconstruction in the CycB, Cdh1, and Cdc20 state space. Left: original trajectories and the sum of each of the three selected conjugate mode pairs; Middle: reconstructed trajectories using the three selected mode pairs; Right: reconstructed trajectories using all collective modes.

**Fig. S2.**
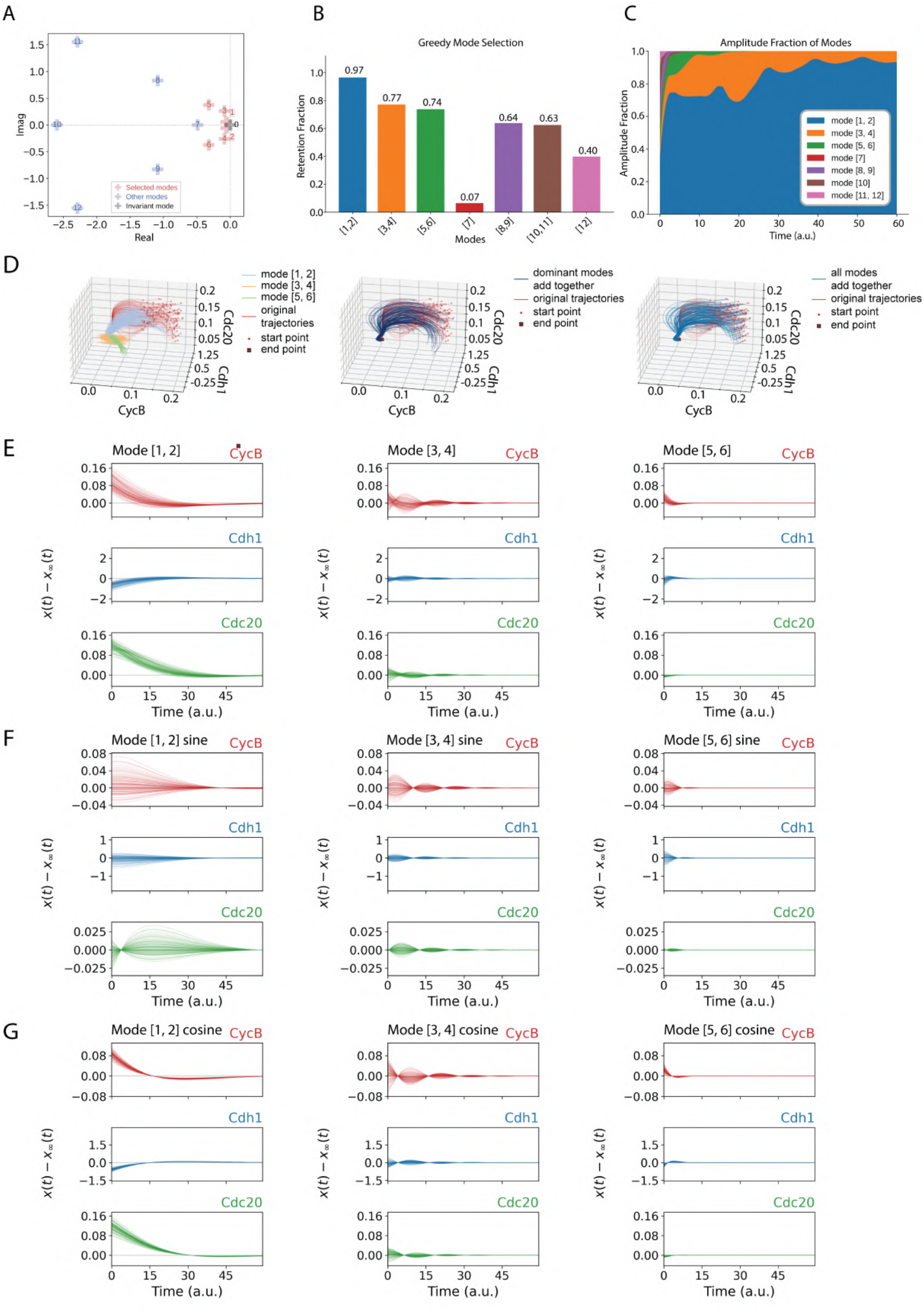
Results of Koopman mode decomposition and reconstruction of trajectories converging to fixed point 0 (the fixed point attractor on the lower branch) at *S* = 0.35 (A) Continuous Koopman eigenvalue spectrum. Modes [1, 2], [3, 4], and [5, 6] are highlighted in red. Mode [0] has eigenvalue very close to 0 and represents the fixed point contribution within numerical precision, and is represented using black cross. All other modes are shown in blue. (B) Greedy mode selection results. The bars show the retention fraction of modes. Modes [1, 2], [3, 4], and [5, 6] were selected with the highest probability. (C) Relative amplitude fractions of different mode groups over time. (D) Trajectory reconstruction in the CycB, Cdh1, and Cdc20 state space. Left: original trajectories and three selected mode pairs; Middle: Reconstruction using the three selected mode pairs [1,2], [3,4], [5,6]; Right: Reconstruction using all the modes. (E-G) The overall sum (see Eq. (16)) and the cosine and sine terms (see Eq. (17)) of three selected mode pairs [1,2], [3,4], [5,6] to CycB, Cdh1, and Cdc20 for individual trajectories, after moving the steady state contribution **x**_*∞*_(*t*) which in this case is time invariant fixed point.

**Fig. S3.**
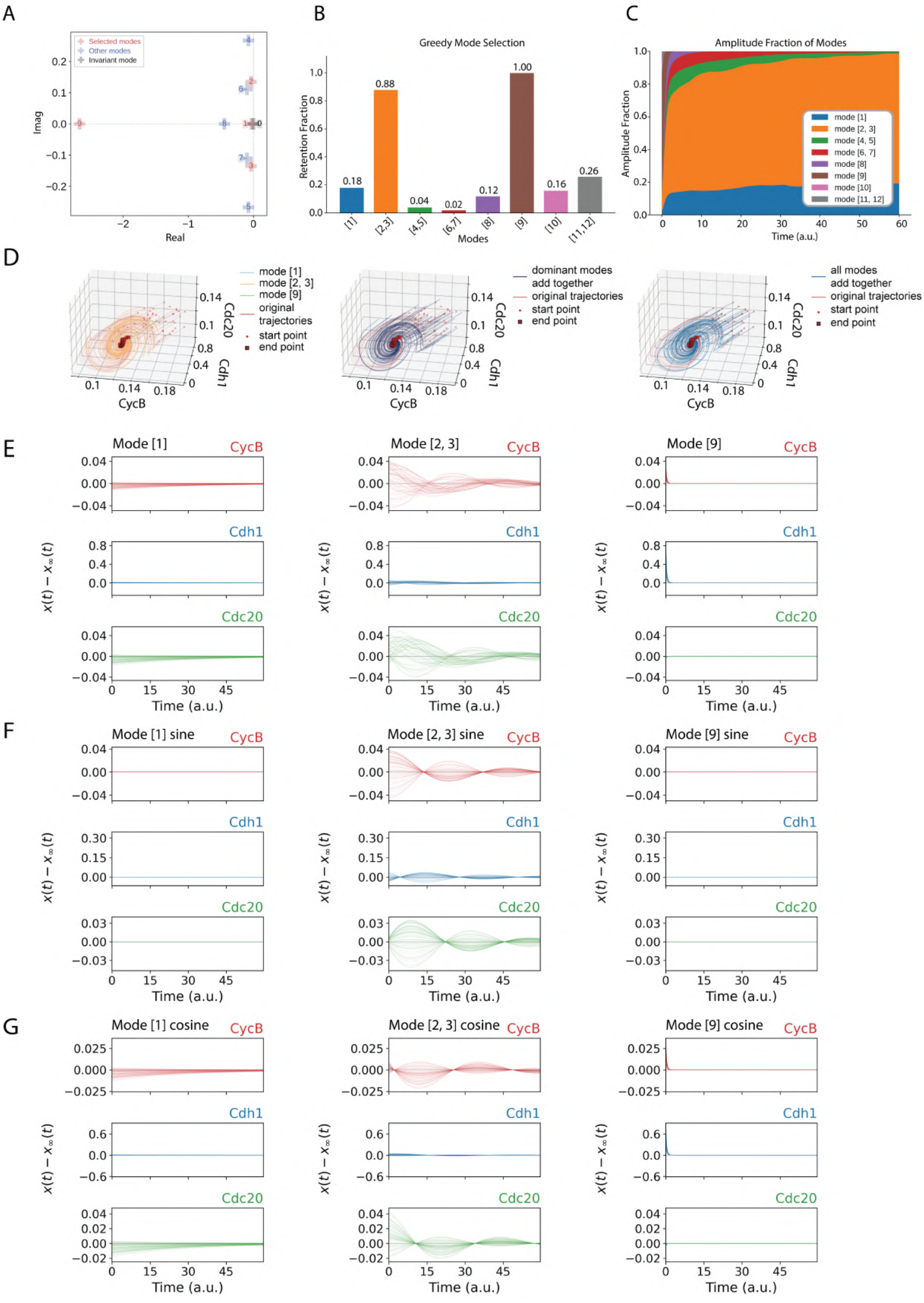
Koopman mode decomposition and reconstruction of trajectories converging to fixed point 1 (the fixed point attractor on the upper branch) at *S* = 0.35. Results of Koopman mode decomposition and reconstruction of trajectories converging to fixed point 1 (the fixed point attractor on the lower branch) at *S* = 0.35 (A) Continuous Koopman eigenvalue spectrum. Modes [1], [2, 3], and [9] are highlighted in red. Mode [0] has eigenvalue very close to 0 and represents the fixed point contribution within numerical precision, and is represented using black cross. All other modes are shown in blue. (B) Greedy mode selection results. The bars show the retention fraction of modes. Modes [1], [2, 3],[9] and [11,12] were selected with the highest probability. (C) Relative amplitude fractions of different mode groups over time. (D) Trajectory reconstruction in the CycB, Cdh1, and Cdc20 state space. Left: original trajectories and three selected mode pairs; Middle: Reconstruction using the three selected mode (pairs) [1], [2,3] and [9]. Mode group [11,12] is dropped because of its negligible amplitude fraction (see results); Right: Reconstruction using all modes. (E-G)The overall sum (see Eq. (16)) and the cosine and sine terms (see Eq. (17)) of three selected mode (pairs) [1], [2,3] and [9] to CycB, Cdh1, and Cdc20 for individual trajectories.

**Fig. S4.**
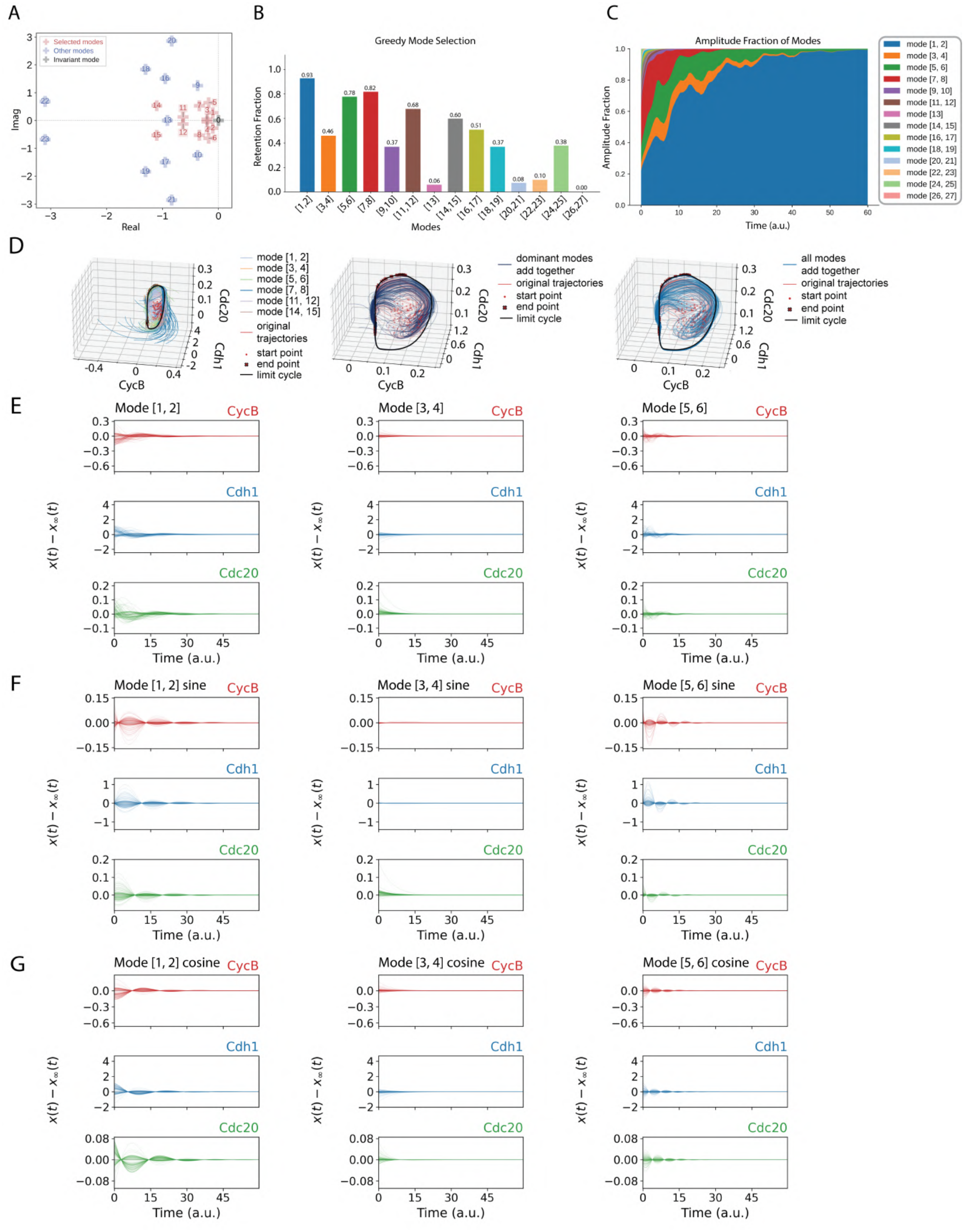
Some results of Koopman mode decomposition and reconstruction of trajectories converging to the limit cycle at *S* = 1.5. (A) Continuous Koopman eigenvalue spectrum. Modes [1, 2], [3, 4], [5, 6], [7, 8], [11, 12],and [14, 15] are highlighted in red. Before training the EDMD with sample trajectories, limit cycle was subtracted and these shifted trajectories **x**(*t*) − **x**_*∞*_, which fall into origin (which now act as the fixed point), were trained on EDMD. Mode [0] has eigenvalue very close to 0 and represents the origin fixed point contribution within numerical precision, and is represented using black cross. All other modes are shown in blue. (B) Greedy mode selection results. The bars show the retention fraction of modes. Modes [1, 2], [5, 6], [7, 8] were selected with the highest probability. (C) Relative amplitude fractions of different mode groups over time. (D) Trajectory reconstruction in the CycB, Cdh1, and Cdc20 state space. Left: original trajectories and the six selected mode pairs; Middle: Reconstruction using the six selected mode pairs; Right: Reconstruction using all modes. (E-G) Contribution from selected modes [1, 2], [3, 4], [5, 6] showing the overall sum (see Eq. (16)) and the cosine and sine terms (see Eq. (17)) to CycB, Cdh1, and Cdc20 for individual trajectories, after removing the steady state contribution **x**_*∞*_(*t*) which in this case is time varying limit cycle. Remaining modes are shown in S5.

**Fig. S5.**
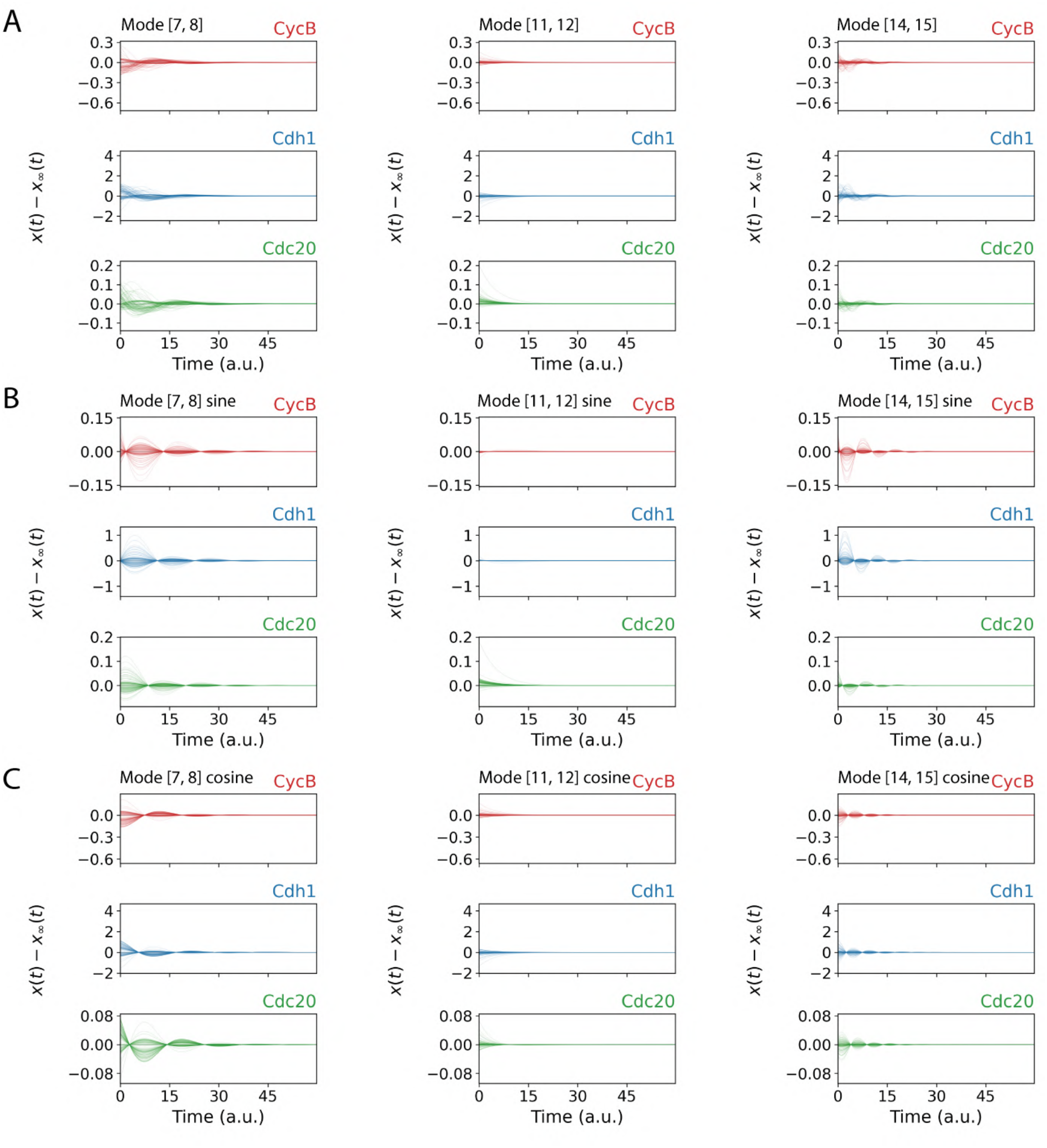
Additional results of Koopman mode decomposition for *S* = 1.5 showing the overall sum (see Eq. (16)) and the cosine and sine terms (see Eq. (17)) if the remaining selected modes [7, 8], [11, 12], [14, 15]

**Fig. S6.**
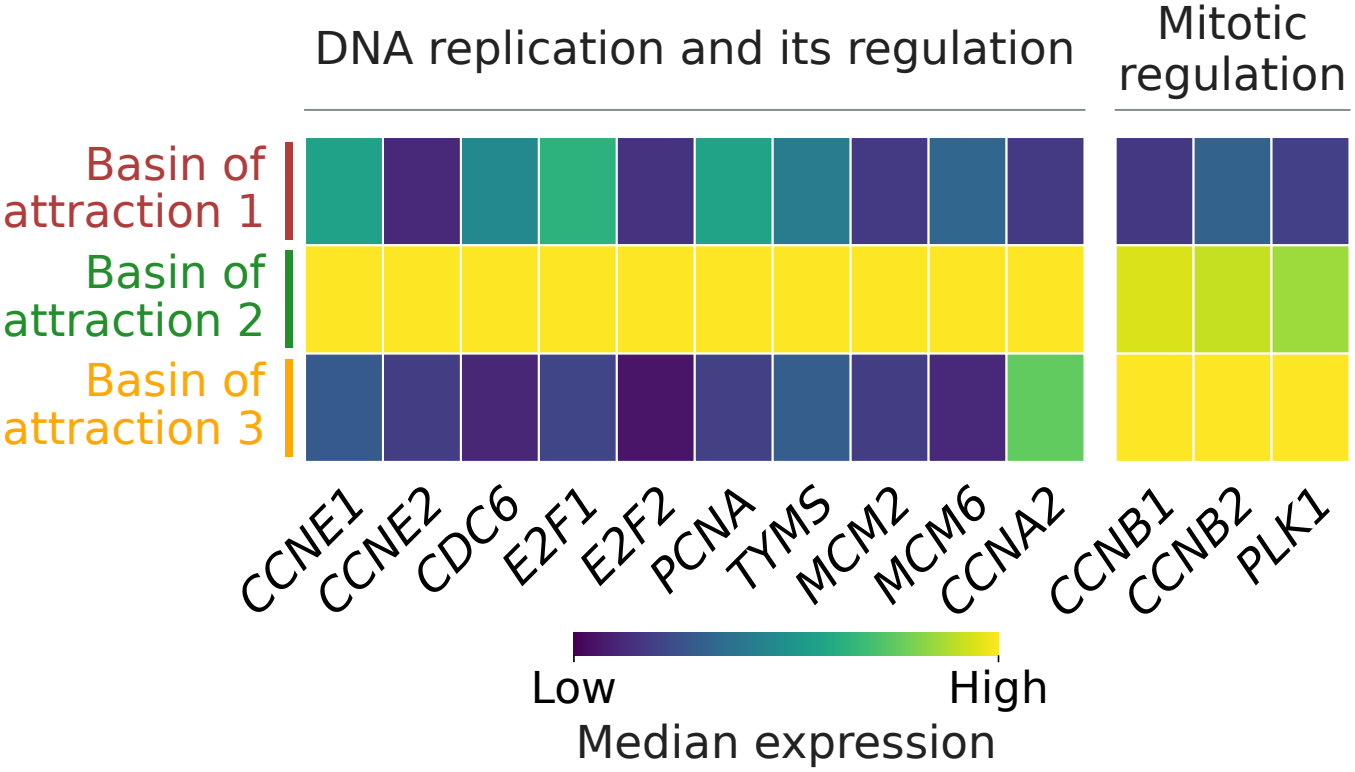
Three basins of attraction in the A549 cell-cycle model. Expression profiles of DNA replication related and mitosis related genes in each basin of attraction. The median expression level of each gene within each fix point group is shown. Based on expression profiles, the basins are designated as belonging to M/G1, S and G2 phases of cell-cycle respectively.

**Fig. S7.**
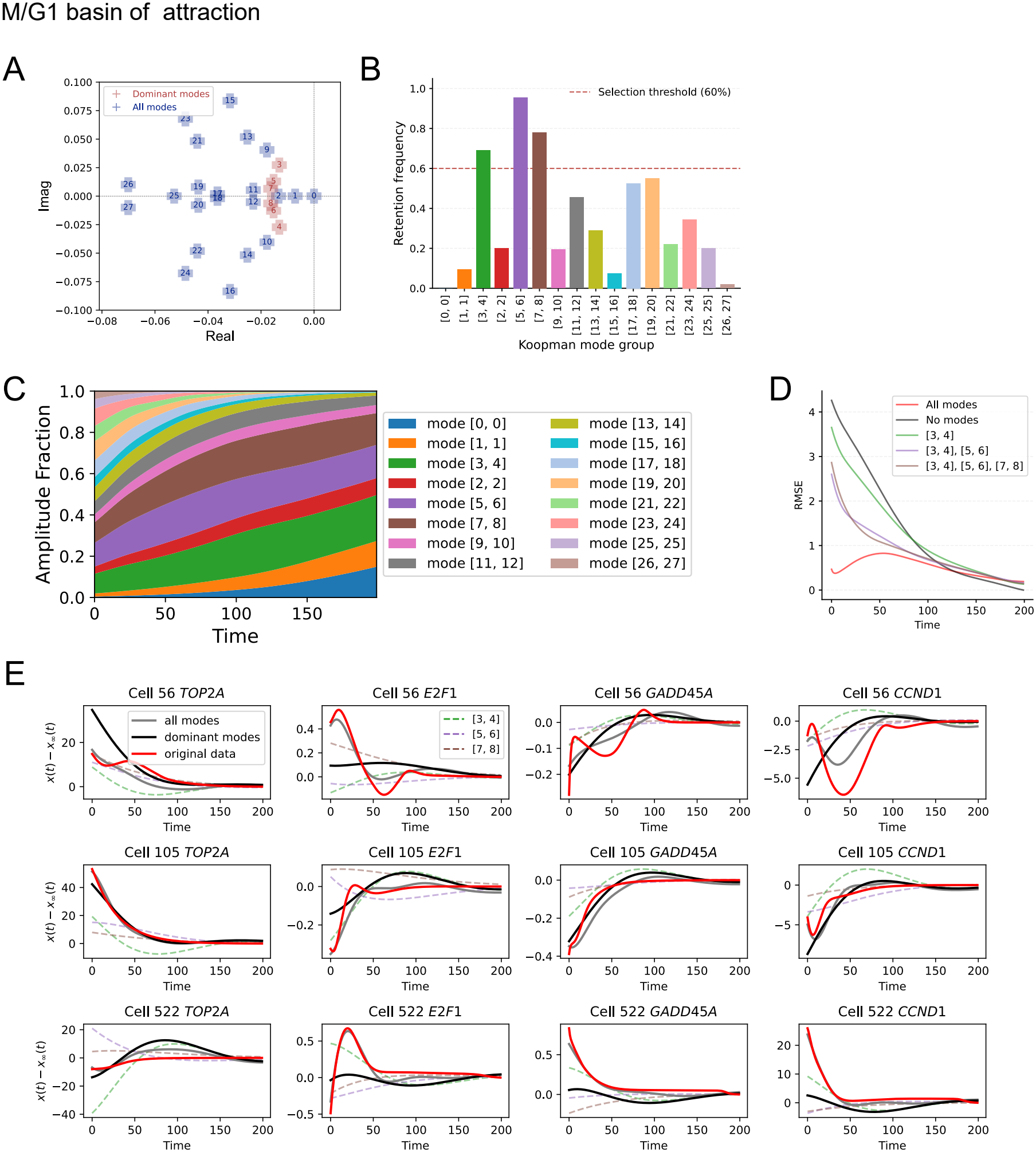
Dominant mode selection for the M/G1 basin. **(A)** Spectrum of Koopman eigenvalues. The dominant modes are highlighted in red. **(B)** Greedy selection of EDMD modes (see Methods). Modes retained in more than 60% of the trajectories are identified as dominant modes. Only mode pairs [3,4], [5,6], and [7,8] meet this threshold. **(C)** Amplitude analysis (see Methods). Modes [3,4], [5,6], and [7,8] have the largest amplitudes. Together with the greedy selection results, we identified these three mode pairs as the dominant modes in the M/G1 basin. **(D)** Reconstruction RMSE (see Methods) at each time point as the three dominant modes are added sequentially. The RMSE values are averaged across all genes and cells. The reconstruction error is largest at early times and is substantially reduced when all modes, including the faster modes, are included, suggesting that the three dominant mode pairs mainly capture long-term relaxation dynamics. **(E)** Reconstruction of four cell cycle marker genes for three randomly selected cells, after removing the steady state contribution x∞(*t*) which in this case is time invariant fixed point. Contributions from individual modes are shown in different colors and labeled accordingly. The original trajectory is shown as a bold red curve, while trajectories reconstructed using all modes or only the dominant modes are shown in gray and black, respectively. Trajectories reconstructed using only the three dominant mode pairs capture the overall trends but miss some details.

**Fig. S8.**
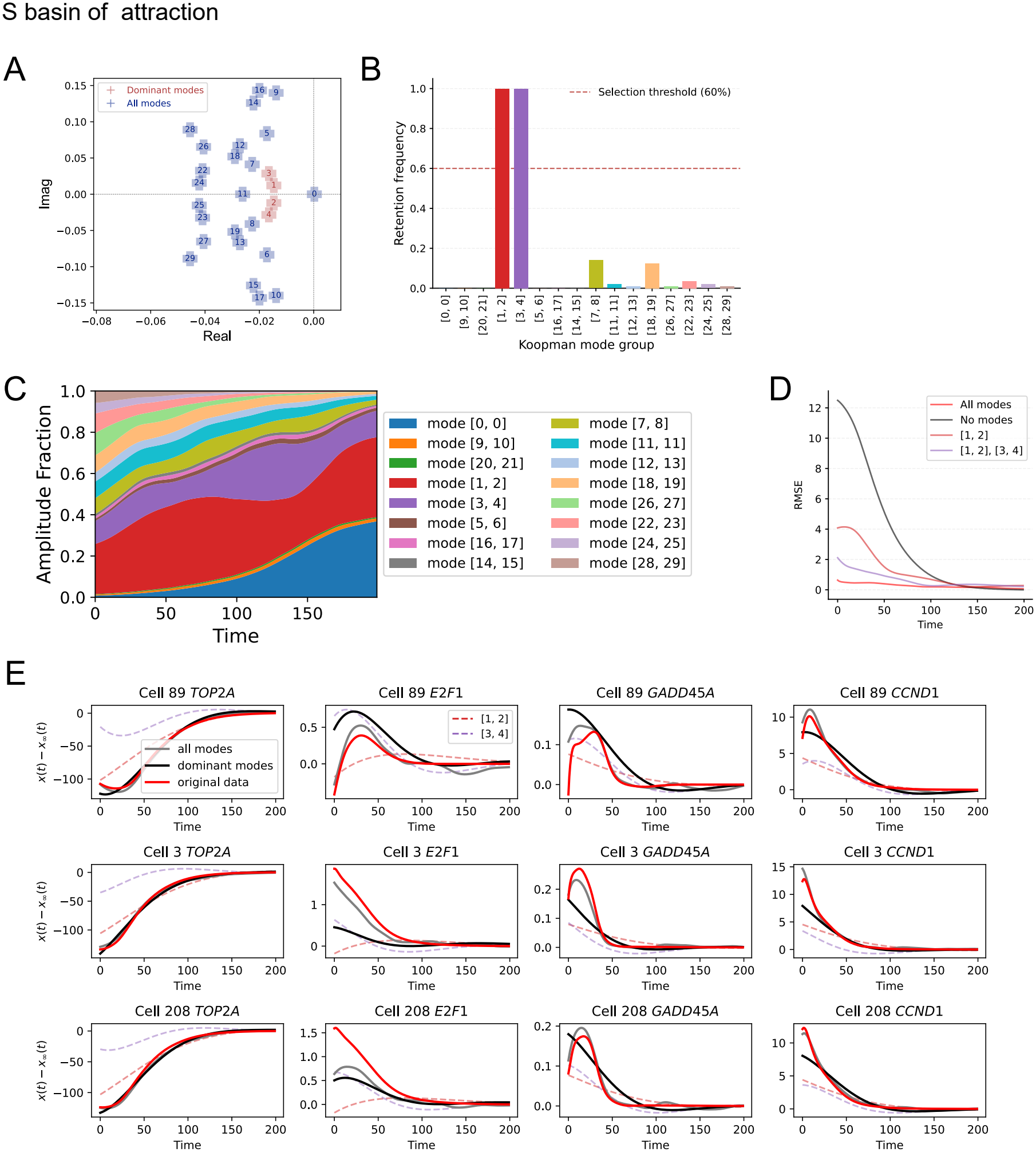
Dominant mode selection for the S basin. **(A–E)** The analyses and panel organization are the same as those described for the M/G1 basin in Fig S7, but are performed here for the S basin.

**Fig. S9.**
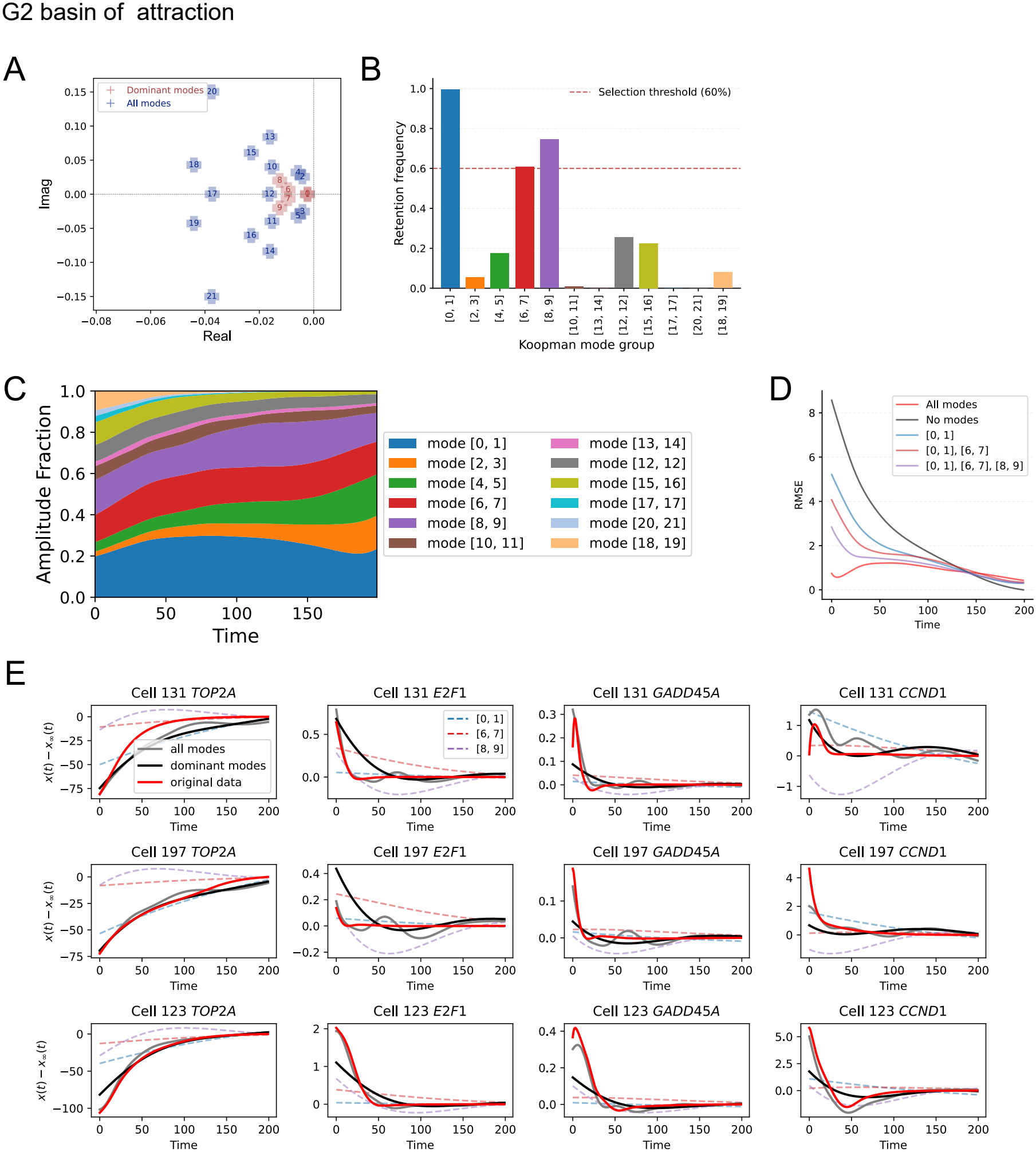
Dominant mode selection for the G2 basin. **(A–E)** The analyses and panel organization are the same as those described for the M/G1 basin in Fig S7, but are performed here for the G2 basin.

**Fig. S10.**
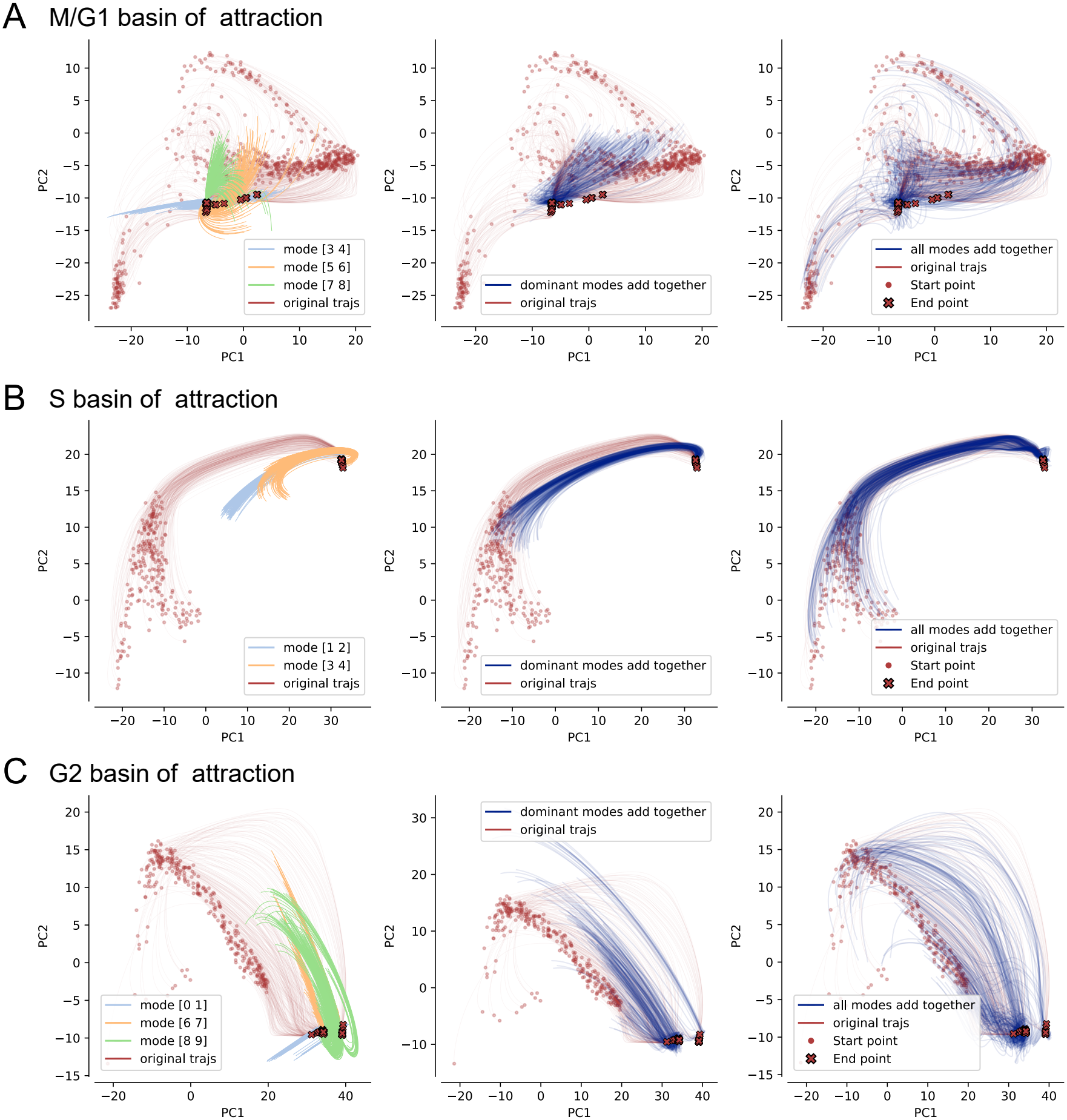
Projections of slow manifolds on two-component PCA space for each basin of attraction. Based on the greedy selection and amplitude analysis (Fig. S7, S8 and S9), we identified three dominant mode pairs in the M/G1 basin, two in the S basin and three in the G2 basin. **(A)**–**(C)** Slow manifolds projected onto PCA space for the M/G1 **(A)**, S **(B)** and G2 **(C)** basin. The left panels show the individual mode projected onto the PCA space, while the middle panels show the sum of the dominant modes, which aligns well with the original trajectories at later times. The right panels show the reconstructed trajectories using all the modes. Interactive HTML visualizations of the trajectories in three-dimensional PCA space are also provided.

**Fig. S11.**
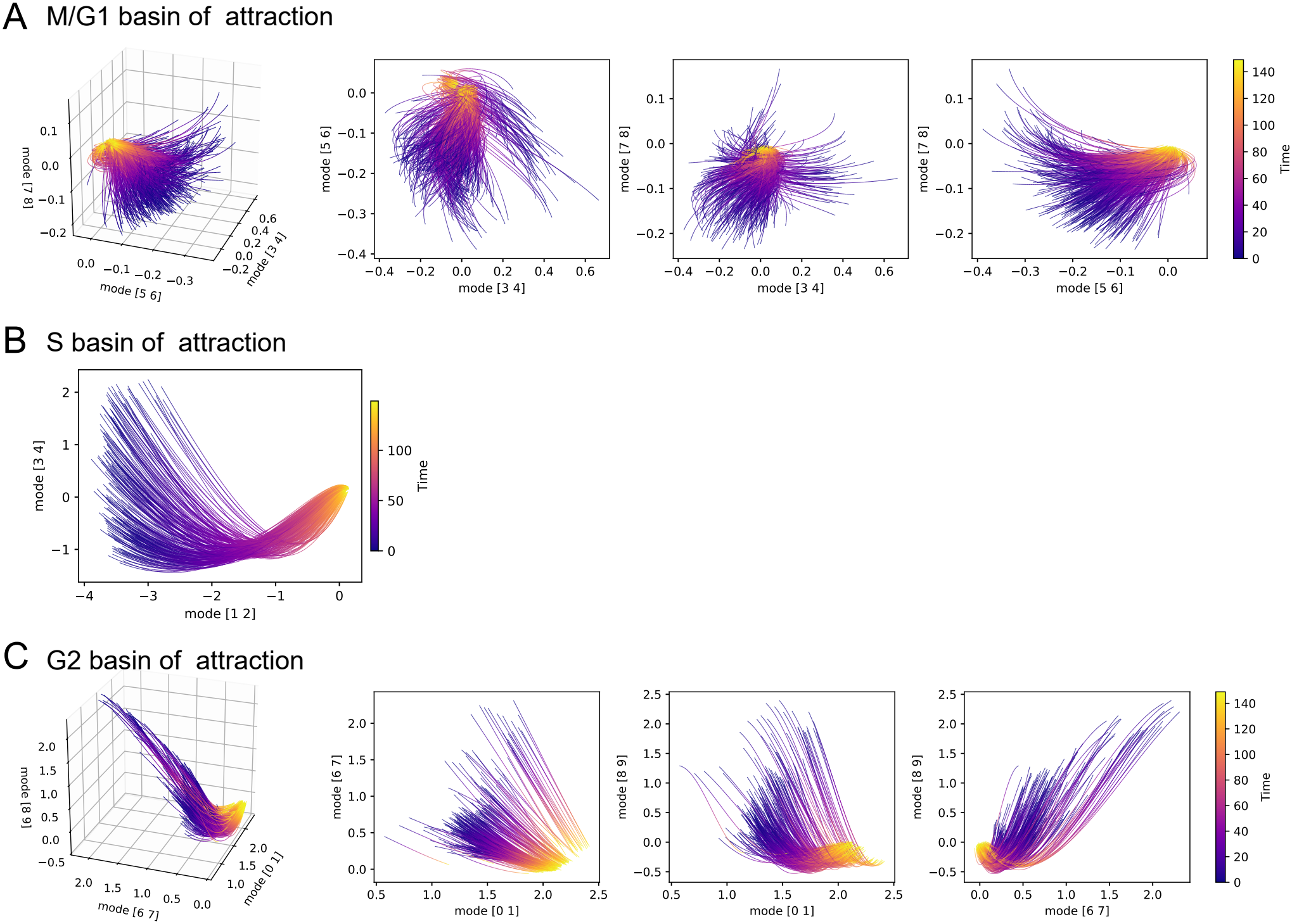
Trajectories represented in the coordinate spaces defined by the dominant Koopman modes for each basin of attraction, revealing the structure of the slow manifolds. In these representations, only the contributions from the Koopman eigenfunctions and eigenvalues are retained, while the Koopman mode coefficients for individual genes are set to unity. Each curve represents a trajectory starting from a different initial cell state. **(A)** M/G1 basin. The left panel shows a three-dimensional representation of the trajectory ensemble in the coordinate space defined by three dominant mode pairs, [3,4], [5,6], and [7,8]. The remaining three panels show the corresponding two-dimensional projections onto the three pairwise combinations of these mode coordinates. **(B)** S basin. The trajectory ensemble is shown in the two-dimensional coordinate space defined by two dominant mode pairs, [1,2] and [3,4]. **(C)** G2 basin. Similar to (A), but using three dominant mode pairs, [0,1], [6,7], and [8,9].

**Fig. S12.**
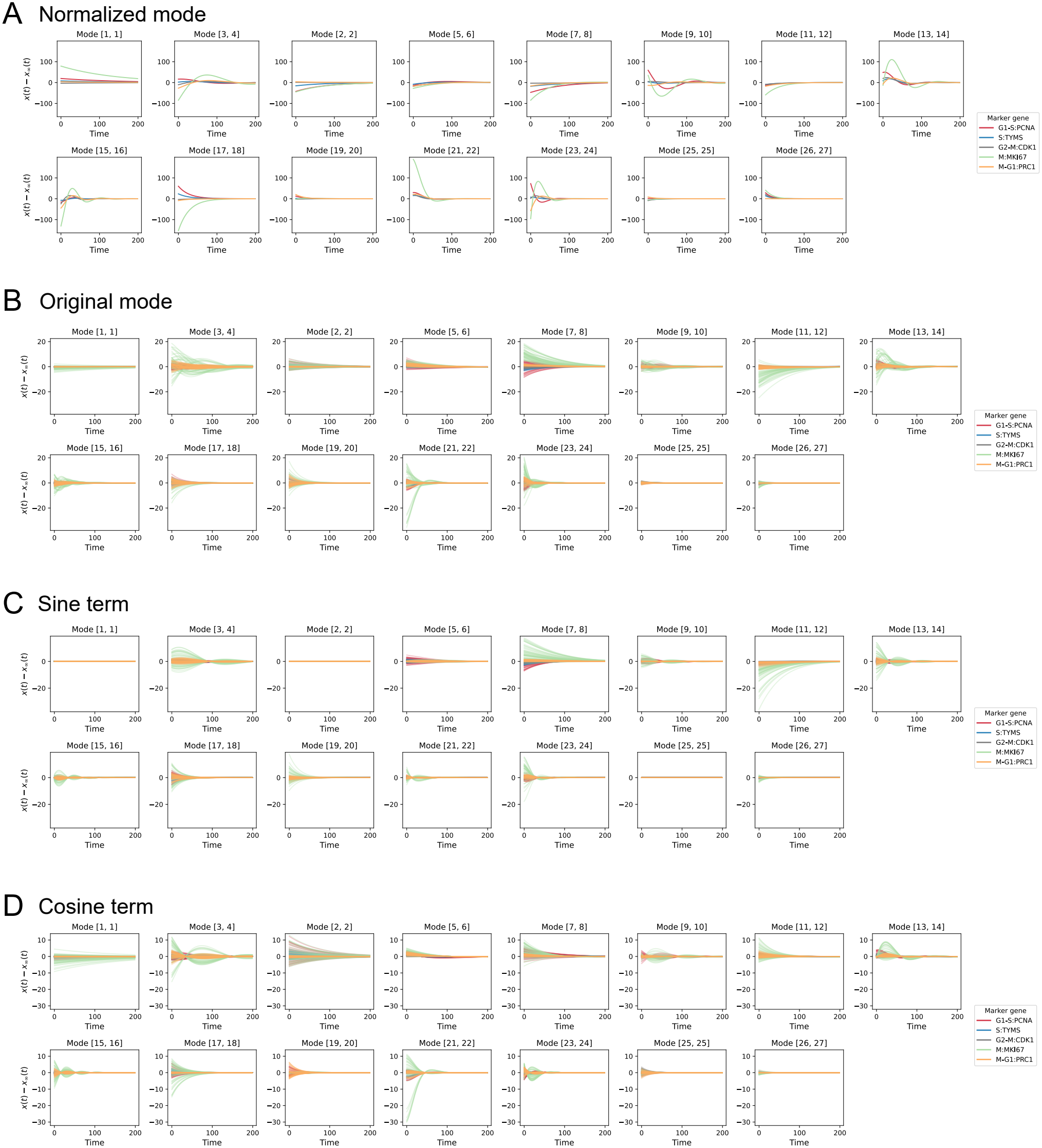
Collective modes in the M/G1 basin of attraction. Each subfigure shows the temporal evolution of a mode. Five marker genes were selected for each cell-cycle phase. Different colors represent different genes, while different lines correspond to trajectories initialized from different cells. We showed **(A)** the normalized modes without the initial condition dependent weighting prefactor (equivalently, *ϕ*_*k*_(x(0)) = 1), **(B)-(D)** The overall sum (see Eq. (16)) and the cosine and sine terms (see Eq. (17)) of the modes for the five selected marker genes, after removing the steady state contribution x_∞_(*t*) which in this case is time invariant fixed point.

**Fig. S13.**
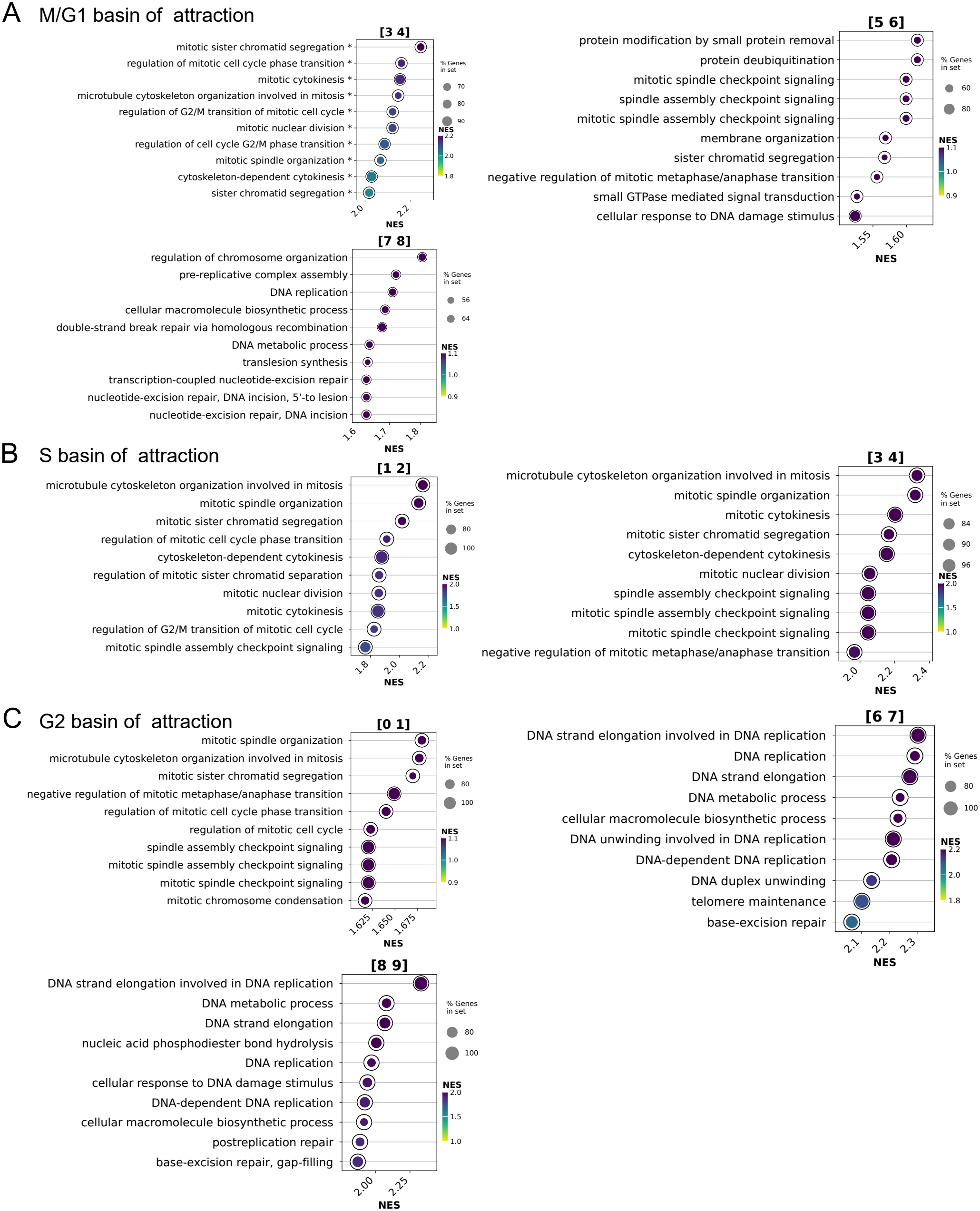
GO analysis results of the dominant modes for M/G1, S and G2 basins.

**Fig. S14.**
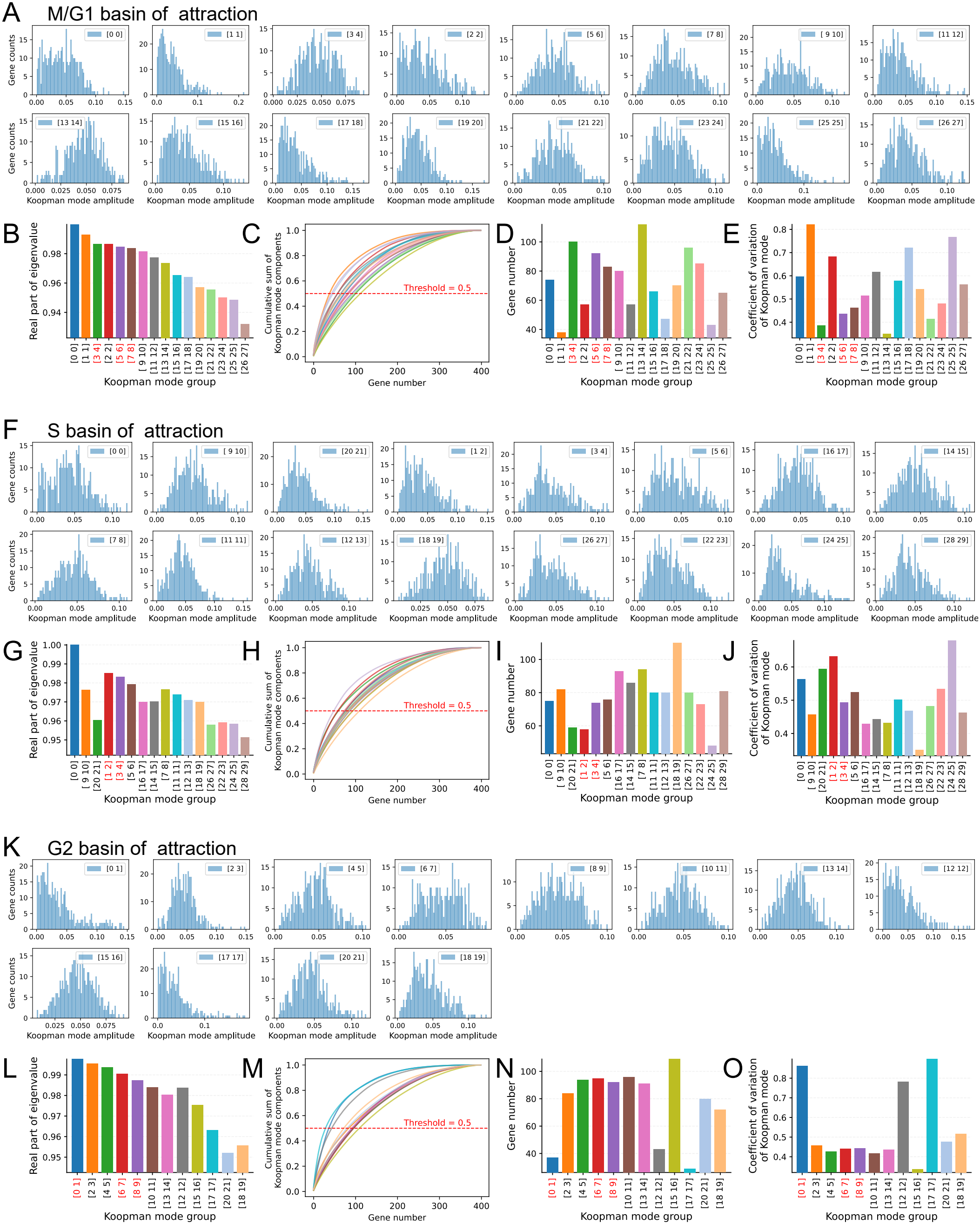
Mode localization across genes in the three basins of attraction. **(A), (F), (K)** Histograms of normalized Koopman mode component amplitudes across genes. **(B), (G), (L)** Real parts of the Koopman eigenvalues for each mode, indicating their characteristic timescales. **(C), (H), (M)** Cumulative fractions of total Koopman mode amplitude, with genes ranked in ascending order of their normalized component amplitudes. **(D), (I), (N)** Minimum numbers of genes required to account for 50% of the total mode amplitude for each mode. **(E), (J), (O)** Coefficients of variation (CV) of normalized Koopman mode component amplitudes across genes. Each set of five panels corresponds to one basin of attraction.

**Fig. S15.**
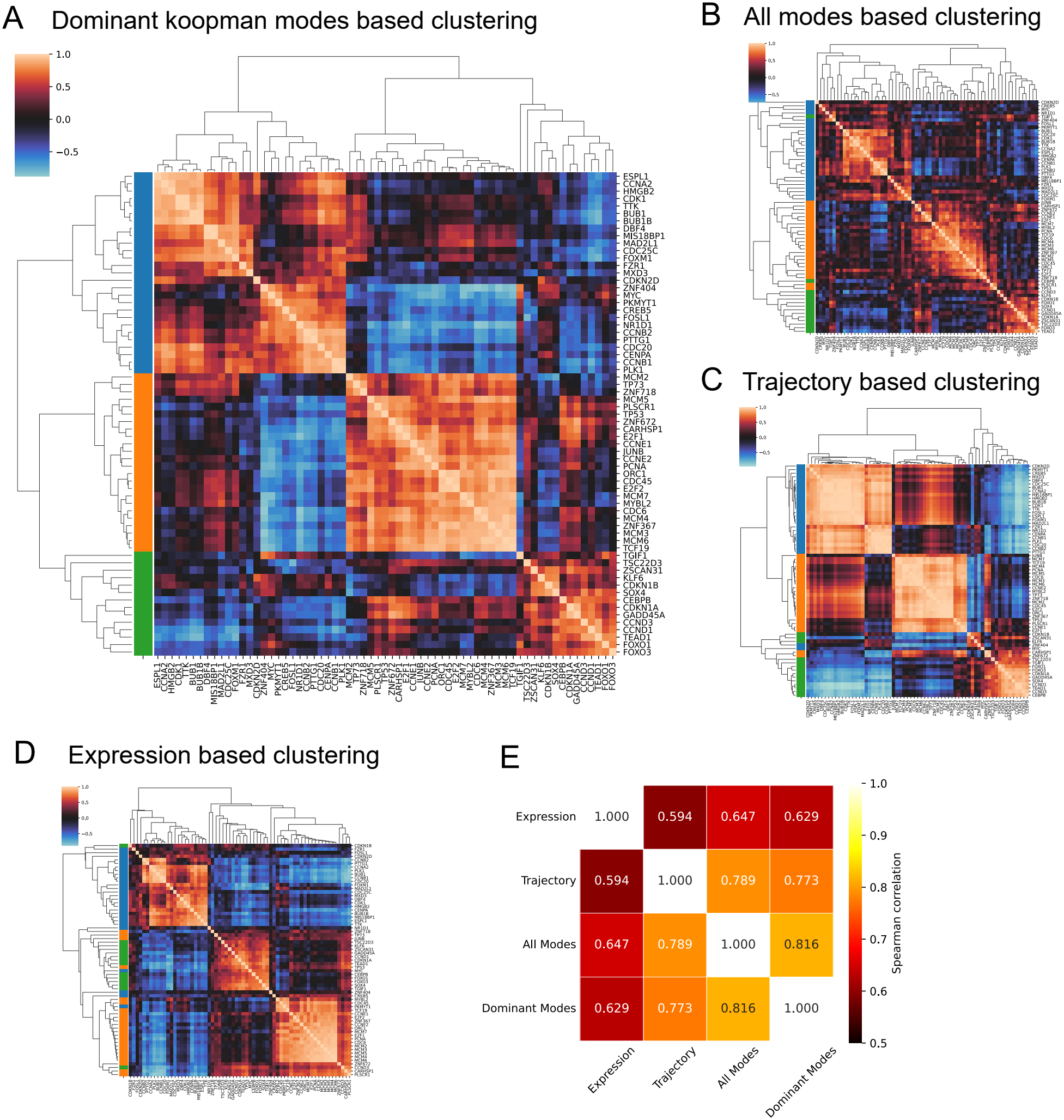
Gene clustering with different methods. (**(A)**–**(D)**) Clustering of core genes using different methods. (**(A)**–**(B)**) Clustering based on the normalized Hermitian inner product of the vectors constructed from Koopman modes (see main text and Methods), using either the dominant modes **(A)** or all modes **(B)**. The similar clustering patterns obtained in **(A)** and **(B)** suggest that the dominant modes retain sufficient information to coarse-grain the gene network. **(C)** Clustering based on trajectory information. For each gene, its trajectories were flattened into a single vector, and pairwise Pearson correlations were calculated between genes (see Methods). **(E)** Clustering based on the gene expression profile. The clustering obtained from Koopman modes is more similar to that obtained from trajectory-based clustering than to that obtained from the expression profile, suggesting that Koopman modes capture dynamic information beyond static expression similarity. **(E)** Quantification of the similarity between gene clustering results using Spearman correlation (see Methods).

**Fig. S16.**
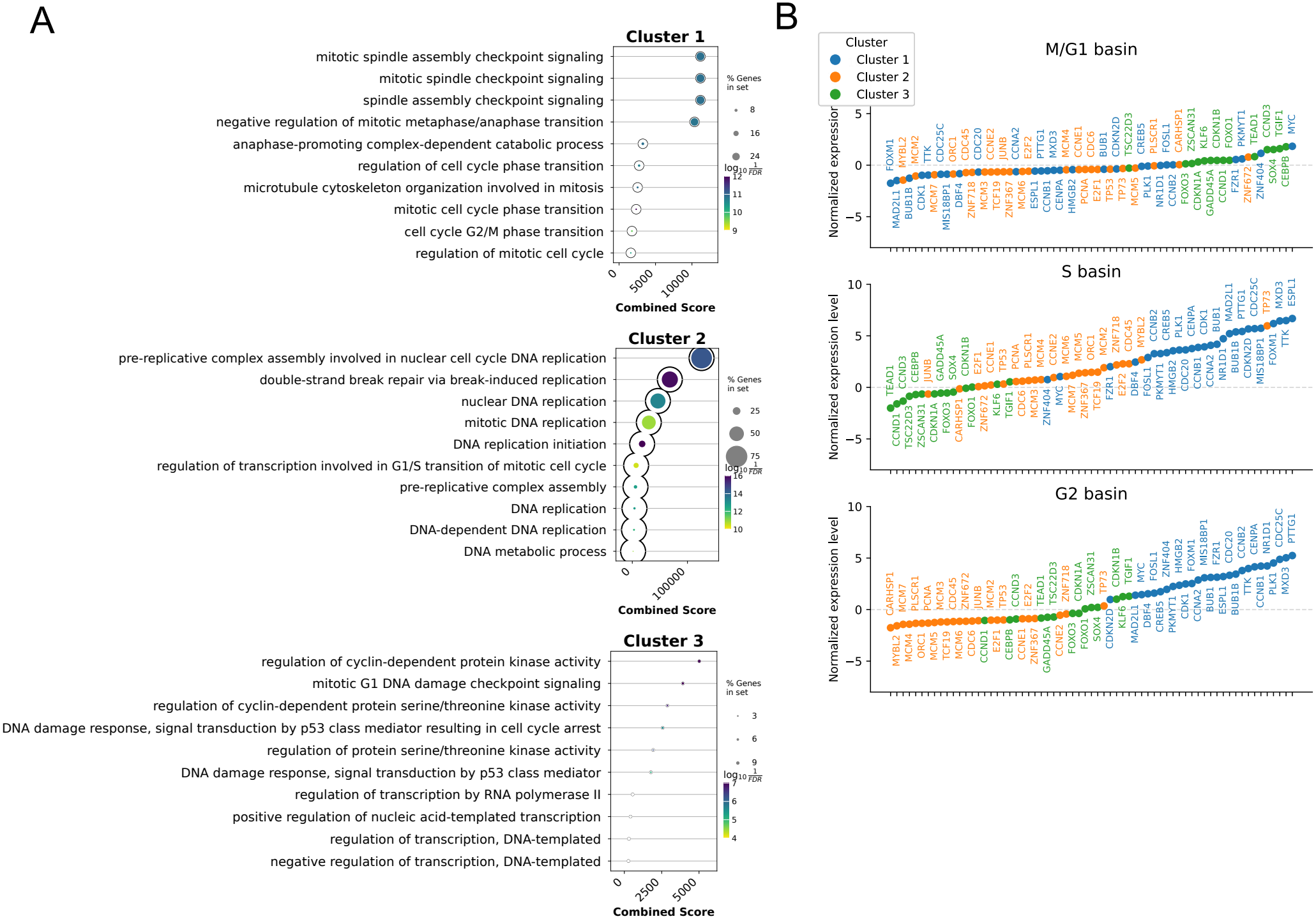
Biological functions of the three gene clusters. **(A)** GO enrichment analysis for each cluster. **(B)** Gene expression profiles of the fixed points in each basin of attraction. Expression values were normalized by subtracting the mean and dividing by the standard deviation of the corresponding gene expression levels across the initial states. Genes are ranked in ascending order of their normalized expression values, with different colors representing the three gene clusters.

## Footnotes

1 *F* : *ℳ → ℳ* to be more precise

2 Otherwise, the contribution of the corresponding term in Eq. (14) to the trajectory **x**(*t*) would be non-real.

3 If the system is time reversible (which happens in conservative or Hamiltonian systems), they form a group. But if the flow is forward only, it is a semigroup. Also, a group is just a special case of semigroup. So, overall, we can always consider the set of Koopman operators as a semigroup in general.

4 From the point of view of the statespace ***ℳ***, the dual space *ℱ* ^*\**^ consists of linear functionals that act on observable functions *ϕ*(**x**).

5 In other works, K can be thought of as the confinement of Koopman operator K within the operator space of *ℱ* _D_.

6 Feature *f* refers to *f* ^th^ state space coordinate *x*_*f*_ . In our case, this is the gene expression of the *f* ^th^ gene.

## Notes

### Competing Interest Statement

The authors have declared no competing interest.

### Funding

- National Institute of General Medical Sciences: GM148525
- U.S. National Science Foundation: 2325149

